# Ancient origins of alkaloid biosynthesis in medicinal clubmosses

**DOI:** 10.64898/2026.06.21.733354

**Authors:** Eric A. Fields, Colin Y. Kim, Ryan S. Nett

**Affiliations:** Department of Molecular and Cellular Biology, Harvard University; Cambridge, MA, 02138 USA

## Abstract

The plant kingdom is rich with medicinal natural products that are complex and difficult to access. Discovering how plants build these molecules can be challenging, especially for biosynthetic pathways that have unusual chemical transformations or require intricate coordination among cellular compartments. Here, we leveraged 300 million years of metabolic conservation to uncover how medicinal clubmosses organize extracellular and intracellular alkaloid biosynthesis to produce the Alzheimer’s disease therapeutic huperzine A (HupA). We reveal not only scaffold-forming enzymes that form key precursors to hundreds of clubmoss alkaloids, but also an essential transporter that connects metabolism across the plasma membrane to enable complete HupA biosynthesis. Our results demonstrate how ancient evolutionary conservation can be used to identify cryptic biosynthetic components and unexpected cellular organization in plant specialized metabolism.

## Main Text

Plants are renowned for producing specialized metabolites that both mediate biological interactions and serve as important medicines (*1*). However, for most plant natural products, we know relatively little about their biological function and the biosynthetic chemistry that generates their complex structures, including how metabolic reactions are organized and compartmentalized in specialized plant cells (*2*). This research gap is exemplified by the Lycopodium alkaloids, a class of nearly 600 molecules produced exclusively in clubmosses (Lycopodiaceae family) whose functions and biosynthesis are largely unknown (*3*). Many of these alkaloids have been conserved for more than 300 million years of clubmoss evolution (*4*), suggesting a critical function in the ecology of these plants. However, most Lycopodium alkaloids are inaccessible for biological study due to the difficulty in sourcing them from slowly growing clubmosses (*5*, *6*) and the complexity of their chemical synthesis (*7*). Nevertheless, the unique structures of Lycopodium alkaloids, often containing multiple fused rings with many stereocenters and unusual bond connectivity, highlight these molecules as a compelling system for identifying unprecedented biosynthetic processes and novel bioactivities.

The most well-studied Lycopodium alkaloid is huperzine A (HupA, **18**), which acts as a potent, reversible inhibitor of acetylcholinesterase (AChE) (*8*, *9*), a major target of both insecticides (*10*) and therapeutics that treat the symptoms of Alzheimer’s disease (*3*). This has prompted extensive studies on the chemistry and pharmacology of HupA (**18**), as well as its clinical use in China as an Alzheimer’s disease treatment (*3*, *11*). Despite the development of many chemical syntheses for HupA (**18**), it is still predominantly sourced from wild populations of *Huperzia serrata*, which has resulted in overharvesting of this species to the point of endangerment (*5*, *6*). Beyond HupA (**18**), the structural complexity and pharmacological promise of Lycopodium alkaloids motivated us to investigate their biosynthesis, with the expectation that this would not only enable metabolic engineering, but also reveal novel enzymatic and cellular mechanisms that plants use to build natural products.

Isotope tracer studies have demonstrated that lycopodine (**22**), the first Lycopodium alkaloid to be isolated (*12*), derives from *L*-lysine and malonyl-CoA (*13*), and subsequent studies further defined several early pathway intermediates (*14–18*). Guided by these results, recent work from our group and others identified a minimal set of enzymes that convert *L*-lysine and malonyl-CoA into a key tricyclic “phlegmarane” scaffold (**9**) that likely serves as a precursor to most Lycopodium alkaloids (**Fig 1A**) (*19–23*). Concurrently, we identified a minimal set of tailoring enzymes that convert the tetracyclic Lycopodium alkaloid precursor flabellidine (**12**) into HupA (**18**) (*22*, *23*). Moreover, flabellidine (**12**) has been identified in many distantly-related clubmosses (**Fig 1B**) (*4*, *24*, *25*), suggesting that it may be a key precursor to many tetracyclic Lycopodium alkaloids, including both HupA (**18**) and lycopodine (**22**). However, despite extensive testing of candidate enzymes predicted to convert the phlegmarane scaffold (**9**) into flabellidine (**12**), these final biosynthetic transformations had remained enigmatic, suggesting that there were unforeseen metabolic factors that we needed to consider.

**Fig. 1.**
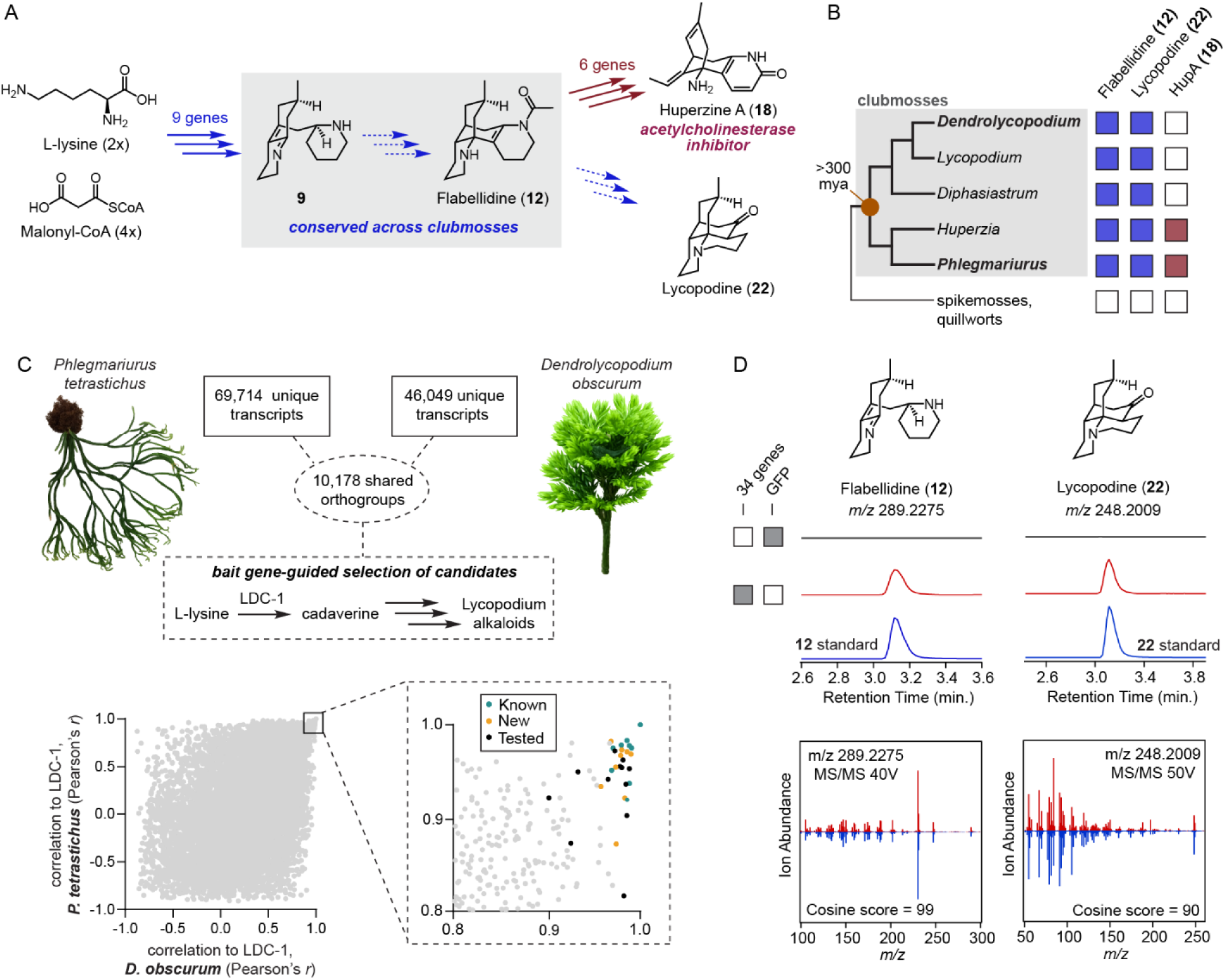
Evolution-guided transcriptomics identifies conserved genes for Lycopodium alkaloid scaffold formation. **(A)** Proposed biosynthetic pathway for conserved Lycopodium alkaloid scaffold formation. Dashed arrows indicate unknown genes. **(B)** Phylogenetic distribution of select Lycopodium alkaloids across major clubmoss lineages (crown age of ∼350 million years) (*26*). Colored squares indicate reported occurrence of each alkaloid class in a lineage with white indicating absence of an alkaloid. **(C)** Comparative transcriptomic strategy for identifying conserved biosynthetic genes from *Phlegmariurus tetrastichus* and *Dendrolycopodium obscurum*. Co-expression plots show the correlation of conserved transcripts to each species’ respective lysine decarboxylase (LDC-1) gene. Teal points indicate previously characterized pathway genes, gold points indicate newly identified biosynthetic genes, and black points indicate other experimentally tested candidates from the batch of 34. **(D)** Transient expression of a 34-gene candidate set in *Nicotiana benthamiana* produces flabellidine (**12**) and lycopodine (**22**), confirmed by matching LC-MS retention times and MS/MS (MS^2^) fragmentation patterns to chemical standards.

To identify the missing components of HupA (**18**) biosynthesis, we exploited the deep evolutionary conservation of Lycopodium alkaloid biosynthesis in clubmosses to guide an unbiased selection of gene candidates that could convert the tricyclic phlegmarane scaffold (**9**) into flabellidine (**12**). Using comparative transcriptomics of clubmoss species that diverged over 300 million years ago (*26*), we identified a minimal gene set that enabled the production of both flabellidine (**12,** lycodane scaffold) and lycopodine (**22**, lycopodane scaffold) in a single experiment, which allowed us to reconstitute a complete HupA (**18**) biosynthetic pathway in a model plant. This approach revealed an unexpected and essential role for a metabolite transporter that connects extracellular and intracellular compartments in specialized Lycopodium alkaloid biosynthetic cells. In parallel, our comparative approach uncovered multiple, unprecedented auxiliary proteins that substantially boost pathway yield and belong to protein families not previously associated with plant specialized metabolism. Thus, over 140 years after the initial isolation of Lycopodium alkaloids (*12*), we reveal how clubmosses use unusual chemistry, novel auxiliary proteins, and mixed cellular compartmentalization to biosynthesize HupA (**18**) and lycopodine (**22**). Our results emphasize that evolution-guided transcriptomics can rapidly uncover both specialized metabolic enzymes and unanticipated non-enzymatic factors that are important for alkaloid production to occur in plant cells.

## Results

### Evolution-guided analysis enables completion of HupA and lycopodine biosynthesis

Previous work in the HupA-producing clubmoss *Phlegmariurus tetrastichus* identified nine enzymatic genes that convert *L*-lysine and malonyl-CoA into a key tricyclic “phlegmarane” scaffold (**9**) (**Fig 1A**) (*22*, *23*). Additionally, this work demonstrated that the tetracyclic Lycopodium alkaloid flabellidine (**12**) is tailored by a series of Fe(II)/2-oxoglutarate dependent dioxygenases (2OGDs) and an alpha/beta hydrolase (ABH) to produce HupA (**18**). However, it remained unclear how the tricyclic phlegmarane scaffold of **9** was converted into flabellidine (**12**) or other tetracyclic Lycopodium alkaloids. Despite the strong co-expression of all previously identified biosynthetic genes (*22*, *23*), further mining of the *P. tetrastichus* RNA-seq data set failed to identify any enzymes that could move the pathway beyond **9**. With this in mind, we hypothesized that a comparative transcriptomic analysis of a second, distantly-related clubmoss species could help us to identify the enigmatic genes involved in tetracyclic Lycopodium alkaloid biosynthesis. For this, we chose to use *Dendrolycopodium obscurum* (*27*), which grows locally in the northeastern United States, but does not produce HupA (**18**). However, both *D. obscurum* (Lycopodioideae subfamily) and *P. tetrastichus* (Huperzioideae subfamily) produce flabellidine (**12**) despite being separated by over 300 million years of evolution (**Fig S1**) (*23*, *28*). Thus, we hypothesized that a core biosynthetic pathway connecting **9** to flabellidine (**12**) is likely conserved between *P. tetrastichus* and *D. obscurum* (**Fig 1A**). With this in mind, we reasoned that comparing gene conservation and co-expression between the two species would allow us to refine an unbiased list of candidate genes involved in core scaffold formation.

To test this hypothesis, we sought to profile gene expression in *D. obscurum* plants that were actively synthesizing Lycopodium alkaloids. For this, we collected *D. obscurum* from Harvard Forest (Petersham, MA) and used D_2_O labeling to confirm active Lycopodium alkaloid biosynthesis in the new growth of shoots (**Fig S2 & S3**) (*22*, *29*). Next, we used RNA-sequencing to measure gene expression in biosynthetically active and inactive tissues. Previous work in *P. tetrastichus* demonstrated that the Lycopodium alkaloid biosynthetic genes have high co-expression with lysine decarboxylase (LDC) (*23*), the first enzymatic gene in the pathway. The LDC ortholog from *D. obscurum* (*Do*LDC-1) showed enriched expression in new growth compared to older leaves and stems, which corroborated both our D_2_O labeling results and the pattern of flabellidine (**12**) accumulation in these tissues (**Fig S4**). Furthermore, putative orthologs of the previously identified biosynthetic genes that produce **9** showed strong co-expression with *Do*LDC, suggesting that Lycopodium alkaloid biosynthesis is tightly co-regulated in this species, much like in *P. tetrastichus* (**Fig S4**). Using Agrobacterium-mediated transient expression in *Nicotiana benthamiana*, we confirmed that the *D. obscurum* orthologs act together to produce **9**, consistent with the ancient conservation of core Lycopodium alkaloid biosynthesis across clubmosses (**Fig S4**).

We then used RNA-seq data sets from both clubmoss species to select a filtered list of gene candidates that met the following two criteria, regardless of their functional annotation: 1) they needed to be conserved (orthologous) among both *P. tetrastichus* and *D. obscurum*; 2) they had to strongly co-express with the previously identified biosynthetic genes in each species (see **Methods**). Using this comparative approach, only 34 gene candidates from *P. tetrastichus* fulfilled these criteria (**Fig S5)**. Importantly, this set of candidates included all currently known Lycopodium alkaloid biosynthetic genes that produce **9** (**Fig 1C**) (*23*). Testing this batch of 34 genes together via transient expression in *N. benthamiana* led to the production of flabellidine (**12**) (**Fig 1D**), indicating that the missing Lycopodium alkaloid biosynthetic genes were in this batch. Individual dropout experiments subsequently showed that four new genes were necessary for converting **9** into flabellidine (**12**): 1) a <u>m</u>edium chain <u>d</u>ehydrogenase/reductase (MDR-1) (*30*); 2) a <u>cy</u>tochrome <u>P</u>450 (CYP7032A12); 3) an <u>ac</u>etyltransferase (ACT-2); and most surprisingly, 4) a transporter belonging to the Nitrate transporter/Peptide transporter family (NPF8.1) (**Fig S6**) (*31*, *32*). Analysis of differentially accumulated mass features in these dropout experiments led us to propose a putative biosynthetic hypothesis (**Fig S6**). First, MDR-1 and NPF8.1 act in tandem to convert **9** into the reduced product **10**. Second, **10** is oxidized by CYP7032A12 to produce a mixture of isomers (**11a**/**11b**). Finally, ACT-2 *N*-acetylates **11b** to produce flabellidine (**12**) (**Fig 2A**).

**Fig. 2.**
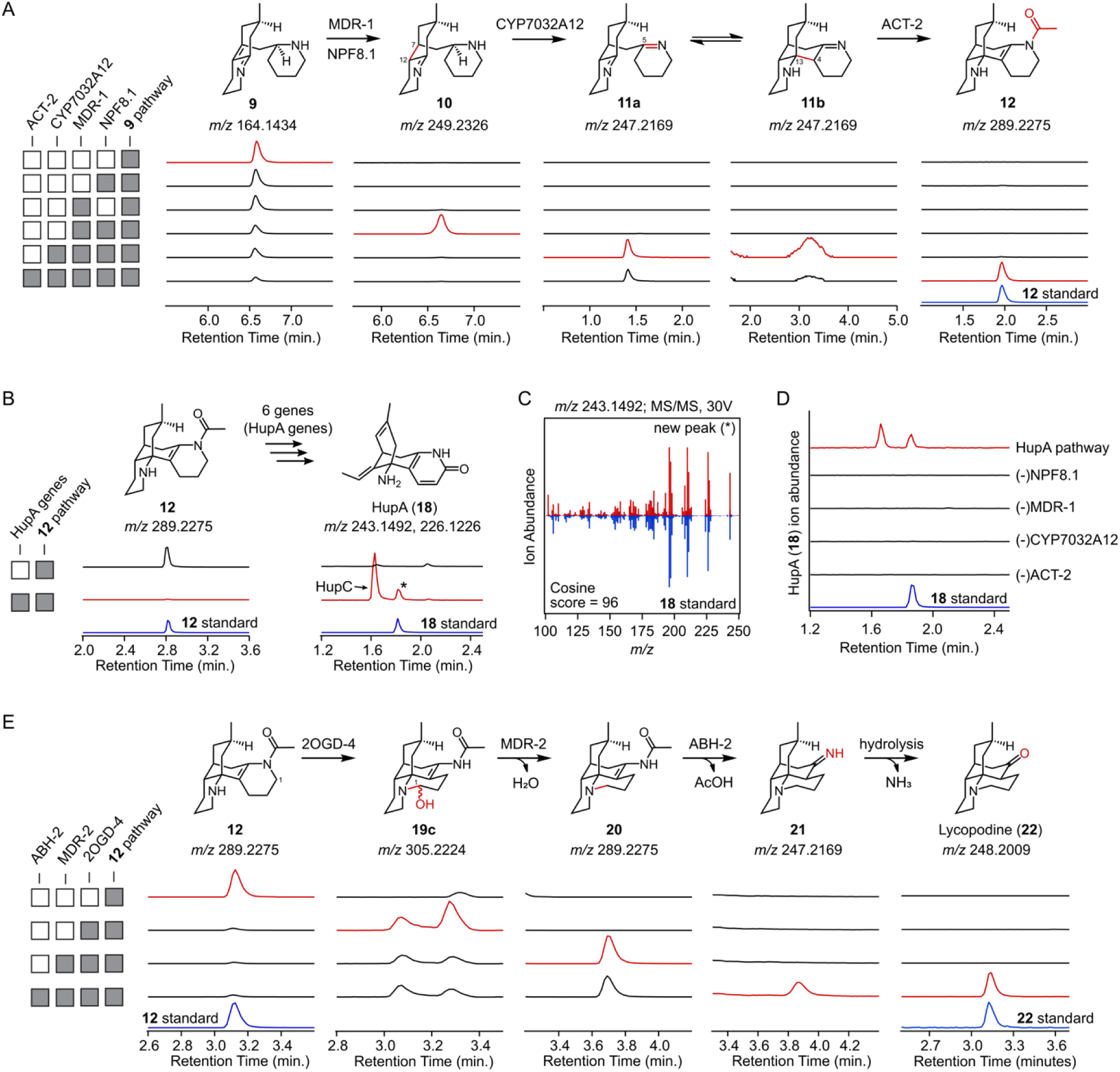
Reconstitution of the complete pathways to huperzine A (18)and lycopodine (22). **(A)** Stepwise reconstruction of flabellidine (**12**) biosynthesis from the tricyclic phlegmarane scaffold **9** in *N. benthamiana*. Chromatograms show extracted ion chromatograms for pathway intermediates. **(B)** Reconstitution of complete HupA (**18**) biosynthesis from flabellidine (**12**) using six previously identified HupA-tailoring enzymes. Extracted ion chromatograms show production of **18** in *N. benthamiana* relative to an authentic standard. **(C)** MS/MS (MS^2^) fragmentation spectrum of the newly detected peak (*m/z* 243.1492) from the reconstructed **18** pathway (red) compared to a standard (blue). **(D)** Dropout analysis of the reconstructed **18** pathway. **(E)** Stepwise reconstruction of lycopodine (**22**) biosynthesis from flabellidine (**12**). Extracted ion chromatograms from pathway dropout experiments are shown below each metabolite. For all panels, gray squares indicate inclusion of pathway genes and white squares indicate omission of genes in each transient-expression condition.

We next systematically tested different combinations of these four newly discovered genes on top of our engineered pathway to **9**. First, we found that **9** was only consumed with the simultaneous expression of both the reductase MDR-1 and NPF8.1, which led to production of a molecule (**10**) consistent with a double bond reduction ([M+H]^+^ = *m/z* 249.2326) (**Fig 2A**). We further demonstrated that co-infiltration of **9** into leaves expressing only MDR-1 and NPF8.1 yielded **10**, and that the CAL-1/CAL-2 enzymes, which directly produce **9** (**Fig S7**), were required for **10** biosynthesis in *N. benthamiana*. Of the two double bonds in **9**, we hypothesized **10** would retain the imine rather than the alkene to enable downstream carbon-carbon coupling. Attempts to isolate **10** from our *N. benthamiana* transient expression system for structural elucidation were unsuccessful, as **10** degraded during purification. To provide additional structural insight into **10**, we derivatized this molecule via NaBD_4_ reduction, which supported the presence of an imine in **10** and a shared phlegmarane scaffold with **9**, consistent with MDR-1 catalyzing a reduction on the alkene in **9** to produce the proposed structure of **10** (**Fig S8**).

Subsequent addition of CYP7032A12 to the pathway led to a depletion of **10** and the production of two putative isomers (**11a** and **11b**) with exact masses that corresponded to a desaturation of **10** ([M+H]^+^ = *m/z* 247.2169) (**Fig 2A**). To enable lycodane scaffold formation, we predicted that this desaturation would occur between C5 and Nα of **10** to yield an imine. The desaturation at this position would produce a molecule with two piperideine rings that could undergo a spontaneous Mannich-like condensation (mimicking 1-piperideine dimerization (*33*)) to produce the lycodane scaffold. Indeed, **11a** and **11b** appeared to be in equilibrium, as attempts to purify each peak led to a time-dependent increase of the other (**Fig S9**). To investigate the structures of these mass features, we first performed MS/MS (MS^2^) fragmentation and found that: 1) the fragmentation of **11a** is nearly identical to that of **10**; and 2) the MS^2^ spectrum of **11b** is highly similar to that of flabellidine (**12**) (**Fig S9**). This is consistent with **11a** having a ring-opened, phlegmarane diimine scaffold, similar to **10**, while **11b** likely possesses a ring-closed lycodane scaffold like flabellidine (**12**), bearing a single imine. Consistent with this, application of NaBD_4_to an extract containing predominantly **11b** produced a singly reduced mass feature ([M+D]^+^ = *m/z* 250.2389), but no masses consistent with a double reduction, further supporting that **11b** possesses the tetracyclic lycodane scaffold (**Fig S10**). As a final level of structural support, hydrolysis of flabellidine (**12**) in strong base to remove its acetyl group led to production of **11b** (**Fig S11**).

We next found that addition of ACT-2 to the transiently expressed pathway to **11a/11b** led to consumption of **11b** and concurrent production of flabellidine (**12**) (**Fig 2A**). With the pathway to flabellidine (**12**) resolved, we next tested if we could successfully engineer the complete pathway to HupA (**18**) through transient expression in *N. benthamiana*. Indeed, the addition of the six previously identified tailoring enzymes (2OGD-4, 2OGD-5, ABH-1, 2OGD-3, 2OGD-1, and 2OGD-2) on top of the pathway to flabellidine (**12**) resulted in the complete reconstitution of HupA (**18**) biosynthesis *in planta* (**Fig 2B-D & S12**) (*22*, *23*). Thus, by identifying the final genes needed for flabellidine (**12**) biosynthesis, we have confirmed a minimal set of 18 genes for the heterologous production of HupA (**18**) in a model plant.

In addition to identifying the necessary genes for flabellidine (**12**) biosynthesis, our 34 gene batch test also led to the production of lycopodine (**22**), which serves as the scaffold precursor to over 170 downstream Lycopodium alkaloids (**Fig 1D & S13**) (*3*). Dropout experiments from our batch of 34 genes demonstrated that three genes are involved in converting flabellidine (**12**) into lycopodine (**22**) (**Fig S14**). This included a previously identified 2OGD enzyme (2OGD-4) that oxidizes flabellidine (**12**) (*23*), a second MDR (MDR-2), and an alpha/beta hydrolase (ABH-2). Based on accumulation of mass features in the dropout experiments and using chemical logic, we hypothesized the following series of biosynthetic transformations: 1) 2OGD-4 hydroxylates flabellidine (**12**) to produce a hemiaminal (**19a**) that can rearrange to form a bond between C1 and Nβ (see **Supplementary Scheme** for atom numbering), thereby generating an intermediate with the lycopodane scaffold (**19c**); 2) C1 of **19c** is reduced by MDR-2 to produce flabelline (**20**); 3) the *N*-acetyl group of **20** is hydrolyzed to yield the imine-containing intermediate **21**, which can then undergo hydrolysis to produce lycopodine (**22**).

To assess our biosynthetic hypothesis, we expressed these genes with the flabellidine (**12**) pathway to reconstruct lycopodine (**22**) biosynthesis step-by-step. The addition of 2OGD-4 to this pathway produced multiple new mass features consistent with the addition of a hydroxyl group [M+H]^+^ = (*m/z* 305.2224) (**19a**/**19b**/**19c**), which was previously observed for 2OGD-4 activity on flabellidine (**12**) as a substrate (**Fig 2E & S15**). Attempts to purify these metabolites with HPLC revealed that they are in equilibrium and their MS^2^ fragmentation suggested that they may represent spontaneously rearranged products that can form after hydroxylation of flabellidine (**12**) at C1 (**Fig S15**). Adding MDR-2 on top of the pathway to **19** produced a new mass feature (**20**) with the same exact mass as flabellidine (**12**) ([M+H] = *m/z* 289.2275), but with a different retention time (**Fig 2E**). We purified the new molecule from *N. benthamiana* leaves transiently expressing this pathway and used NMR analysis to confirm that it is flabelline (**20**), which possesses the lycopodane scaffold (**Fig S16-S22**). Finally, adding ABH-2 to the transiently expressed pathway to flabelline (**20**) produced both a new molecule (**21**) consistent with an *N*-deacetylation ([M+H]^+^ = *m/z* 247.2169) and lycopodine (**22**). Thus, we predict that ABH-2 hydrolyzes the *N*-acetyl group of flabelline (**20**) to produce **21**, which would contain a primary imine that can undergo hydrolysis to yield lycopodine (**22**) as a final product (**Fig 2E**).

In addition to establishing the biosynthetic logic for lycopodane scaffold formation, we found that 2OGD-4 activity serves as a branch-point in biosynthesis. Specifically, a single hydroxylation at C1 of flabellidine (**12**) by 2OGD-4 yields an intermediate (**19a**) that can rearrange to produce an imine-containing molecule (**19c**) with the lycopodane scaffold. Alternatively, 2OGD-4 can catalyze an additional oxidation at the C1 carbon of flabellidine (**12**) to produce an imide, which allows for the lycodane scaffold configuration to be “locked” in place for subsequent conversion into HupA (**18**) (*23*). Interestingly, we find that the 2OGD-4 ortholog from *P. tetrastichus* readily performs both a single and double oxidation at C1, while the ortholog from *D. obscurum* produces almost exclusively the singly oxidized product (**Fig S23**). These differing activities mirror the major products found in each clubmoss species, with *P. tetrastichus* producing high levels of HupA (**18**, lycodane scaffold) (*22*), and *D. obscurum* accumulating lycopodine (**22**, lycopodane scaffold) (*28*). Thus, 2OGD-4 activity seems to act as a putative means of control between lycodane and lycopodane scaffold formation.

### Cellular organization in Lycopodium alkaloid biosynthesis

The key result that allowed us to complete HupA (**18**) biosynthesis was the discovery that conversion of **9** into **10** required both a reductase enzyme (MDR-1) and a transporter (NPF8.1) during pathway reconstruction in *N. benthamiana*. To decouple the functions of NPF8.1 and MDR-1, we first showed that MDR-1 converts **9** into **10** independently of NPF8.1 *in vitro* using whole cell lysates from *N. benthamiana* (**Fig 3A & S24**). This suggested that NPF8.1 was likely crucial for transport rather than directly contributing to MDR-1 enzyme activity. Previously, we determined that production of **9** occurs in the extracellular space of the plant cell (apoplast) through the activity of secreted CAL enzymes (CAL-1, CAL-2, and CAL-3) (*23*). We hypothesized that **9**, which would be predominantly charged at the low pH of the apoplast (pH 5-6), is too polar to readily diffuse across membranes and that NPF8.1 transports **9** to the cytosol, where MDR-1 is predicted to localize (*34*). We attempted to express NPF8.1 in *Xenopus laevis* oocytes and yeast cells to directly measure metabolite transport (*35*), but we did not observe functional protein production in either system. As an alternative, we investigated the effect of adding NPF8.1 onto the pathway to **9** in *N. benthamiana* transient expression. Of the six pathway intermediates detectable in this experiment, we only found that **9** decreased significantly when NPF8.1 was co-expressed (**Fig S25**). This depletion co-occurred with a dramatic increase in an unknown mass feature ([M+H]^+^ = *m/z* 275.2118) that appears to be **9**-derived (**Fig S25**). Though indirect, this result supports that NPF8.1 likely transports **9**, putatively from the apoplast to the cytosol, where native *N. benthamiana* metabolism converts this molecule into off-target products.

**Fig. 3.**
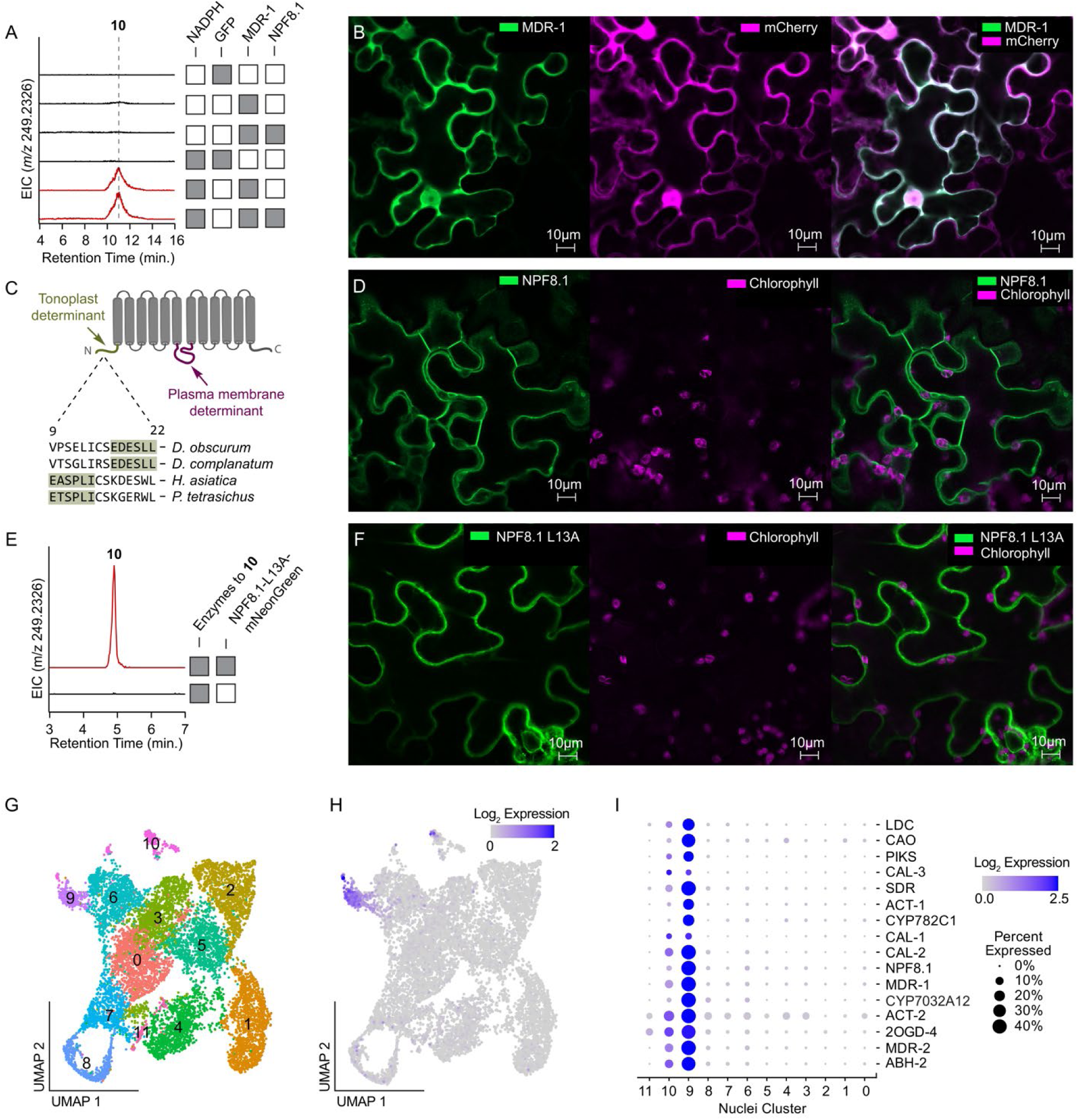
An essential transporter connects extracellular scaffold-forming reactions to intracellular Lycopodium alkaloid biosynthesis. **(A)** In vitro assays with *N. benthamiana* lysates demonstrate that MDR-1 converts the phlegmarane scaffold (**9**) to a reduced intermediate (**10**) independently of NPF8.1. Extracted ion chromatograms for **10** are shown across lysate and cofactor conditions. **(B)** Confocal microscopy of transiently expressed MDR-1–mNeonGreen in *N. benthamiana* epidermal cells. Green (left panel) indicates MDR-1, magenta (middle panel) indicates the mCherry cytosolic marker, and white in the merged image (right panel) indicates colocalization. **(C)** Cartoon representation of NPF8.1 membrane topology and putative localization signals. Aligned N-terminal sequences from representative clubmoss NPF8.1 orthologs highlight conservation of a tonoplast localization motif. **(D)** Confocal microscopy of NPF8.1–mNeonGreen in *N. benthamiana* epidermal cells (green) in comparison to chloroplasts (magenta). **(E)** Mutation of the conserved N-terminal tonoplast localization motif in NPF8.1 to alanine (L13A) preserves the ability of NPF8.1 to enable **10** production in *N. benthamiana* during pathway reconstruction, shown here via LC-MS chromatograms. **(F)** Confocal microscopy of NPF8.1-L13A–mNeonGreen localization. Green indicates NPF8.1-L13A and magenta indicates chlorophyll autofluorescence. **(G)** UMAP clustering of single-nucleus RNA-seq profiles from biosynthetically active *D. obscurum* tissue. **(H)** Expression of Lycopodium alkaloid biosynthetic genes in *D. obscurum* nuclei. **(I)** Dot plot showing cluster-specific expression of Lycopodium alkaloid biosynthetic genes.

We next wanted to validate the localization of both NPF8.1 and MDR-1. To assess this, we generated functional fluorophore-tagged MDR-1 and NPF8.1 constructs and used confocal microscopy to assess their localization in plant cells during transient expression in *N. benthamiana* (**Fig S26**). This analysis confirmed that MDR-1 localizes to the cytosol (**Fig 3B & S27**) and demonstrated that NPF8.1 co-localizes with both plasma membrane and tonoplast (vacuole) membrane markers (**Fig S28 & S29**) (*36*). Moreover, we observed signal for NPF8.1 on both sides of chloroplasts, further supporting its localization to the plasma membrane and tonoplast membrane (**Fig 3D**). Because **9** is synthesized in the apoplast, we suspected that the plasma membrane was the key location for NPF8.1 function within Lycopodium alkaloid biosynthetic pathway reconstitution. To test this, we considered strategies to “force” this protein exclusively to the plasma membrane. A specific N-terminal motif causes tonoplast localization for plant NPF family transporters (*37*), and we found that this motif is present in the N-terminus of NPF8.1 orthologs (**Fig 3C**). Mutating this sequence in *Pt*NPF8.1 (L13A) led to localization on the plasma membrane and no detectable signal on the tonoplast (**Fig 3F, S30**). Importantly, this plasma membrane-localized mutant still facilitated Lycopodium alkaloid biosynthesis (**Fig 3E & S31**), supporting the hypothesis that NPF8.1 transports **9** from the apoplast into the cytosol to enable MDR-1 activity.

The mixture of intracellular and extracellular biosynthetic compartments and the crucial role of metabolite transport led us to assess how Lycopodium alkaloid production was organized among clubmoss cells. In particular, we sought to understand if complete biosynthesis occurs within individual clubmoss cells, or if upstream (pre-scaffold formation) and downstream (post-scaffold formation) biosynthetic modules were compartmentalized among multiple cell types, which has been observed for other complex alkaloid pathways in plants (*2*, *38*). To assess how expression of each pathway gene was organized among clubmoss cells, we isolated nuclei from biosynthetically-active new growth tissue of *D. obscurum* and performed single nuclei RNA-sequencing (snRNA-seq). In this experiment, we found that expression of Lycopodium alkaloid biosynthetic genes was enriched in a small, distinct cluster of nuclei (∼1% of total nuclei), with no obvious sub-clustering of upstream and downstream biosynthetic modules into their own clusters (**Fig 3G-I & S32**). Indeed, multiple individual nuclei within this cluster express genes from the early and late portions of lycopodine (**22**) biosynthesis, indicating that individual cells have the capacity to carry out a complete pathway autonomously, and that pathway modules are not separated among different cell types (**Fig S33**). Thus, we propose a model in which each biosynthetic cell can be self-autonomous for Lycopodium alkaloid biosynthesis, with the apoplast serving as a separate compartment for scaffold formation.

### Conserved auxiliary proteins from clubmosses enhance Lycopodium alkaloid yields

By considering the evolutionary conservation of core Lycopodium alkaloid biosynthesis across clubmosses, we have uncovered the necessary and sufficient set of genes to reconstitute complete pathways for both HupA (**18**) and lycopodine (**22**) (**Fig 4A**). Additionally, while investigating the conserved batch of 34 genes that contained the missing biosynthetic components, we noticed that four additional genes boosted the yield of Lycopodium alkaloid biosynthesis: 1) an <u>A</u>TP <u>b</u>inding <u>c</u>assette family transporter (ABCC-1) (*39*); 2) a fourth, unique carbonic anhydrase-like protein (CAL-4); 3) a protein from the antibiotic biosynthesis monooxygenase family (Lycopodium alkaloid biosynthesis factor 1, or LABF-1) (*40*); and 4) a protein that has no predicted Pfam annotation (Lycopodium alkaloid biosynthesis factor 2, or LABF-2). Rather than produce new biosynthetic products, these genes affected the accumulation of established pathway intermediates, suggesting that they may have an auxiliary role in facilitating biosynthesis. Indeed, we found that each of these genes individually increased the accumulation of flabellidine (**12**) when they were co-expressed with the minimal set of biosynthetic genes needed to make this product (**Fig 4B**). Further, we found that the combination of all the auxiliary genes together with the previously reported CAL-3 led to a 24-fold increase in flabellidine (**12**) and an 8-fold increase in HupA (**18**) when co-expressed with the minimal HupA pathway in *N. benthamiana* (**Fig 4C**).

**Fig. 4.**
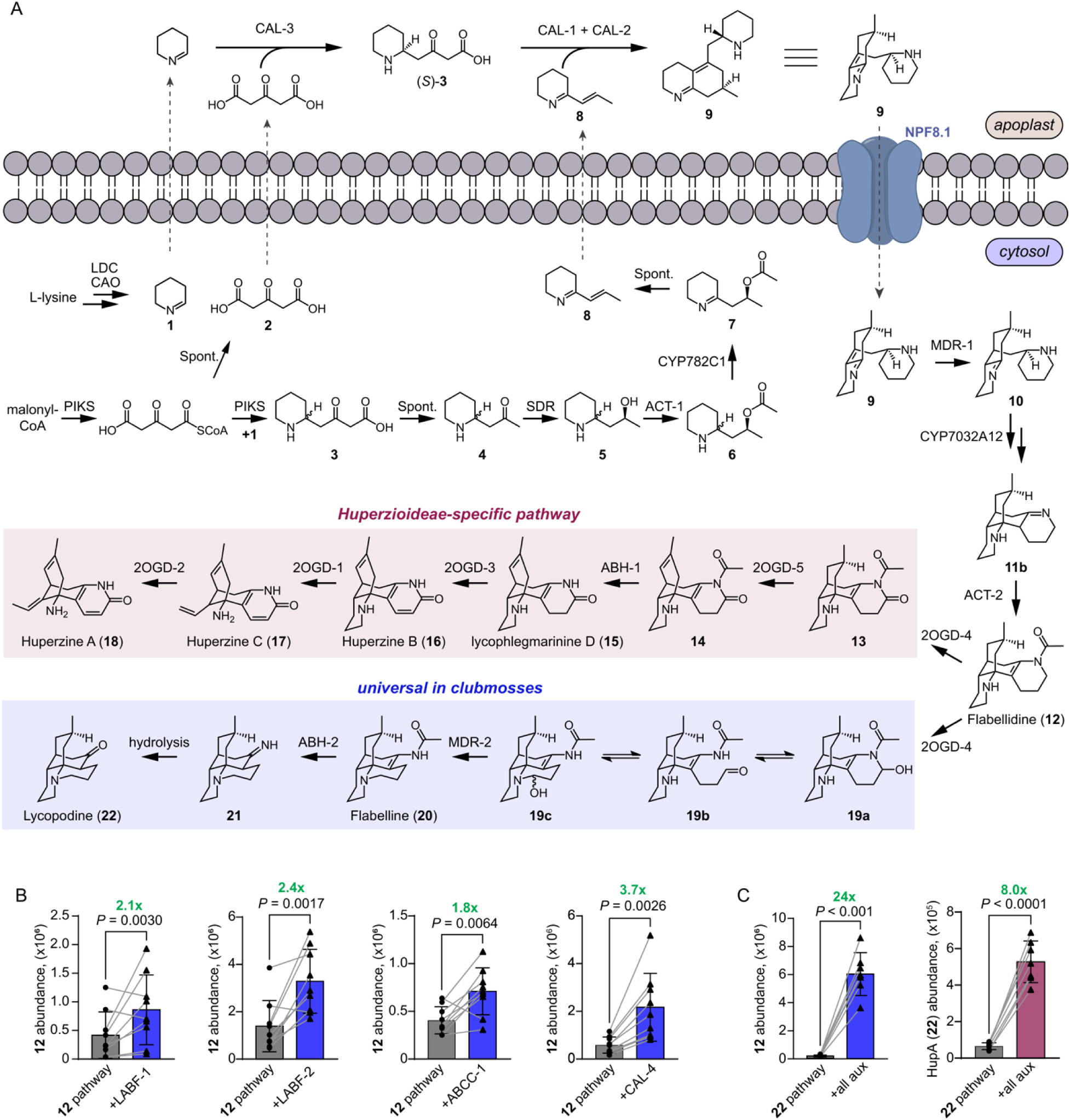
The complete biosynthesis of Lycopodium alkaloids. **(A)** Summary of enzyme-catalyzed transformations that convert *L*-lysine and malonyl-CoA to HupA (**18**) and lycopodine (**22**). Dashed arrows indicate proposed metabolite movement between the apoplast and cytosol. **(B)** Individual auxiliary genes increase flabellidine (**12**) accumulation during pathway reconstitution in *N. benthamiana*. Bars show mean ion abundance, error bars indicate standard deviation, and gray lines connect paired leaf infiltrations. Fold changes and *P* values are indicated above each comparison. **(C)** Combined expression of auxiliary genes increases the production of flabellidine (**12**) and huperzine A (**18**) during biosynthetic pathway reconstitution in *N. benthamiana*. Bars show mean ion abundance, error bars indicate standard deviation, and gray lines connect paired biological comparisons. Fold changes and *P* values are indicated above each comparison.

Each auxiliary gene is expressed in the same cell cluster as the essential Lycopodium alkaloid biosynthetic genes in *D. obscurum* (**Fig S34**), supporting a dedicated role in this specialized metabolic pathway. While the precise functions of the auxiliary proteins will be a focus of future in-depth biochemical studies, we could obtain preliminary insight by observing their effects on intermediate accumulation during Lycopodium alkaloid pathway reconstitution in *N. benthamian*a. This analysis suggested that the transporter ABCC-1 likely facilitates movement of intermediates upstream of phlegmarane scaffold (**9**) formation (**Fig S35**), further emphasizing the importance of understanding the cellular compartmentalization of alkaloid biosynthesis (*2*, *35*, *41*). Similarly, CAL-4 appears to act early in the pathway, and its predicted apoplast secretion signal suggests that it may contribute to the CAL-mediated extracellular reactions (**Fig S36**) (*23*). Finally, LABF-1 and LABF-2 seem to be involved in the formation of phlegmarane and lycodane scaffolds, respectively (**Fig S37 & S38**). Although not strictly essential, the conservation of these genes over 300 million years of clubmoss evolution highlights the importance of auxiliary factors in facilitating the efficient biosynthesis of Lycopodium alkaloids, and plant natural products more broadly (*42*).

## Discussion

Despite major advances in our ability to characterize plant biosynthetic pathways (*43*), many critical biosynthetic features – including the genes and metabolic organization needed to synthesize unique chemical scaffolds – remain difficult to predict and identify. Here, our results demonstrate how an unbiased, evolution-guided comparative analysis can identify cryptic elements of plant specialized metabolism regardless of functional annotation. A key example of this was the discovery of an essential transporter (NPF8.1) that connects extracellular (apoplastic) and intracellular (cytosolic) reactions in Lycopodium alkaloid biosynthesis. While the expression of NPF8.1 correlated well with that of LDC-1 in our original *P. tetrastichus* transcriptome (Pearson’s *r* = 0.93), it was not within the top 100 co-expressed genes (rank = 141). Typically, Agrobacterium transient expression experiments are limited to 30-40 gene constructs that can be tested in a single batch (*44*). Without any additional information, it was highly improbable that NPF8.1 would have been tested in the same batch combination with the next required biosynthetic enzyme (MDR-1). We emphasize that this comparative transcriptomic approach was crucial for identifying the missing components of Lycopodium alkaloid biosynthesis, which resolves a major bottleneck in understanding how clubmosses coordinate metabolism across cellular compartments to produce structurally-diverse chemical scaffolds.

The role of transporters in specialized metabolism is well-established as a mechanism to facilitate metabolite movement between cellular compartments and among biosynthetic cells (*31*, *35*). However, the role of the apoplast in plant specialized metabolism seems to be less well-defined. Indeed, while the apoplast is home to many essential processes in plants (e.g. lignin polymerization) (*45*), there are relatively few examples of this compartment being used as a dedicated location for specialized metabolite biosynthesis (*46*, *47*). The acidic, oxidizing environment of the apoplast may facilitate certain biosynthetic transformations (*45*) including the scaffold forming reactions for Lycopodium alkaloids. However, we note here that the use of the apoplast for biosynthesis also presents an organizational challenge to the cell if downstream reactions occur in the cytosol. This is particularly applicable to alkaloids, as their amine functional groups are predominantly charged in acidic cellular compartments like the apoplast and vacuole, thus limiting diffusion across membranes and causing these compartments to act as a “pH trap”. Indeed, our data suggest that this occurs in Lycopodium alkaloid biosynthesis, where **9** is effectively trapped in the apoplast following its production by CAL enzymes, thus requiring a transporter to facilitate its movement back into the cell (**Fig 4A**). Given that the cytosol of plant cells is bounded on the exterior by the acidic apoplast and on the interior by the acidic vacuole, NPF family transporters, which couple a proton gradient to metabolite transport (*32*), provide a solution for concentrating alkaloids into this compartment by utilizing the pH gradient maintained by cells. In Lycopodium alkaloid biosynthesis, the observed localization of NPF8.1 on both the tonoplast and plasma membranes is particularly compelling, as it could provide a single gene solution to the issue of pH trapping across multiple, acidic subcellular compartments. We anticipate that similar transport mechanisms will be an essential consideration in the future discovery of alkaloid biosynthesis pathways, particularly if reactions are compartmentalized in acidic cellular environments.

Beyond establishing pathways to two major Lycopodium alkaloids, our study revealed auxiliary proteins from multiple, functionally-distinct protein families that increase biosynthetic yield. This includes a transporter from the well-documented ABC family (*35*), as well as a fourth CAL from the α-carbonic anhydrase family, which has recently emerged as a prominent, expanded group of biosynthetic enzymes in clubmosses (*22*). However, the most unusual auxiliary proteins we identified were LABF-1 and LABF-2, which do not have obvious sequence homology to any proteins previously described in plant metabolism. For LABF-1, which belongs to the antibiotic biosynthesis monooxygenase (ABM) family, the closest non-clubmoss homologs in the NCBI database (via PSI-BLAST) are from cyanobacteria. Though LABF-2 does not have a predicted protein family annotation, PSI-BLAST searches indicated that this protein has homology to LABF-1 (e-value = 1.3×10^-10^). Thus, despite having very low amino acid identity (∼25%), we conclude that LABF-2 is likely related to the ABM family or its parent dimeric α-β barrel superfamily (IPR011008). While many dimeric α-β barrel proteins are characterized in polyketide and antibiotic biosynthesis in microbes (*40*, *48–50*), only one member of this superfamily has previously been described in plants (*51*). Our results suggest a broader role for dimeric α-β barrel proteins in plant natural product biosynthesis than previously recognized (*42*). More broadly, there are likely many protein families with roles in plant specialized metabolism that are yet to be identified, and the use of techniques that are agnostic to functional annotation – such as the evolution-guided approach used here – will be paramount for uncovering novel components of biosynthesis.

In summary, we have not only identified the unknown scaffold-forming enzymes for both HupA (**18**) and lycopodine (**22**), but also revealed how Lycopodium alkaloid biosynthesis is coordinated among cellular compartments in specialized biosynthetic cells. Collectively, our results provide a foundation for the metabolic engineering of the medicinal Lycopodium alkaloid HupA (**18**) that considers both enzyme catalysts and metabolic organization, thereby enabling a potential alternative to the overharvesting of HupA-producing clubmosses (*5*, *6*). More broadly, the discovery of pathways for both the lycodane and lycopodane scaffolds opens the door to studies on the biosynthesis and biological function for hundreds of downstream Lycopodium alkaloids (*3*), which we expect to play critical ecological roles that have contributed to the evolutionary success of the clubmoss family.

## Acknowledgments

We are grateful to Prof. Elín Soffía Ólafsdóttir and Maonian Xu (University of Iceland) for providing us with lycopodine (**22**) and Prof. Yun Yee Low (University of Malaya) for providing flabellidine (**12**). We acknowledge the use of Harvard Forest resources and specifically thank Audrey Barker Plotkin and Greta VanScoy for their assistance with locating and identifying clubmoss specimens. We also thank the entire Nett lab group for assistance with clubmoss harvesting and sample preparation, and the Balskus lab for assistance with small molecule purification. We thank Prof. George Lomonosoff for providing us with the pEAQ-HT plasmid and Prof. David Nelson (University of Tennessee) for assisting with cytochrome P450 nomenclature. We are grateful to the Harvard Center for Biological Imaging (HCBI) for assistance with microscopy, the Bauer Core Facility for guidance on single nuclei RNA-sequencing, and the CCB Nuclear Magnetic Resonance Core Facility for NMR analysis.

## Author Contributions

RSN and EAF conceptualized the project. All experiments and data analysis were carried out by EAF, with the exception of snRNAseq, which was led by CYK. RSN and EAF wrote the manuscript with editing from CYK.

## Data, code, and materials availability

All data, code, and materials generated through this study will be made available upon request, following publication. Code used in this study is provided as a Supplementary Material. Raw sequencing data from *D. obscurum* (short read, long read, snRNAseq) will be deposited to the National Center for Biotechnology Information (NCBI) database and made available upon publication. Coding sequences for all genes functionally characterized in this study (from *P. tetrastichus* and *D. obscurum*) will be deposited to GenBank and made available upon publication.

## Supplementary Materials

Materials and Methods

Figs. S1 to S38

Supplementary Methods

Supplementary Code

Supplementary Scheme

References (*52–72*)

## References and Notes

1. J. R. Jacobowitz, J. K. Weng, Exploring uncharted territories of plant specialized metabolism in the postgenomic era. Annu. Rev. Plant Biol. 71, 631–658 (2020).

2. J. Ziegler, P. J. Facchini, Alkaloid biosynthesis: metabolism and trafficking. Annu. Rev. Plant Biol. 59, 735–769 (2008).

3. Z. J. Zhang, S. Jiang, Q. S. Zhao, The chemistry and biology of Lycopodium alkaloids. Chem. Biodivers. 21, e202400954 (2024).

4. X. Ma, D. R. Gang, The Lycopodium alkaloids. Nat. Prod. Rep. 21, 752–772 (2004).

5. X. Ma, C. Tan, D. Zhu, D. R. Gang, A survey of potential huperzine A natural resources in China: The Huperziaceae. J. Ethnopharmacol. 104, 54–67 (2006).

6. X. Ma, C. Tan, D. Zhu, D. R. Gang, Is there a better source of huperzine A than *Huperzia serrata*? Huperzine A content of Huperziaceae species in China. J. Agric. Food Chem. 53, 1393–1398 (2005).

7. P. Siengalewicz, J. Mulzer, U. Rinner, “Lycopodium alkaloids - synthetic highlights and recent developments” in Alkaloids: Chemistry and Biology, H.-J. Knolker, Ed. (Elsevier Inc., Oxford, UK, 2013; 10.1016/B978-0-12-407774-4.00001-7) vol. 72, pp. 1–151.

8. J.-S. Liu, Y.-L. Zhu, C.-M. Yu, Y.-Z. Zhou, Y.-Y. Han, F.-W. Wu, B.-F. Qi, The structures of huperzine A and B, two new alkaloids exhibiting marked anticholinesterase activity. Can. J. Chem. 64, 837–839 (1986).

9. Y. E. Wang, D. X. Yue, X. C. Tang, Anti-cholinesterase activity of huperzine A. Acta Pharmacol. Sin. 7, 110–113 (1986).

10. G. D. Ainge, S. D. Lorimer, P. J. Gerard, L. D. Ruf, Insecticidal activity of huperzine A from the New Zealand clubmoss, *Lycopodium varium*. J. Agric. Food Chem. 50, 491–494 (2002).

11. C. X. Ban, S. F. Xiao, X. Lin, T. Wang, Q. Qiu, M. J. Zhu, X. Li, Clinicians’ prescription preferences for treating patients with Alzheimer’s disease in Shanghai. Transl. Neurodegener. 5 (2016).

12. K. Bödeker, Lycopodin, das erste Alkaloïd der Gefässkryptogamen. Justus Liebigs Ann. Chem. 208, 363–367 (1881).

13. M. Castillo, R. N. Gupta, D. B. MacLean, I. D. Spenser, Biosynthesis of lycopodine from lysine and acetate. The pelletierine hypothesis. Can. J. Chem. 48, 1893–1903 (1970).

14. M. Castillo, R. N. Gupta, Y. K. Ho, D. B. MacLean, I. D. Spenser, Biosynthesis of lycopodine. Incorporation of 1-piperideine and of pelletierine. Canadian Journal of Botany 48, 2911–2918 (1970).

15. J.-C. Braekman, R. N. Gupta, D. B. MacLean, I. D. Spenser, Biosynthesis of lycopodine. Pelletierine as an obligatory intermediate. Can. J. Chem. 50, 2591–2602 (1972).

16. T. Hemscheidt, I. D. Spenser, Biosynthesis of lycopodine: incorporation of acetate via an intermediate with C_2v_ symmetry. J. Am. Chem. Soc. 115, 3020–3021 (1993).

17. W. D. Marshall, I. D. Spenser, T. T. Nguyen, D. B. MacLean, Biosynthesis of lycopodine. The question of the intermediacy of piperidine-2-acetic acid. Can. J. Chem. 53, 41–50 (1975).

18. T. Hemscheidt, I. D. Spenser, A classical paradigm of alkaloid biogenesis revisited: acetonedicarboxylic acid as a biosynthetic precursor of lycopodine. J. Am. Chem. Soc. 118, 1799–1800 (1996).

19. S. Bunsupa, K. Katayama, E. Ikeura, A. Oikawa, K. Toyooka, K. Saito, M. Yamazaki, Lysine decarboxylase catalyzes the first step of quinolizidine alkaloid biosynthesis and coevolved with alkaloid production in Leguminosae. Plant Cell 24, 1202–1216 (2012).

20. B. Xu, L. Lei, X. Zhu, Y. Zhou, Y. Xiao, Identification and characterization of L-lysine decarboxylase from *Huperzia serrata* and its role in the metabolic pathway of lycopodium alkaloid. Phytochemistry 136, 23–30 (2017).

21. J. Wang, Z.-K. Zhang, F.-F. Jiang, B.-W. Qi, N. Ding, S. Y. Y. Hnin, X. Liu, J. Li, X. Wang, P.-F. Tu, I. Abe, H. Morita, S.-P. Shi, Deciphering the biosynthetic mechanism of pelletierine in *Lycopodium* alkaloid biosynthesis. Org. Lett. 22, 9725–8729 (2020).

22. R. S. Nett, Y. Dho, Y.-Y. Low, E. S. Sattely, A metabolic regulon reveals early and late acting enzymes in neuroactive Lycopodium alkaloid biosynthesis. Proceedings of the National Academy of Sciences 118, e2102949118 (2021).

23. R. S. Nett, Y. Dho, C. Tsai, D. Passow, J. Martinez Grundman, Y. Y. Low, E. S. Sattely, Plant carbonic anhydrase-like enzymes in neuroactive alkaloid biosynthesis. Nature 624, 182–191 (2023).

24. S. N. Alam, A. H. Adams, D. B. MacLean, Lycopodium alkaloids. XV. Structure and mass spectra of some minor alkaloids of *L. flabelliforme*. Can. J. Chem. 42, 2456–2466 (1964).

25. J. S. Y. Yeap, K. H. Lim, K. T. Yong, S. H. Lim, T. S. Kam, Y. Y. Low, *Lycopodium* alkaloids: Lycoplatyrine A, an unusual lycodine-piperidine adduct from *Lycopodium platyrhizoma* and the absolute configurations of lycoplanine D and lycogladine H. J. Nat. Prod. 82, 324–329 (2019).

26. W. Testo, A. Field, D. Barrington, Overcoming among-lineage rate heterogeneity to infer the divergence times and biogeography of the clubmoss family Lycopodiaceae. J. Biogeogr. 45, 1929–1941 (2018).

27. A. R. Petlewski, D. A. Hauser, M. Kim, J. Schmutz, J. Grimwood, F. W. Li, Re-evaluating the systematics of *Dendrolycopodium* using restriction-site associated DNA-sequencing. Front. Plant Sci. 13 (2022).

28. W. A. Ayer, G. C. Kasitu, Some new Lycopodium alkaloids. Can. J. Chem. 67, 1077–1086 (1989).

29. R. S. Nett, X. Guan, K. Smith, A. M. Faust, E. S. Sattely, C. R. Fischer, D_2_O labeling to measure active biosynthesis of natural products in medicinal plants. AIChE Journal 64, 4139–4330 (2018).

30. K. L. Kavanagh, H. Jörnvall, B. Persson, U. Oppermann, Medium- and short-chain dehydrogenase/reductase gene and protein families: The SDR superfamily: Functional and structural diversity within a family of metabolic and regulatory enzymes. Cellular and Molecular Life Sciences 65, 3895–3906 (2008).

31. C. Kanstrup, H. H. Nour-Eldin, The emerging role of the nitrate and peptide transporter family: NPF in plant specialized metabolism. Curr. Opin. Plant Biol. 68, 102243 (2022).

32. Y.-F. Tsay, C.-C. Chiu, C.-B. Tsai, C.-H. Ho, P.-K. Hsu, Nitrate transporters and peptide transporters. FEBS Lett. 581, 2290–2300 (2007).

33. B. R. Lichman, The scaffold-forming steps of plant alkaloid biosynthesis. Nat. Prod. Rep. 38, 103–129 (2021).

34. M. T. Ødum, F. Teufel, V. Thumuluri, J. J. Almagro Armenteros, A. R. Johansen, O. Winther, H. Nielsen, DeepLoc 2.1: multi-label membrane protein type prediction using protein language models. Nucleic Acids Res. 52, W215–W220 (2024).

35. B. A. Halkier, D. Xu, The ins and outs of transporters at plasma membrane and tonoplast in plant specialized metabolism. Nat. Prod. Rep. 39, 1483–1491 (2022).

36. B. K. Nelson, X. Cai, A. Nebenführ, A multicolored set of in vivo organelle markers for co-localization studies in Arabidopsis and other plants. Plant Journal 51, 1126–1136 (2007).

37. N. Y. Komarova, S. Meier, A. Meier, M. S. Grotemeyer, D. Rentsch, Determinants for *Arabidopsis* peptide transporter targeting to the tonoplast or plasma membrane. Traffic 13, 1090–1105 (2012).

38. C. Li, J. C. Wood, A. H. Vu, J. P. Hamilton, C. E. Rodriguez Lopez, R. M. E. Payne, D. A. Serna Guerrero, K. Gase, K. Yamamoto, B. Vaillancourt, L. Caputi, S. E. O’Connor, C. Robin Buell, Single-cell multi-omics in the medicinal plant *Catharanthus roseus*. Nat. Chem. Biol. 19, 1031–1041 (2023).

39. P. A. Rea, Plant ATP-binding cassette transporters. Annu. Rev. Plant Biol. 58, 347–375 (2007).

40. G. Sciara, S. G. Kendrew, A. E. Miele, N. G. Marsh, L. Federici, F. Malatesta, G. Schimperna, C. Savino, B. Vallone, The structure of ActVA-Orf6, a novel type of monooxygenase involved in actinorhodin biosynthesis. EMBO J. 22, 205–215 (2003).

41. U. Gani, R. A. Vishwakarma, P. Misra, Membrane transporters: the key drivers of transport of secondary metabolites in plants. Plant Cell Rep. 40, 1–18 (2021).

42. M. Dastmalchi, Elusive partners: a review of the auxiliary proteins guiding metabolic flux in flavonoid biosynthesis. Plant Journal 108, 314–329 (2021).

43. K. S. Singh, J. J. J. van der Hooft, S. C. M. van Wees, M. H. Medema, Integrative omics approaches for biosynthetic pathway discovery in plants. Nat. Prod. Rep. 39, 1876–1896 (2022).

44. E. D. Carlson, J. Rajniak, E. S. Sattely, Multiplicity of the *Agrobacterium* infection of *Nicotiana benthamiana* for transient DNA delivery. ACS Synth. Biol. 12, 2329–2338 (2023).

45. R. A. Dixon, J. Barros, Lignin biosynthesis: old roads revisited and new roads explored. Open Biol. 9, 190215 (2019).

46. J.-L. Lin, W.-K. Wu, G.-B. Nie, J.-X. Li, X. Fang, Y.-G. Sheng, M.-M. Wang, Q.-Y. Zheng, X.-X. Guo, J.-F. Huang, L.-Y. Ma, L.-J. Wang, J.-X. Liu, S.-S. Wang, B. Xu, Y. Gao, Y. Li, D. Wang, C. Martin, X.-Y. Chen, J.-Q. Huang, A dirigent protein redirects extracellular terpenoid metabolism for defense against biotic challenges. Nat. Commun.v 16, 9270 (2025).

47. L. B. Davin, N. G. Lewis, Dirigent proteins and dirigent sites explain the mystery of specificity of radical precursor coupling in lignan and lignin biosynthesis. Plant Physiol. 123, 453–462 (2000).

48. Z. Qin, R. Devine, M. I. Hutchings, B. Wilkinson, A role for antibiotic biosynthesis monooxygenase domain proteins in fidelity control during aromatic polyketide biosynthesis. Nat. Commun. 10, 3611 (2019).

49. M. M. Machovina, R. J. Usselman, J. L. Du Bois, Monooxygenase substrates mimic flavin to catalyze cofactorless oxygenations. Journal of Biological Chemistry 291, 17816–17828 (2016).

50. R. N. Patkar, P. I. Benke, Z. Qu, Y. Y. Constance Chen, F. Yang, S. Swarup, N. I. Naqvi, A fungal monooxygenase-derived jasmonate attenuates host innate immunity. Nat. Chem. Biol. 11, 733–740 (2015).

51. S. J. Gagne, J. M. Stout, E. Liu, Z. Boubakir, S. M. Clark, J. E. Page, Identification of olivetolic acid cyclase from *Cannabis sativa* reveals a unique catalytic route to plant polyketides. Proceedings of the National Academy of Sciences 109, 12811–12816 (2012).

52. J.-M. Jiang, D. Xia, X.-L. Zhu, D. Zhu, X.-W. Yang, K. Pan, Lycophlegmarinines A–F, new *Lycopodium* alkaloids from *Phlegmariurus phlegmaria*. Tetrahedron 114, 132782 (2022).

53. L. Fu, B. Niu, Z. Zhu, S. Wu, W. Li, CD-HIT: Accelerated for clustering the next-generation sequencing data. Bioinformatics 28, 3150–3152 (2012).

54. A. M. Bolger, M. Lohse, B. Usadel, Trimmomatic: A flexible trimmer for Illumina sequence data. Bioinformatics 30, 2114–2120 (2014).

55. N. L. Bray, H. Pimentel, P. Melsted, L. Pachter, Near-optimal probabilistic RNA-seq quantification. Nat. Biotechnol. 34, 525–527 (2016).

56. M. J. L. de Hoon, S. Imoto, J. Nolan, S. Miyano, Open source clustering software. Bioinformatics 20, 1453–1454 (2004).

57. D. M. Emms, S. Kelly, OrthoFinder: Phylogenetic orthology inference for comparative genomics. Genome Biol. 20, 1–14 (2019).

58. J. G. Yu, J. Y. Tang, R. Wei, M. F. Lan, R. C. Xiang, X. C. Zhang, Q. P. Xiang, The first homosporous lycophyte genome revealed the association between the recent dynamic accumulation of LTR-RTs and genome size variation. Plant Mol. Biol. 112, 325–340 (2023).

59. C. Li, D. Wickell, L. Y. Kuo, X. Chen, B. Nie, X. Liao, D. Peng, J. Ji, J. Jenkins, M. Williams, S. Shu, C. Plott, K. Barry, S. Rajasekar, J. Grimwood, X. Han, S. Sun, Z. Hou, W. He, G. Dai, C. Sun, J. Schmutz, J. H. Leebens-Mack, F. W. Li, L. Wang, Extraordinary preservation of gene collinearity over three hundred million years revealed in homosporous lycophytes. Proc. Natl. Acad. Sci. U. S. A. 121 (2024).

60. R. Burstein, M. Ashina, K. Nozaki, R. P. Kraig, N. T. Zervas, M. A. Moskowitz, S. A. Raymond, R. Burstein, The Selaginella genome identifies genetic changes associated with the evolution of vascular plants. Science (1979). 339, 1095–1099 (2013).

61. J. Cui, Y. Zhu, H. Du, Z. Liu, S. Shen, T. Wang, W. Cui, R. Zhang, S. Jiang, Y. Wu, X. Gu, H. Yu, Z. Liang, Chromosome-level reference genome of tetraploid *Isoetes sinensis* provides insights into evolution and adaption of lycophytes. Gigascience 12 (2022).

62. A. Stoddard, V. Rolland, I see the light! Fluorescent proteins suitable for cell wall/apoplast targeting in Nicotiana benthamiana leaves. Plant Direct 3 (2019).

63. C. A. Smith, E. J. Want, G. O’Maille, R. Abagyan, G. Siuzdak, XCMS: Processing mass spectrometry data for metabolite profiling using nonlinear peak alignment, matching, and identification. Anal. Chem. 78, 779–787 (2006).

64. N. Nayak, S. Mehrotra, R. Mehrotra, Isolation and transfection of *Nicotiana benthamiana* mesophyll protoplasts for fluorescent protein visualization (2024). 10.17504/protocols.io.e6nvw1my9lmk/v1.

65. M. Martin, Cutadapt removes adapter sequences from high-throughput sequencing reads. EMBnet. J. 17, 10 (2011).

66. W. Shen, B. Sipos, L. Zhao, SeqKit2: A Swiss army knife for sequence and alignment processing. iMeta 3 (2024).

67. B. Kaminow, D. Yunusov, A. Dobin, STARsolo: accurate, fast and versatile mapping/quantification of single-cell and single-nucleus RNA-seq data. bioRxiv, doi: 10.1101/2021.05.05.442755 (2021).

68. Y. Hao, T. Stuart, M. H. Kowalski, S. Choudhary, P. Hoffman, A. Hartman, A. Srivastava, G. Molla, S. Madad, C. Fernandez-Granda, R. Satija, Dictionary learning for integrative, multimodal and scalable single-cell analysis. Nat. Biotechnol. 42, 293–304 (2024).

69. M. D. Young, S. Behjati, SoupX removes ambient RNA contamination from droplet-based single-cell RNA sequencing data. Gigascience 9 (2020).

70. C. Hafemeister, R. Satija, Normalization and variance stabilization of single-cell RNA-seq data using regularized negative binomial regression. Genome Biol. 20, 296 (2019).

71. C. S. McGinnis, L. M. Murrow, Z. J. Gartner, DoubletFinder: Doublet detection in single-Cell RNA sequencing data using artificial nearest neighbors. Cell Syst. 8, 329–337.e4 (2019).

72. H.-C. Xue, Z.-G. Xu, Y.-J. Liu, L. Wang, X. Ming, Z.-Y. Wu, H. Lian, Y.-W. Han, J. Xu, Z.-D. Zhang, Q.-L. Shao, K. Liu, F.-X. Wang, A.-H. Wang, J. Zhao, J. Zhang, J. Chen, Y. Mao, J.-W. Wang, A unified cell atlas of vascular plants reveals cell-type foundational genes and accelerates gene discovery. Cell 188, 6370–6390.e29 (2025)

