## Supplementary Materials for "Ancient origins of alkaloid biosynthesis in medicinal clubmosses"

**The PDF file includes:**

Materials and Methods
Figs. S1 to S38
Supplementary Methods
Supplementary Code description
Supplementary Scheme
References (*52-72*)

**Other Supplementary Materials for this manuscript include the following:**

Supplementary Code

### Supplementary Materials

#### Materials and Methods

##### *Dendrolycopodium obscurum* plant material

For all experiments with clubmoss tissue except for single nuclei isolation, whole-plant material of *Dendrolycopodium obscurum* was collected from Harvard Forest (Petersham, MA, USA) on June 27, 2023. For RNA-sequencing, plant tissue was separated into the following tissue types: new growth, sub-new growth leaf, sub-new growth stem, old leaf, old stem, newly growing rhizome, and roots (**Fig S2**). Four replicates were isolated from the same above-ground shoot for all above-ground tissues. Similarly, four replicates from the most closely connected below-ground tissue were isolated for below-ground tissues. All tissue was rinsed with deionized water thoroughly. Immediately after dissection, tissues were frozen on a heat block inserted into dry ice for transport and stored at −80 °C until processing. For isotope-labeling experiments, an above-ground shoot from a plant harvested at the same location and time was used.

##### *Nicotiana benthamiana* growth conditions

*Nicotiana benthamiana* plants were grown in growth chambers at 30 °C with a 16 hour light and 8 hour dark cycle without humidity control. For soil, Pro-Mix HP AGTIV Reach was used with Southern Ag PowerPak 20-20-20 Water Soluble Fertilizer with micronutrients. Seeds were germinated under humidity domes for the first week and then without domes for the remaining growth period. Soil was kept damp with routine watering. In general, 4-5 week old plants were used for *Agrobacterium*-mediated transient expression.

##### Microbial strains

*Escherichia coli* strain NEB 10-beta (New England Biolabs) was used for routine cloning of gene candidates into the pEAQ-HT vector. *Agrobacterium tumefaciens* strain GV3101 was used for transient expression in *N. benthamiana*. For attempts to express NPF8.1 in yeast, Invitrogen™ INV*Sc*1 *S. cerevisiae* was used with the pESC-leu2D plasmid purchased from Addgene (Plasmid #20120).

##### Chemicals and reagents

Unless otherwise noted, common reagents were purchased from commercial vendors and used without further purification. Chemical standards used for LC–MS/MS metabolite identification included: (-)-huperzine A (**18**) from Apex Biotechnology LLC; huperzine C (**17**) from Shanghai Tauto Biotech Co., Ltd; huperzine B (**16**) from MilliporeSigma; lycophlegmarinine D (**14**) isolated from *Phlegmariurus phlegmaria* and provided by Ke Pan (China Pharmaceutical University) (*52*); lycopodine (**22**) isolated from *Huperzia selago* and provided by Elín Soffía Ólafsdóttir and Maonian Xu; and flabellidine (**12**) isolated from *Lycopodium platyrhizoma* provided by Professor Yun Yee Low (University of Malaya) (*25*). Flabelline (**20**) was obtained by purification from *N. benthamiana* extracts as described below. Sodium borodeuteride ($NaBD_4$,

99% D, 95% chemical purity; DLM-226-1) and deuterium oxide (99% D; DLM-7005) were purchased from Cambridge Isotope Laboratories.

*D. obscurum* tissue sampling for metabolite extraction

Frozen *D. obscurum* tissues were homogenized in a mortar and pestle under liquid nitrogen. Metabolites were extracted with cold 80% (v/v) methanol (20% water) at 50 µL per mg wet tissue, and extracts were incubated on ice for 1 h. Extracts were centrifuged at 10,000 × g for 5 min to pellet debris, and supernatants were diluted into the appropriate solvent for LC–MS analysis via both C18 (water +0.1% formic acid) and HILIC (acetonitrile +0.1% formic acid) analysis.

$D_2O$ labeling to identify sites of active Lycopodium alkaloid biosynthesis

To identify tissues actively synthesizing Lycopodium alkaloids, freshly excised *D. obscurum* shoots from Harvard Forest were placed in 50 mL conical tubes containing 20% (v/v) $D_2O$ in water (*29*), which was replenished as needed during the labeling period. The plants were maintained at room temperature with a 16 hour light and 8 hour dark cycle. After 14 days, tissues were sectioned into three biological replicates per tissue type in the same manner as in preparation for RNA-seq (**Fig S2**) and homogenized under liquid nitrogen. Metabolites were extracted with 80% methanol at 50 µL per mg wet tissue for 2 h, then diluted into water with 0.1% formic acid (FA) for LC–MS analysis. For quantification of deuterium incorporation into flabellidine (**12**), the $[M+D]^+$ = *m/z* 290.2338 (M1) ion abundance was extracted and the estimated natural $^{13}C$ isotope abundances based on the $[M+H]^+$ = *m/z* 289.2275 (M0) ion abundances were subtracted. The ratios of M1 to M0 without estimated $^{13}C$ isotope removal were also calculated.

RNA extraction, library preparation, and sequencing

*D. obscurum* tissues were homogenized to a fine powder under liquid nitrogen with a mortar and pestle and portions of each tissue replicate were split for metabolomic analysis and RNA extraction. RNA was extracted using the Qiagen RNeasy Plant Mini Kit (cat. 74904) with on-column DNase digestion (Qiagen RNase-Free DNase Set, cat. 79254) following the manufacturer's instructions. RNA concentration was measured by absorbance at 260 nm using a NanoDrop spectrophotometer and RNA was stored at −80°C. This was performed for all tissue types (new growth, sub new growth leaf, sub new growth stem, old leaf, old stem, newly growing rhizome, roots) with four replicates. Older rhizome tissue that did not seem to be actively growing yielded no detectable RNA.

For long-read sequencing, single replicates from each tissue type were pooled in equimolar ratios and sequenced by the Bauer Core Facility at Harvard University using PacBio Iso-Seq (SMRT Cell 8M) on a Sequel IIe, yielding 3-4 million reads. For short-read sequencing, individual RNA samples were shipped to Genewiz for library preparation (NEBNext Ultra II with poly(A) selection) and sequenced on an Illumina HiSeq platform to generate 150 bp paired-end reads (~20 million reads per sample).

Transcriptome assembly and gene-expression quantification

PacBio Iso-Seq data were processed using the recommended Iso-Seq pipeline (https://isoseq.how/), including cDNA primer removal with lima, poly(A) tail and concatemer removal using isoseq refine, and isoform-level clustering using isoseq cluster2. After clustering, 231,379 high-quality transcripts were obtained. Transcripts were then collapsed at 99% sequence identity using CD-HIT-EST (*53*) (word size n=5) to generate a non-redundant set of 69,714 transcripts.

Short-read RNA-seq reads were adapter-trimmed using Trimmomatic (*54*) and pseudoaligned to the CD-HIT-collapsed Iso-Seq transcriptome using kallisto (*55*) with 100 bootstrap replicates. For gene expression analysis, Pearson's *r* calculations were performed in Microsoft Excel using the $log_2$-transformed TPM estimations from kallisto across all samples. All scripts used for transcriptome assembly, quantification, and downstream analyses are available at (**Supplementary Code**).

Comparative co-expression and evolutionary conservation analysis for gene candidate selection

To prioritize gene candidates, the *P. tetrastichus* lysine decarboxylase (*Pt*LDC-1) was used as a BLAST query against the *D. obscurum* transcriptome to identify a putative ortholog (*Do*LDC-1). The hit with the lowest e-value was assumed to be *Do*LDC-1, and transcripts that strongly co-expressed with *Do*LDC-1 were extracted (Pearson's r > 0.6; 6,651 transcripts). Cluster 3.0 (*56*) was used to perform hierarchical clustering of the genes with Pearson's r > 0.6 using the average linkage algorithm based on uncentered correlation of centered and normalized $log_2$ expression profiles. This identified a minimal cluster (146 transcripts) containing orthologs of the known core biosynthetic genes previously identified from *P. tetrastichus (22, 23)*. This clustering was visualized in TreeView software (https://jtreeview.sourceforge.net/).

From the 146-transcript cluster, 34 candidate gene orthologs from *P. tetrastichus* were selected based upon the following criteria: (i) they were highly conserved between *D. obscurum* and *P. tetrastichus* (based upon tblastn, e-value < 1e−70); and (ii) the gene expression of the presumed orthologs in *P. tetrastichus* (selected here as the best BLAST hits within a bitscore threshold of 15 of the hit with highest bitscore) correlated to *Pt*LDC-1 with Pearson's r > 0.8. For visualization of the correlations of the selected genes, orthogroups were inferred using OrthoFinder 2.5.5 (*57*) based on proteomes from five clubmoss species (*D. obscurum, P. tetrastichus, Lycopodium clavatum, Huperzia asiatica, and Diphasiastrum complanatum*) and two lycophyte outgroups (*Isoetes sinensis, Selaginella moellendorffii)*. The *P. tetrastichus* proteome was previously generated from a long-read transcriptome (*22*). The *D. obscurum* proteome was generated similarly using Transdecoder with the CD99 long-read transcriptome generated in this work. The proteomes for all other species in the analysis were previously generated from reference genomes (*58–61*). For Pearson's correlations of biosynthetic genes between *P. tetrastichus* and *D. obscurum* (shown in **Figure 1C**), the orthogroups containing known *P. tetrastichus* biosynthetic genes were identified, and the transcript within each orthogroup from both plants that had the highest correlation to the corresponding species' LDC ortholog was plotted.

##### Plant expression constructs

Previously cloned *P. tetrastichus* genes were used as described in prior work (*22*, *23*). For *D. obscurum* genes, cDNA was first synthesized from new-growth tissue RNA using the SuperScript™ III First-Strand Synthesis SuperMix. Coding sequences were then amplified from the cDNA through PCR using Q5 Hot Start High-Fidelity 2X Master Mix with primers introducing Gibson Assembly homology arms for the pEAQ-HT vector digested with AgeI/XhoI. When PCR amplification was unsuccessful, coding sequences were synthesized by Twist Bioscience with codon optimization for *N. benthamiana*. Fragments were assembled into the digested pEAQ-HT vector using NEBuilder HiFi DNA Assembly Master Mix following the manufacturer's protocol.

Gibson assemblies were transformed into NEB 10-beta *E. coli* using the heat shock protocol provided by the manufacturer, plated on LB agar containing kanamycin (50 µg/mL), and incubated overnight at 37 °C. The gene fragment inserted into pEAQ-HT vector was screened by colony PCR on transformed *E. coli* followed by agarose gel electrophoresis. Positive colonies were grown in 4 mL LB with kanamycin (50 µg/mL) overnight at 37 °C, and plasmids were purified by miniprep. Plasmid sequences were verified by whole-plasmid sequencing (Genewiz). Verified plasmids were transformed into *A. tumefaciens* GV3101 by freeze–thaw transformation and selected on LB agar containing kanamycin (50 µg/mL) and gentamicin (30 µg/mL) for 2 days at 30 °C. *Agrobacterium* transformants were verified by PCR and amplicon sequencing with Genewiz and grown in liquid culture with LB containing kanamycin (50 µg/mL) and gentamicin (30 µg/mL) for 1-2 days and then stored as 25% glycerol stocks at −80 °C.

##### Site-directed mutagenesis and fluorescent protein fusions

The mutant version of *Pt*NPF8.1 (*Pt*NPF8.1-L13A) was generated by PCR with the following primers: a 5' mutagenesis primer that simultaneously added Gibson homology arms to the pEAQ-HT vector (XhoI/AgeI digested) and introduced a mutation swapping leucine 13 to an alanine (TTA -> GCA), and a 3' standard primer that added Gibson homology arms. The gene fragment for *Pt*NPF8.1-L13A was then cloned as previously described using Gibson assembly into pEAQ-HT.

For fluorescent fusions, gene fragments were generated using PCR reactions with primers that generate Gibson homology overhangs linking the fluorophore to the coding sequence of the gene of interest with a flexible GSGS amino acid linker. Gibson homology overhangs for annealing into the pEAQ-HT backbone were also added onto the appropriate primers amplifying the 5' or 3' ends of what became the fused sequence during the Gibson reaction. All constructs reported in the paper were fused with mNeonGreen at the C-terminus of the fusion protein. mNeonGreen was selected as the fluorophore for its stability at different pHs that it might encounter when expressed in tobacco *(62)*. The mNeonGreen gene fragment was synthesized by Twist Bioscience with *N. benthamiana* codon optimization. To confirm that the fluorophore tagged version of *Pt*MDR-1 was functional, we expressed it on top of the pathway to **9** with *Pt*NPF8.1 and found it still enabled production of **10** (**Fig S26**). Similarly, we confirmed that fluorophore tagged versions of *Pt*NPF8.1

and *Pt*NPF8.1-L13A were still able to enable *Pt*MDR-1 activity during pathway reconstruction in *N. benthamiana* (**Fig S31**).

##### *Agrobacterium*-mediated transient expression in *Nicotiana benthamiana*

For all results, the genes cloned from *P. tetrastichus* were used in *Agrobacterium* transient expression unless explicitly stated otherwise. *Agrobacterium* strains were grown on LB agar or in LB liquid culture with appropriate antibiotic selection for 1–2 days. Liquid cultures were pelleted (8,000 × g, 5 min) and resuspended in 0.5 mL LB, pelleted again to reduce carryover antibiotics, and resuspended in induction medium (10 mM MES, 10 mM $MgCl_2$, 150 µM acetosyringone, pH 5.6). Bacteria scraped from plates were similarly first resuspended in LB and then pelleted, followed by a final resuspension in induction medium. Suspensions were incubated at room temperature for at least 1 h prior to infiltration.

Optical density at 600 nm (OD600) was measured for each strain and strains were combined such that each strain's final OD600 was 0.3 (unless otherwise noted). Strain mixtures were infiltrated into the abaxial side of leaves from 4–5 week old *N. benthamiana* plants using a needleless syringe (~0.2 mL per leaf). For leaf-paired comparisons, different mixtures were infiltrated on opposite sides of the midrib of the same leaf. Plants were grown in the growth chambers for 3–11 days post-infiltration depending on the experiment (typically 3 days for early intermediates up to flabellidine (**12**); 5–7 days for later intermediates en route to HupA (**18**) or lycopodine (**22**); and 11 days for preparative-scale metabolite production).

##### Metabolite extraction from *N. benthamiana*

Infiltrated leaf tissue was excised, placed into pre-weighed 2 mL Safe-Lock tubes (Eppendorf), and flash-frozen in liquid nitrogen. Samples were lyophilized overnight to dryness and weighed to determine dry mass. Dried tissue was homogenized with 5 mm steel beads using a ball mill homogenizer (Retsch MM 400; 25 Hz, 2 min). Cold extraction solvent (water containing 0.1% formic acid unless otherwise noted) was added at 20 µL per mg dry tissue, mixed thoroughly, and incubated for ≥30 min. Samples were centrifuged (10,000 × g, 5 min, 4 °C), and supernatants were diluted into solvent appropriate for the LC method (water +0.1% formic acid for C18; acetonitrile +0.1% formic acid for HILIC/HILIC-Z) such that the quantified signals did not saturate the detector, occasionally requiring more than one dilution from the same sample. The diluted samples were then filtered using Multiscreen® 96 well Plate, 0.45 µm PTFE membrane plates from Sigma-Aldrich prior to LC–MS analysis.

##### LC–MS and MS/MS analysis

For all LC–MS analysis, specific parameters of methods used are listed in the **Supplemental Methods**. All analyses were performed on an Agilent 1290 Infinity II UHPLC coupled to an Agilent 6546 quadrupole time-of-flight (Q-TOF) mass spectrometer. Electrospray ionization (ESI) was performed in positive ion mode.

Early pathway intermediates (prior to **12**) were analyzed via HILIC or HILIC-Z chromatography using either an InfinityLab Poroshell 120 HILIC (2.1 × 150 mm, 1.9 µm) or a

Poroshell 120 HILIC-Z (2.1 × 100 mm, 1.9 µm) column. Mobile phases consisted of water and 9:1 acetonitrile:water, each containing 10 mM ammonium formate and 0.1% formic acid. Later intermediates (**12** and intermediates downstream of this) were analyzed using a combination of C18, HILIC, and HILIC-Z methods. A ZORBAX RRHD Eclipse Plus C18 (2.1 × 50 mm, 1.8 µm) column was used for C18 analyses. For targeted MS/MS ($MS^2$) analyses, collision energies were optimized for each target molecule and typically ranged from 10–40 V.

##### LC–MS data processing and quantification

Extracted ion chromatograms (EICs) were generated in Agilent MassHunter Qualitative Analysis using a ±20 ppm mass window unless otherwise specified. Peak areas were quantified using the Extract Chromatograms function with manual baseline correction when needed. For diastereomers of **5** with partially co-eluting peaks, peak areas were obtained by fitting skewed Gaussian functions using a custom Python script (**Supplementary Code**). For untargeted analyses, raw LC–MS files were converted to mzML format using MSConvert and processed in R using the XCMS package (*63*) using a custom script (**Supplementary Code**).

##### Chemical reduction and derivatization with sodium borodeuteride/borohydride

For structural analysis of double bond-containing pathway intermediates and to distinguish imine versus alkene reductions, metabolite extracts from infiltrated *N. benthamiana* tissue expressing the pathways to compounds **8**, **9**, **10**, and **11** were treated with sodium borodeuteride ($NaBD_4$). For $NaBD_4$ reactions, lyophilized and homogenized tissue was extracted with cold 100% methanol (20 µL per mg dry tissue) for 30 min on ice and clarified by centrifugation (10,000 × g, 5 min). Supernatants (400 µL) were split into treatment and no-reagent controls (200 µL each). An excess of $NaBD_4$ (10 mg) was added to treatment samples and reactions were incubated for 1 h on ice with caps vented in a fume hood. Water (100 µL) was then added and samples were incubated for 15 min to quench excess reagent. Samples were dried in a SpeedVac for 2 h at room temperature, resuspended in 500 µL of 90% acetonitrile (10% water) containing 0.1% formic acid, filtered, and analyzed by LC–MS (HILIC-Z).

##### In vitro MDR-1 activity assays using *N. benthamiana* lysates

To assess MDR-1 activity independent of NPF8.1, crude protein lysates were prepared from *N. benthamiana* leaves transiently expressing MDR-1, MDR-1 + NPF8.1, or GFP harvested at 3 days post infiltration (N=3 biological replicates). Tissue (0.05g) was homogenized in 500µL of ice-cold lysis buffer (100 mM Tris pH 8, 10 mM DTT, 10% glycerol, 1/10 of the plant leaf mass of polyvinylpyrrolidone) using a pipette tip for mechanical disruption. Protein lysate (49 µL) was then aliquoted into tubes containing 50 µL of a water extract of *N. benthamiana* expressing the pathway to **9** three days post infiltration (20 µL/mg dry tissue) and 1 µL of either 500 µM NADPH or water. The samples were incubated at room temperature for 2 hours before pelleting at 10,000 x g for 5 minutes and diluting 10-fold into cold acetonitrile +0.1% formic acid for analysis with LC-MS (HILIC-Z).

Co-infiltration of purified **9** to support MDR-1 substrate identity

To support that **9** is the substrate for MDR-1, *N. benthamiana* leaves expressing MDR-1 and NPF8.1 were co-infiltrated with HPLC purified and dried **9**. Although **9** degrades during purification (*23*) and thus the isolated product is not pure, no other known pathway metabolites were detected when the purified **9** sample was screened with LC-MS. The amount of **9** purified and co-infiltrated was too small to be quantified. To select an appropriate amount for detection of MDR-1 activity, we resuspended the purified **9** into *Agrobacterium* induction media (10 mM MES, 10 mM $MgCl_2$, 150 µM acetosyringone, pH 5.6) and empirically determined what dilution of this solution would reach the upper limit of detection by our LC-MS without saturating the detector. The sample was then further diluted into *Agrobacterium* induction media and infiltrated into leaf tissue during day 3 of *Agrobacterium* mediated expression of MDR-1 and NPF8.1 such that the estimated ion counts following extraction would be near this upper limit of detection assuming no consumption of **9**. On day 4 (1 day after substrate co-infiltration with a needleless syringe), the tissue was harvested and prepared for LC-MS as previously described for standard metabolite extractions from *N. benthamiana*.

Metabolite purification by flash chromatography and HPLC

For preparative metabolite isolation, pathway genes were transiently expressed in *N. benthamiana* for 11 days. Infiltrated tissue was harvested, lyophilized, and processed as described above. Approximately 5 g dry tissue was used for flabelline (**20**) purification, 0.5 g for flabellidine (**12**) purification, and 0.1 g for purification of compound **9**.

For flabelline (**20**) and flabellidine (**12**), tissue was extracted with 100% methanol (sufficient volume to submerge tissue) overnight at room temperature with shaking (30 rpm). A second extraction with half the initial methanol volume was performed for 1 h. Extracts were then combined, filtered, and dried by rotary evaporation at 30 °C. Residues were resuspended in 150 mL 3% (w/v) tartaric acid (aq) and extracted three times with 75 mL ethyl acetate to remove hydrophobic molecules. The aqueous phase was basified to ~pH 11 with 6 M NaOH and extracted five times with 75 mL ethyl acetate. This organic phase was dried of water with anhydrous magnesium sulfate and clarified with filter paper. The organic layer was then fully dried by rotary evaporation at 30 °C and resuspended in minimal methanol.

Resuspended extracts were dry-loaded onto a Biotage® Sfär KP-Amino Samplet® (5–10 g) and separated on a Biotage® Sfär KP-Amino D Duo 50 µm, 11 g cartridge using hexanes/ethyl acetate gradients on a Biotage Selekt instrument with 20 mL fractions (**Supplemental Methods**). Fractions were collected and screened by LC–MS. Fractions containing the metabolite of interest were pooled and dried by rotary evaporation at 30 °C. For purification of **9**, flash chromatography was not performed and the samples were immediately extracted with water with 0.1% formic acid on ice for 1 hour.

For final purification, pooled fractions were resuspended and injected repeatedly (100 µL per injection) onto an Agilent Infinity 1260 HPLC equipped with the following columns, as appropriate: Poroshell 120 HILIC-Z (2.1 × 150 mm, 2.7 µm) for compound **9**; Poroshell 120

HILIC (4.6 × 100 mm, 2.7 μm) for flabellidine (**12**); and Poroshell 120 EC-C18 (4.6 × 100 mm, 2.7 μm) for flabelline (**20**) (**Supplemental Methods**). Samples were resuspended in 90% acetonitrile (10% water) with 0.1% formic acid for HILIC/HILIC-Z runs and in water with 0.1% formic acid for C18 runs. Fractions were collected, screened by LC–MS, pooled, and lyophilized. The amounts of **9** and flabellidine (**12**) isolated were too small to quantify, but were the only identifiable peaks in the sample when analyzed with LC-MS (HILIC-Z chromatography for **9** and C18 for **12)**. Purification of flabelline (**20**) yielded approximately 1 mg.

<u>Base hydrolysis of flabellidine (**12**)</u>

Flabellidine (**12**, <1 mg, mass not quantifiable) purified from *N. benthamiana* extracts was resuspended in 4 mL ethanol and split into two samples (hydrolysis and control). For hydrolysis, 1.8 mL of 4 M KOH was added; for the control, 1.8 mL water was added. Samples were stirred at 100°C for 7 h. Ethanol was removed by rotary evaporation at 30°C. To isolate alkaloids from the KOH treated sample, 2 mL ethyl acetate was added and the organic phase was isolated, dried by rotary evaporation, and resuspended in 1 mL water with 0.1% FA. The control was treated with ethyl acetate to mimic handling and evaporated to dryness, then resuspended in 1 mL water with 0.1% formic acid. Both samples were diluted 10-fold into acetonitrile and analyzed by LC–MS (HILIC-Z).

<u>Structural confirmation of flabelline (**20**) through NMR</u>

Flabelline (**20**) was purified from *N. benthamiana* extracts as described above and lyophilized. Approximately 1 mg of purified flabelline (**20**) was resuspended in deuterated methanol (MeOD). One-dimensional spectra were acquired on a Bruker Avance 600 MHz spectrometer (Ascend magnet; 600.4 MHz for $^{1}$H) at ambient temperature. $^{1}$H spectra were acquired with 16 scans, a spectral width of 19.83 ppm, 65,536 time-domain points, an acquisition time of 2.75 s, and a relaxation delay of 1.0 s. $^{13}$C spectra were acquired using the zgpg30 pulse program with 4,096 scans, a spectral width of 236.5 ppm (35,714.3 Hz), 65,536 time-domain points, and an acquisition time of 0.92 s. Proton decoupling was applied during acquisition. Two-dimensional spectra were acquired on a JEOL 400 MHz instrument at ambient temperature. $^{1}$H–$^{1}$H COSY spectra were acquired with a $^{1}$H spectral width of 15 ppm using a data matrix of 1,024 × 256 complex points, 4 scans per increment, and a relaxation delay of 1.5 s. Multiplicity-edited $^{1}$H–$^{13}$C HSQC spectra were acquired with spectral widths of 15 ppm ($^{1}$H) and 170 ppm ($^{13}$C), using a data matrix of 1,024 × 128 complex points, 8 scans per increment, and a relaxation delay of 1.5 s. $^{1}$H–$^{13}$C HMBC spectra were acquired with spectral widths of 15 ppm ($^{1}$H) and 200 ppm ($^{13}$C), using a data matrix of 1,024 × 256 complex points, 32 scans per increment, and a relaxation delay of 1.5 s. NMR data were processed using MestReNova (Mestrelab Research). Chemical shifts were referenced to residual methanol-$d_4$ (MeOD): δ 3.31 ppm for $^{1}$H and δ 49.0 ppm for $^{13}$C. Automatic phase correction was applied. For baseline correction, the Whittaker baseline correction was applied. COSY spectra were symmetrized using the "Symmetrize (COSY-like)" function in MestReNova. Assignments supporting the structure of flabelline (**20**) were established by standard analysis of 1D and 2D spectra.

Fluorescent protein tagging and confocal microscopy

*N. benthamiana* leaf epidermal cells transiently expressing C-terminal mNeonGreen fusion proteins were imaged 72 h post-infiltration. Previously published subcellular marker constructs tagged with mCherry were used for co-localization analyses (*36*)· Leaf discs were excised from infiltrated regions, mounted in water under 22 mm coverslips with the abaxial side facing up, and imaged by confocal microscopy.

Confocal imaging was performed using a Zeiss LSM 980 laser-scanning confocal microscope mounted on an Axio Observer.Z1 inverted stand with a Plan-Apochromat 63×/1.40 NA oil immersion objective and GaAsP photomultiplier tube detectors. The pinhole was set to 1.0 Airy unit for all channels. Images were acquired with a pixel dwell time of 1.02 µs, 2× line averaging, z-step size of 0.23 µm, and channels acquired simultaneously. For mNeonGreen detection, a 488 nm laser was used for excitation with a detection window of 491-570 nm. For mCherry detection, a 594 nm laser was used for excitation with a detection window of 606-694 nm. Laser intensity was adjusted based on signal intensity in each leaf. For all experiments detecting a particular fluorophore fused protein, a control with no fluorophore fused protein was imaged and reported using the highest laser intensity setting of any leaf imaged within that same experiment (**Fig. S27, S29, & S30**).

Z-stacks were processed in ZEISS ZEN software. Maximum intensity projections were generated in ZEN. Background subtraction was applied using the rolling ball algorithm (radius = 100 pixels). For displaying images, brightness and contrast were adjusted in ZEN using linear histogram stretching. Adjustments were applied uniformly across all images within an experiment.

Protoplast isolation for localization clarification

To clarify the localization of NPF8.1–mNeonGreen, protoplasts were prepared from *Agrobacterium*-infected leaf tissue using an adaptation of a published protocol (*64*), with the exception of using our own *Agrobacterium* mediated infiltration procedure. Cell wall digestion buffer contained 20 mM MES (pH 5.7), 20 mM KCl, 400 mM mannitol, 1.5% cellulase R-10 (Yakult), 0.4% macerozyme R-10 (Yakult), 10 mM $CaCl_2$, and 0.1% BSA. Leaves expressing NPF8.1-mNeonGreen were cut into 0.5–1 mm strips and submerged in 3 mL digestion buffer in a 6-well plate. Buffer was vacuum-infiltrated into tissues for 2 min at ~600 mmHg vacuum and the vacuum was slowly released. Samples were incubated in the dark with shaking (50 rpm) at 30 °C for 3 h. Partially digested leaf strips were mounted in digestion buffer under 0.22 mm glass coverslips and intact protoplasts were imaged by confocal microscopy as described above.

Library construction and data processing for single-nucleus RNA sequencing

Nuclei were isolated from new leaf tissues of *D. obscurum* plants collected from Harvard Forest (324 North Main Street, Petersham, MA 01366). New leaf tissues were excised from the plants and stored in RNAlater solution at room temperature pre-processing. All nuclei extraction procedures were performed at 4 ºC and wide-bore pipette tips were used to handle the nuclei. Tissues were manually disrupted by two double edge razor blades (Feather) on a glass petri dish for 2 mins in 300 µL of nuclei isolation buffer (NIB). NIB consisted of 0.3 M sucrose, 1.25%

Ficoll, 2.5% Dextran T40, 15 mM Tris-HCl pH 8, 20 mM MES, 10 mM $MgCl_2$, 60 mM KCl, 15 mM NaCl, 0.5 mM spermine, 0.5 mM spermidine, and 0.1% Triton X-100, with 5 mM DTT, 1 mM PMSF, 1% Plant Protease Inhibitors (Sigma P9599), 0.8% BSA, and 0.2 U/µL Protector RNase Inhibitor (Roche) added immediately before use. The resulting solution containing chopped tissues was transferred to a pre-chilled 2 mL Dounce tissue grinder (Kimble, 885302), along with 400 µL of NIB used to wash and obtain the remaining tissues on the petri dish. Pestle A was used to further grind the samples by 10 up-down movements. The solution in the Dounce was incubated on ice to finish the lysis of the cells for 5 mins. 5 additional up-down movements with Pestle A were performed to release the rest of the nuclei. To remove debris, disrupted tissue solution was then passed through a pre-wet 20-µm cell strainer (CellTrics, 04-0042-2315) into a prechilled 1.5 mL Eppendorf with 500 µL of NIB to wash the remains of the Dounce tissue grinder. Nuclei were gently pelleted at 500 x *g* for 10 mins using a swinging bucket rotor. Supernatant was removed and nuclei were resuspended with 50 µL of nuclei resuspension buffer (NRB). NRB consisted of 0.3 M sucrose, 15 mM Tris HCl pH 8, 60 mM KCl, 15 mM NaCl, 0.5 mM spermine, 0.5 mM spermidine, and 15 mM MES, with 5 mM DTT, 1% Plant Protease Inhibitors (Sigma P9599), 2% BSA, and 0.2u/µL Protector RNase Inhibitor (Roche) added immediately before use. After nuclei isolation, Acridine Orange (AO)/Propidium Iodide (PI) dye was added to stain the nuclei. An automated cell counter (LUNA-FX7™ Logos Biosystems) was used to count the number of nuclei and quality control for nuclei integrity before loading into the 10x Genomics Chromium controller. Libraries were generated using the Chromium GEM-X Single Cell 3' Reagent Kits v4. Libraries were then sequenced on the NovaSeq X Series (Illumina).

For data processing, adapter and poly(A) trimming were performed on cDNA reads using Cutadapt (v5.2) (*65*). Bases with Phred quality score <30 were trimmed and reads shorter than 30 nucleotides were discarded, then read pairs were synchronized using SeqKit (v2.13.0) (*66*). STARsolo (v2.7.11b) (*67*) was used to align the read pairs against a *de novo* Iso-Seq transcriptome assembly of *D. obscurum* with full-length transcripts collapsed at 97% nucleotide identity using CD-HIT (v4.8.1) (*53*). A STAR reference was generated using options --runMode genomeGenerate --sjdbOverhang 89 --genomeSAindexNbases 12 and .gtf file generated from TransDecoder (v5.7.1) (https://github.com/TransDecoder/TransDecoder), and read alignment was conducted using options --alignIntronMax 1 --alignSJDBoverhangMin 999 --soloBarcodeReadLength 0 --soloUMIlen 12 --soloStrand Forward --soloCellFilter EmptyDrops_CR --soloFeatures Gene --soloMultiMappers EM --soloType CB_UMI_Simple. Downstream analyses were conducted using the R package Seurat (v5.4) (*68*). Ambient RNA contamination was removed using SoupX (v1.6.2) (*69*), then the data were normalized using the SCTransform method (*70*). Cells with fewer than 200 detected features (nFeature_RNA) were filtered out. For further processing, PCA was performed using the first 20 principal components and DoubletFinder (*71*) removed doublets in the Seurat object. 10,679 cells classified as singlets with 2,577 median UMI and 781 median genes were retained for all downstream analyses. All scripts and parameters are provided to ensure reproducibility (**Supplementary Code**).

To annotate cell clusters in the *D. obscurum* new leaf snRNA-seq dataset, we transferred cell-type information from the published *Lycopodium japonicum* shoot-apex atlas (*72*) using top differentially expressed marker genes for each of the 13 cell-types (EC Epidermal Cell, 82 genes; EC Guard Cell, 98 genes; Endodermis, 100 genes; MC Mesophyll/Cortex, 100 genes; PC G2/M Phase, 100 genes; PC S Phase, 100 genes; Unknown, 100 genes; VC Immature Phloem, 89 genes; VC Immature Xylem, 84 genes; VC Parenchyma Cell, 100 genes; VC Phloem Sieve Element, 100 genes; VC Vascular Cell Type I, 23 genes; and VC Xylem Tracheary Element, 100 genes). Mapping statistics and scoring for each marker set transfer are summarized in **Supplementary Code**.

To test whether the individual nuclei from cluster 9 (**Figure 3G**) express genes from both pre- and post-transporter lycopodine biosynthetic modules, the SCT data values were linearized and the mean expression of all genes in each module was calculated for each nucleus in the dataset. For the binary classification analysis, thresholds were estimated from all nuclei outside cluster 9 by calculating the per-cell mean expression for each module, serving as the non-biosynthetic background. A nucleus within cluster 9 was classified as "on" if its mean module expression exceeded its corresponding module-specific threshold. Fisher's exact test was applied to measure the strength of association between the two module binary classification tables.

Quantification and Statistical Analysis

All statistical analyses were performed on measurements from distinct biological samples not technical repeat injections. For transient expression experiments in *N. benthamiana*, leaf-paired designs were used to reduce leaf and plant specific pathway variance by infiltrating compared conditions into opposite sides of the same leaf. In such cases, paired data points are indicated in the corresponding plots and paired t-tests were used. For unpaired comparisons, biological replicates were distributed across plants and leaves of different relative ages to reduce plant-to-plant and leaf-age effects. Bar heights depict the mean and error bars indicate the standard deviation. For comparisons between two conditions without leaf-paired designs, two-sided Student's t-tests were used unless otherwise noted. For calculations of the ratios of two metabolites, the t-test was performed on log transformed data (lognormal distribution model).

Homology searches for LABF-1 and LABF-2

The amino acid sequences of LABF-1 and LABF-2 were searched against the NCBI non-redundant protein database (June 2026) using PSI-BLAST with default parameters. For PSI-BLAST queries to *P. tetrastichus* and *D. obscurum* proteomes, see **Supplementary Code**.

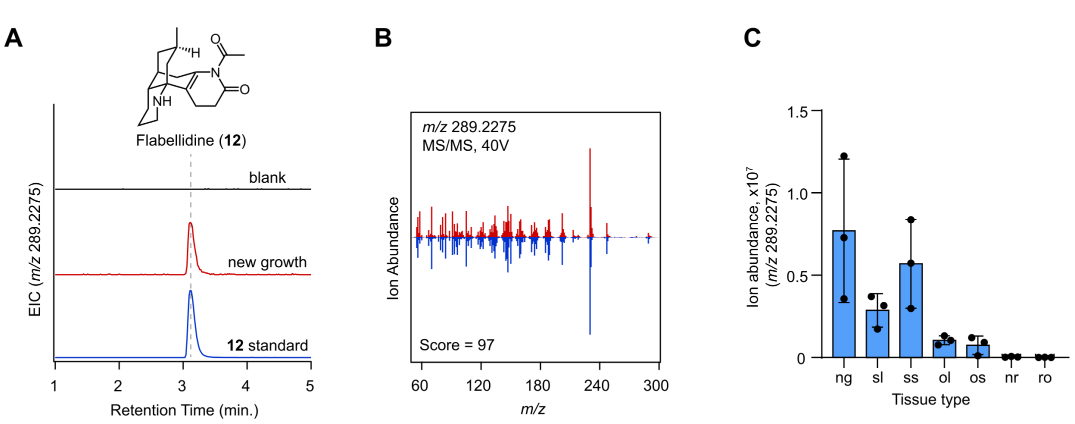

**Fig. S1. Validation of flabellidine (12) in *D. obscurum* and quantification across tissues.**
**(A)** Extracted ion chromatograms (EICs) from LC-MS analysis of methanolic extracts of *D. obscurum* new growth tissue (red) for flabellidine (**12**) ([M+H]+ = *m/z* 289.2275) compared to an authentic standard (blue). **(B)** Comparison of $MS^2$ spectra between putative flabellidine (**12**) from *D. obscurum* (red) and a standard (blue). **(C)** Quantification of flabellidine (**12**) accumulation in the following *D. obscurum* tissues harvested for RNA-sequencing: new growth leaves (ng), sub-new growth leaves, (sl), sub-growth stems (ss), old leaves (ol), old stems (os), new rhizome (nr), and roots (ro).

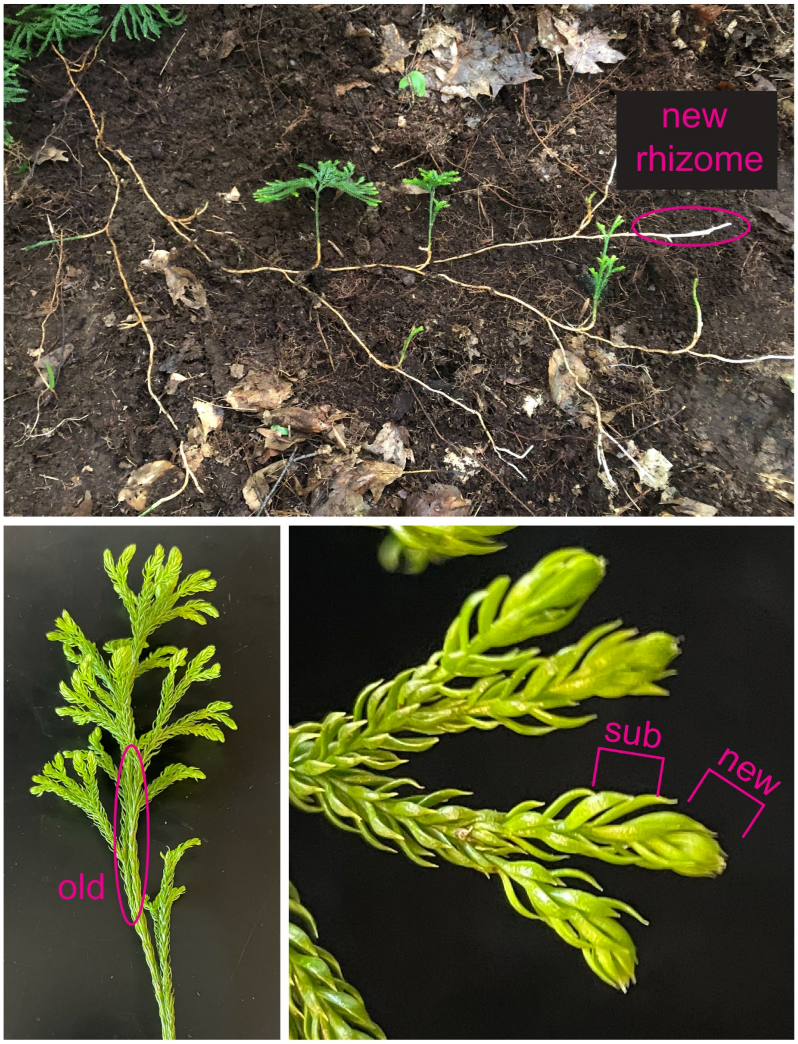

**Fig. S2. *D. obscurum* tissue types isolated for $D_2O$ labeling and RNA-sequencing.** Tissue types isolated for RNA-sequencing and metabolomic analysis are annotated in pink. The new rhizome consisted of white-colored, putative actively-growing tissue at the end of the rhizome. Roots (not shown) were removed from the rhizome.

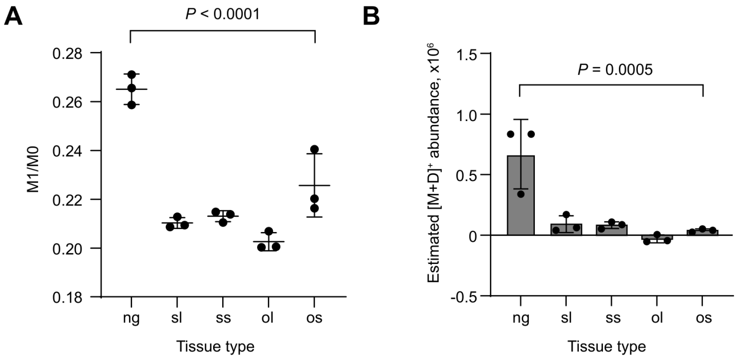

**Fig. S3. Deuterium enrichment of flabellidine (12) after $D_2O$ labeling of *D. obscurum*.** **(A)** Quantification of flabellidine (**12**) isotopologues following incubation of a *D. obscurum* cutting with 20% $D_2O$, shown as the ratio of the M+1 flabellidine isotopologue (M1 = *m/z* 290.2338) compared to the monoisotopic mass (M0 = *m/z* 289.2275). New growth = ng, sub-new growth leaves = sl, sub-new growth stems = ss, old leaves = ol, old stems = os, new rhizome = nr, and roots = ro. $P < 0.0001$ using one way ANOVA of log(M1/M0) values across all tissues. **(B)** Isotopologue enrichment visualized by estimating the total M+D isotopologue abundance. This was done by subtracting the expected M1 isotopologue abundance from the M1 peak assuming a natural isotope abundance of $^{13}C$ and $^{2}H$ in relation to M0 abundance. $P = 0.0005$ using one way ANOVA across all tissues (N=3).

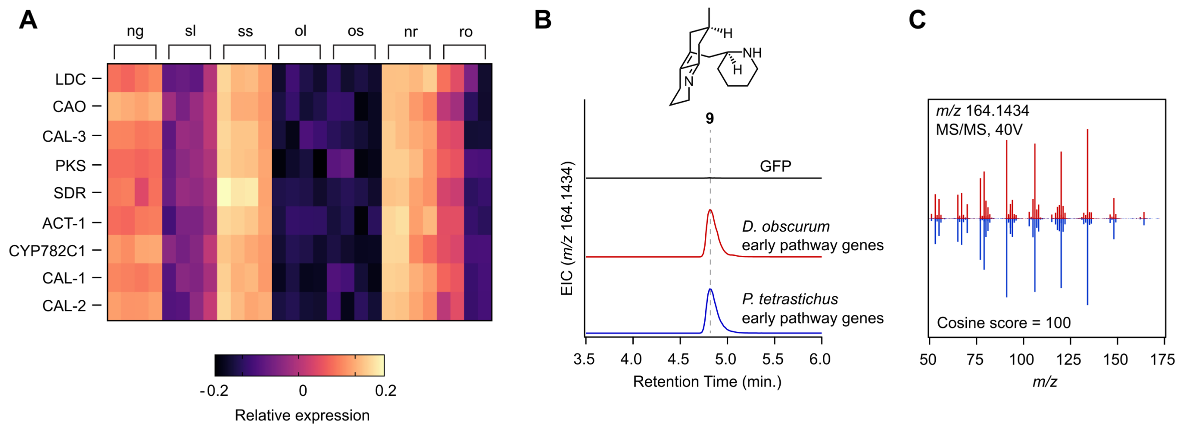

**Fig. S4. Identification and functional characterization of *D. obscurum* orthologs for previously identified Lycopodium alkaloid biosynthetic genes.**
**(A)** Tissue specific relative expression ($\log_2$ of TPM followed by centering to mean and normalizing to vector unit length) of *D. obscurum* orthologs to the previously characterized "early pathway" (**9**-producing) biosynthetic genes from *P. tetrastichus* (*22*, *23*). New growth = ng, sub-new growth leaves = sl, sub-new growth stems = ss, old leaves = ol, old stems = os, new rhizome = nr, and roots = ro. **(B)** Extracted ion chromatograms (EICs) assessing **9** production during transient expression of early pathway biosynthetic genes (from LDC through CAL1/CAL2) in *N. benthamiana*. Shown are EICs for transiently expressed *D. obscurum* early pathway gene orthologs (red) compared to transiently expressed orthologs from P. tetrastichus (blue) **(C)** Comparison of $MS^2$ spectra for **9** produced by early pathway orthologs from *D. obscurum* (red) and *P. tetrastichus* (blue).

| Gene Name | Pfam | Do*LDC-1 r* | Pt*LDC-1 r* | % identity |
|---|---|---|---|---|
| CAL-4 | Carbonic anhydrase | 0.98 | 0.97 | 82.0 |
| LABF-2 | --- | 0.99 | 0.97 | 56.8 |
| --- | Transferase | 0.99 | 0.95 | 65.9 |
| --- | Alcohol dehydrogenase | 0.99 | 0.95 | 63.8 |
| --- | Antibiotic biosynthesis monooxygenase | 0.99 | 0.97 | 33.6 |
| ACT-2 | Transferase | 0.99 | 0.98 | 69.5 |
| LABF-1 | Antibiotic biosynthesis monooxygenase | 0.99 | 0.97 | 66.5 |
| ABH-2 | Alpha/Beta hydrolase | 0.99 | 0.98 | 67.1 |
| --- | Carbonic anhydrase | 0.99 | 0.94 | 75.8 |
| CYP7032A12 | P450 | 0.98 | 0.92 | 75.3 |
| --- | Transferase | 0.98 | 0.96 | 80.8 |
| --- | 2OG-Fe(II) oxygenase | 0.98 | 0.82 | 55.6 |
| 2OGD-4 | 2OG-Fe(II) oxygenase | 0.98 | 0.98 | 68.7 |
| --- | Antibiotic biosynthesis monooxygenase | 0.98 | 0.95 | 64.9 |
| ABCC-1 | ABC transporter | 0.98 | 0.97 | 85.2 |
| --- | Alpha/Beta hydrolase | 0.98 | 0.86 | 67.7 |
| MDR-2 | Alcohol dehydrogenase | 0.98 | 0.95 | 74.2 |
| MDR-1 | Alcohol dehydrogenase | 0.97 | 0.93 | 85.5 |
| NPF8.1 | POT family peptide transporter | 0.96 | 0.93 | 71.0 |
| --- | Molybdenum cofactor sulfurase family | 0.93 | 0.87 | 72.0 |
| --- | Glycosyl hydrolase | 0.91 | 0.92 | 78.9 |
| --- | Amino acid transporter | 0.9 | 0.95 | 76.6 |

**Fig. S5. Previously uncharacterized gene candidates included in the batch of 34 well-conserved and well-correlated genes.**

The listed 22 gene candidates were selected based on having high homology between putative orthologs from *D. obscurum* and *P. tetrastichus* (represented here by percent identity) and strong co-expression with Lycopodium alkaloid biosynthetic genes in both species, represented here via Pearson's correlation to LDC ortholog expression.

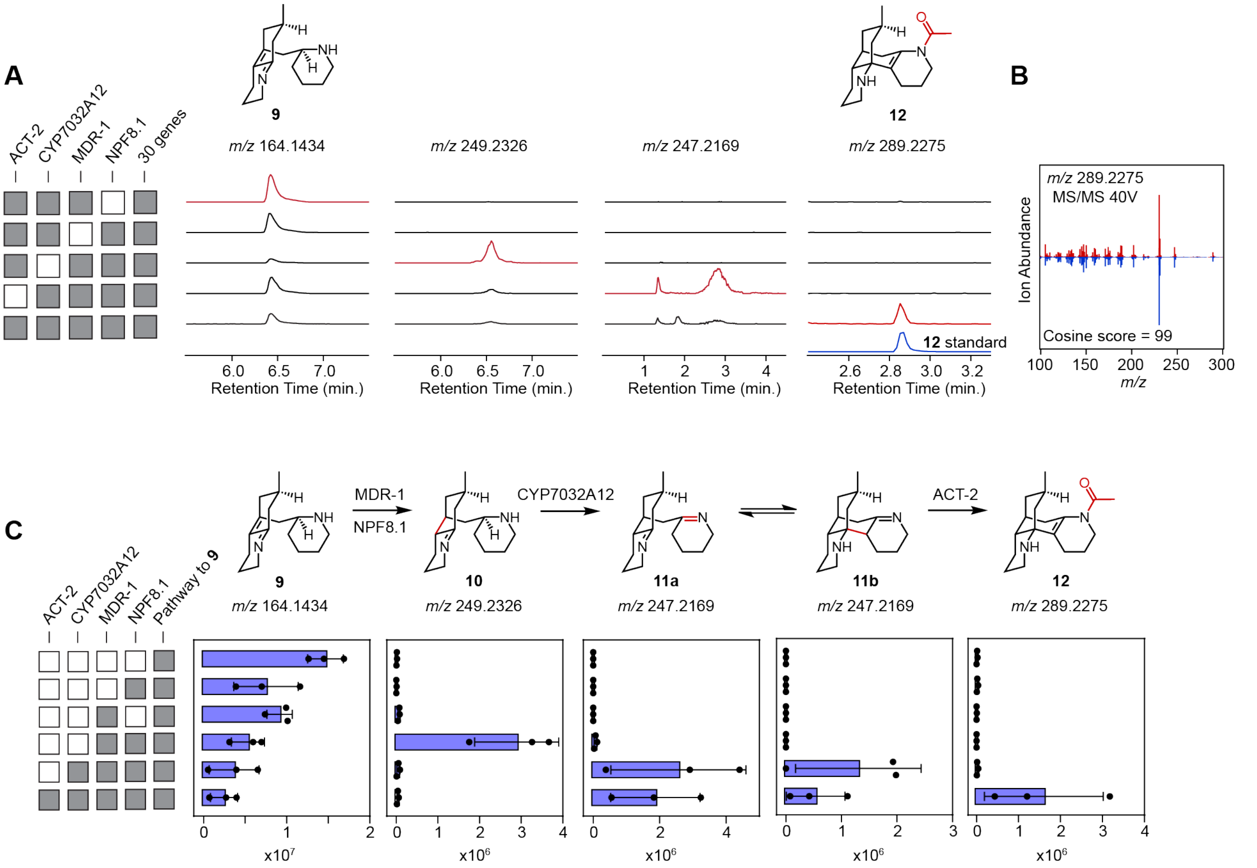

**Fig. S6. LC-MS analysis of *N. benthamiana* extracts expressing different combinations of genes required for flabellidine (12) biosynthesis**

**(A)** Extracted ion chromatograms (EICs) for putative biosynthetic intermediates found in extracts from *N. benthamiana* transient expression when individual genes from the batch of 34 are dropped out. **(B)** $MS^2$ fragmentation pattern of the *m/z* 289.2275 mass feature produced by the batch of 34 genes when expressed in *N. benthamiana* compared to a flabellidine (**12**) standard. **(C)** LC-MS quantification of intermediates during pathway reconstruction from **9** to **12**. Note that **11a** corresponds to the early eluting *m/z* 247.2169 peak and **11b** corresponds to the late eluting 247.2169 peak.

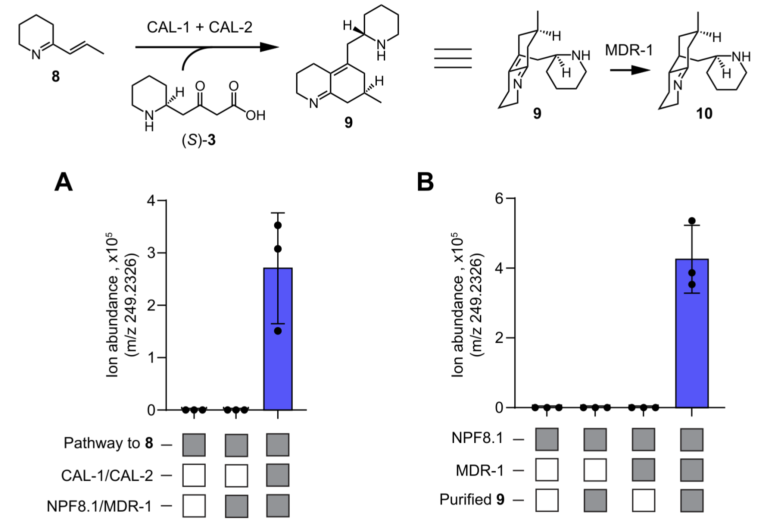

**Fig. S7. Substrate specificity of MDR-1.**
**(A)** Quantification of **10** abundance (*m/z* 249.2326) in extracts from *N. benthamiana* transiently expressing gene combinations that produce **9** (the product of CAL-1/CAL-2) or lack the capacity to produce **9**. **(B)** Quantification of **10** abundance in extracts from *N. benthamiana* expressing gene combinations of NPF8.1 and MDR-1 with co-infiltration of HPLC-isolated **9** into leaves as substrate.

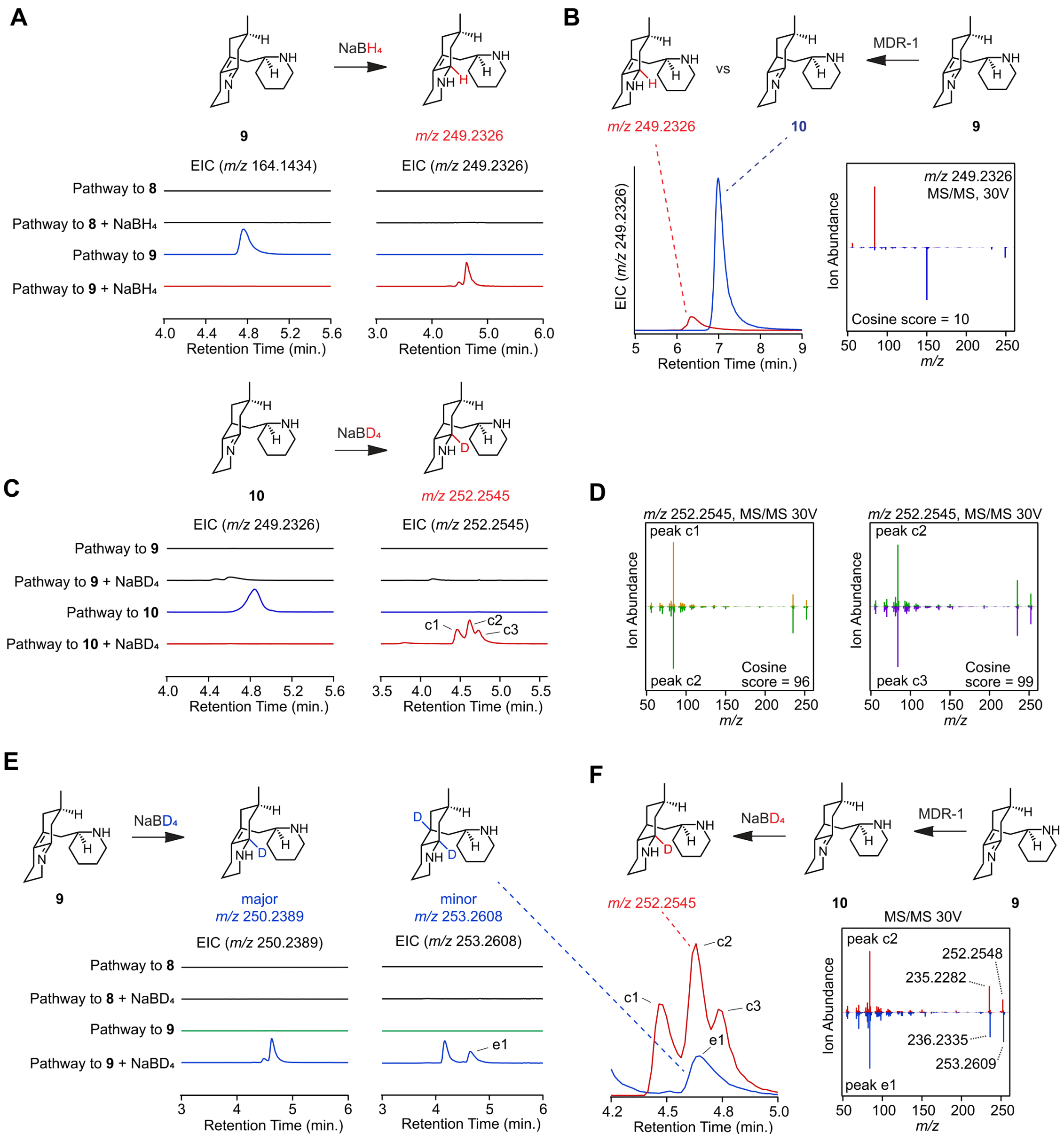

**Fig. S8. Support for the structure of 10 from chemical derivatization.**
**(A)** Extracted ion chromatograms (EICs) from extracts of *N. benthamiana* transiently expressing the pathway to **8** or **9** following reduction with sodium borohydride ($NaBH_4$) **(B)** Comparisons of the retention times and MS$^2$ fragmentation patterns of the products of $NaBH_4$ treatment on extracts of the pathway to **9** (red) compared to **10** produced enzymatically from MDR-1 when expressed on top of the pathway to **9** in *N. benthamiana* (blue) confirm that the $NaBH_4$ reduced version of **9** is distinct from enzymatically produced **10** **(C)** EICs from extracts of *N. benthamiana* transiently expressing the pathway to **9** or **10** following reduction with $NaBD_4$ **(D)** MS$^2$ fragmentation patterns of the *m/z* 252.2545 mass features shown in panel (C) (yellow = early retention time, green = middle retention time, purple = late retention time). Similar fragmentation patterns suggest that these peaks are likely diastereomers. **(E)** EICs from extracts of *N. benthamiana* transiently expressing the pathway to **8** or **9** following reduction with $NaBD_4$ **(F)** Comparisons of the retention

times and $MS^2$ fragmentation patterns of the product of $NaBD_4$ treatment on extracts of the pathway to **9** (blue, *m/z* 253.2608) compared to the product of $NaBD_4$ treatment on extracts of the pathway to **10** (red, *m/z* 252.2545). The fragmentation patterns display a nearly identical fragmentation pattern other than the large fragments that differ by a mass equal to the difference in deuterium incorporations (cosine score considering full spectra = 75, considering just below *m/z* 200 cosine score = 94), suggesting the product of MDR-1 (**10**) retains the phlegmarane scaffold of **9**.

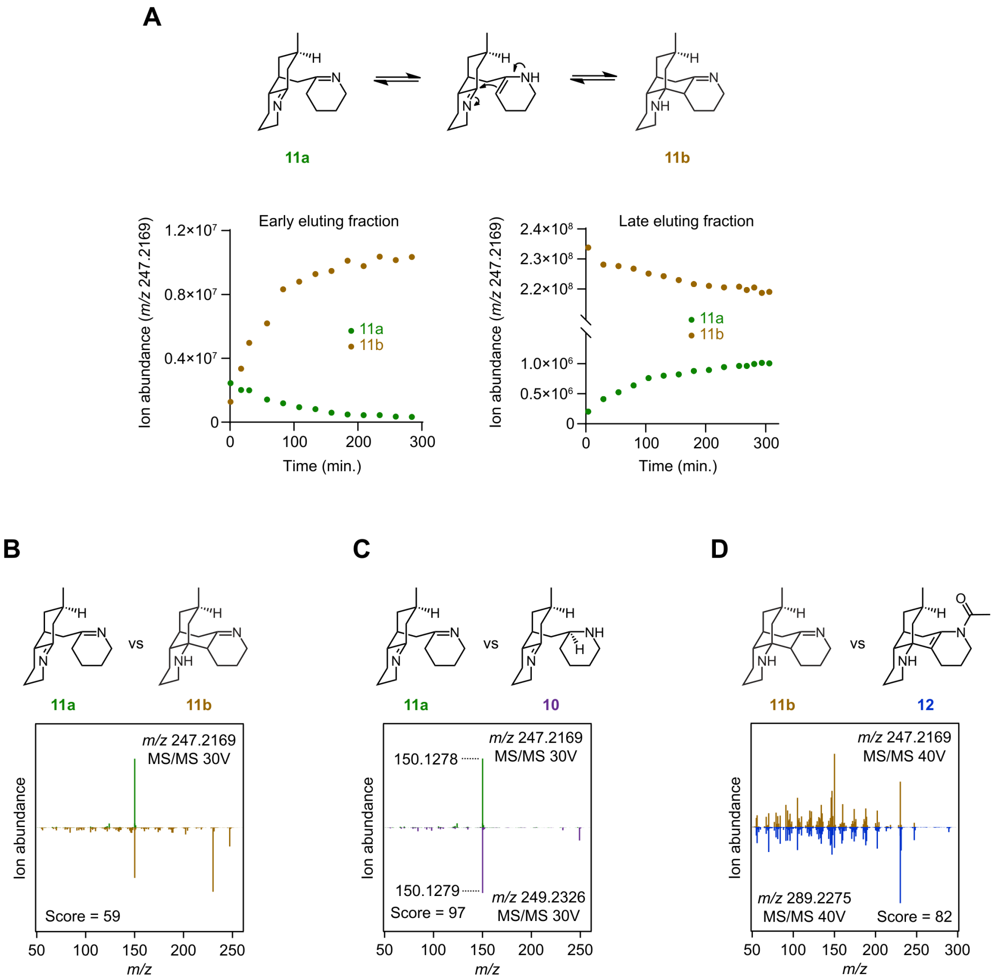

**Fig. S9. Equilibration and MS$^2$ fragmentation patterns of 11a and 11b**

**(A)** Following HPLC fractionation of extractions from *N. benthamiana* expressing the pathway to **11**, samples were repeatedly analyzed by LC-MS over time. Shown are the quantifications of **11a** and **11b** over time. Note that the initially isolated species decreases while the isomeric species increases. **(B)** MS$^2$ fragmentation patterns of **11a** (green) and **11b** (bronze) **(C)** MS$^2$ fragmentation pattern of **11a** (green) compared to **10** (purple) **(D)** MS$^2$ fragmentation pattern of **11b** (bronze) compared to **12** (blue).

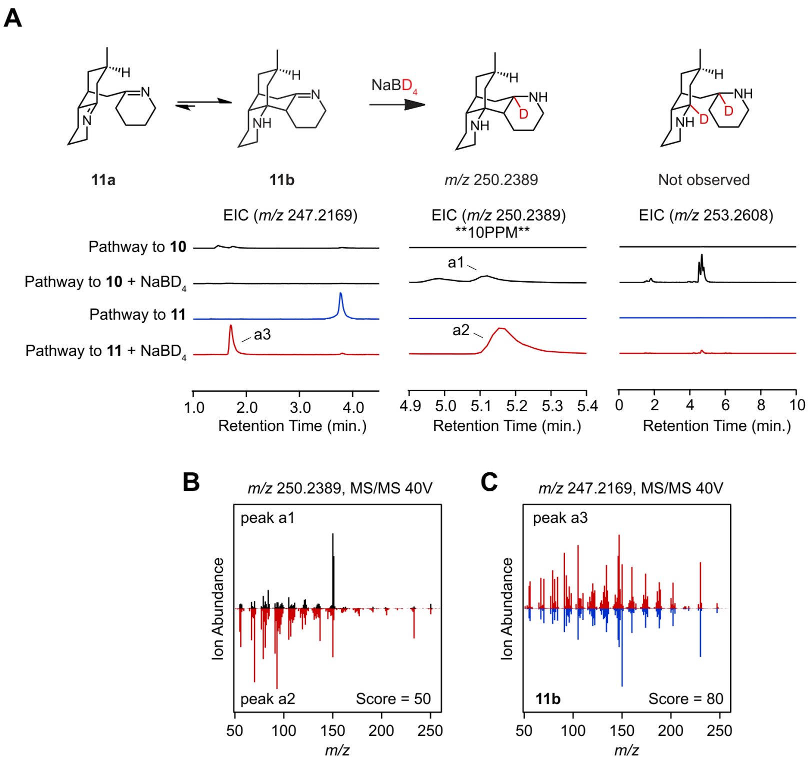

**Fig. S10. LC-MS analysis of extracts of 11b treated with $NaBD_4$**

**(A)** Expressing the pathway to **11** in *N. benthamiana* and treating this extract with $NaBD_4$ yields a unique *m/z* 250.2389 mass feature (a2; EIC at 10ppm to avoid $^{13}C$ isotopologue of **10**). Comparison to the pathway to **10** treated with $NaBD_4$ suggests that a2 is a singly reduced form of **11b**. **(B)** Two closely eluting *m/z* 250.2389 mass features were produced when pathways to **10** or **11** were treated with $NaBD_4$ (panel A). $MS^2$ fragmentation was performed to further confirm they are distinct molecules (black = peak a1, red = peak a2). **(C)** An unknown *m/z* 247.2169 mass feature (peak a3, panel A) arises after treatment of extract to pathway **11** with $NaBD_4$. Peak a3 (red) shares a very similar fragmentation pattern to enzymatically-produced **11b** (blue). No doubly reduced *m/z* 253.2608 mass features are observed from treatment with $NaBD_4$ to the pathway to **11** extract, suggesting **11b** only has a single double bond that can be reduced.

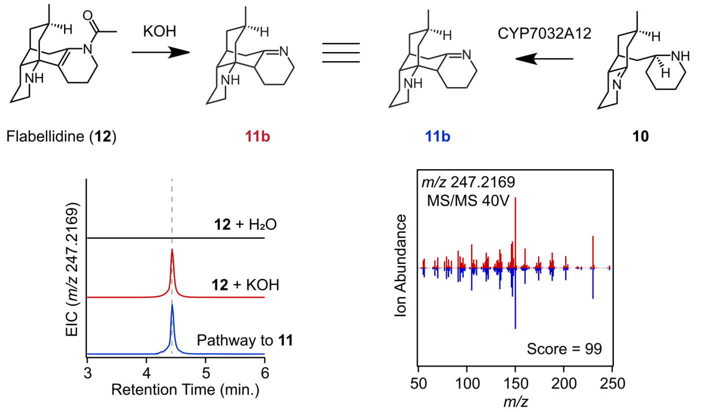

**Fig. S11. Base-catalyzed hydrolysis of flabellidine (12).**
LC-MS analysis following base-catalyzed hydrolysis of flabellidine (**12**) demonstrates the formation of a mass feature (*m/z* 247.2169, red) with a matching retention time and $MS^2$ fragmentation pattern to **11b** (blue).

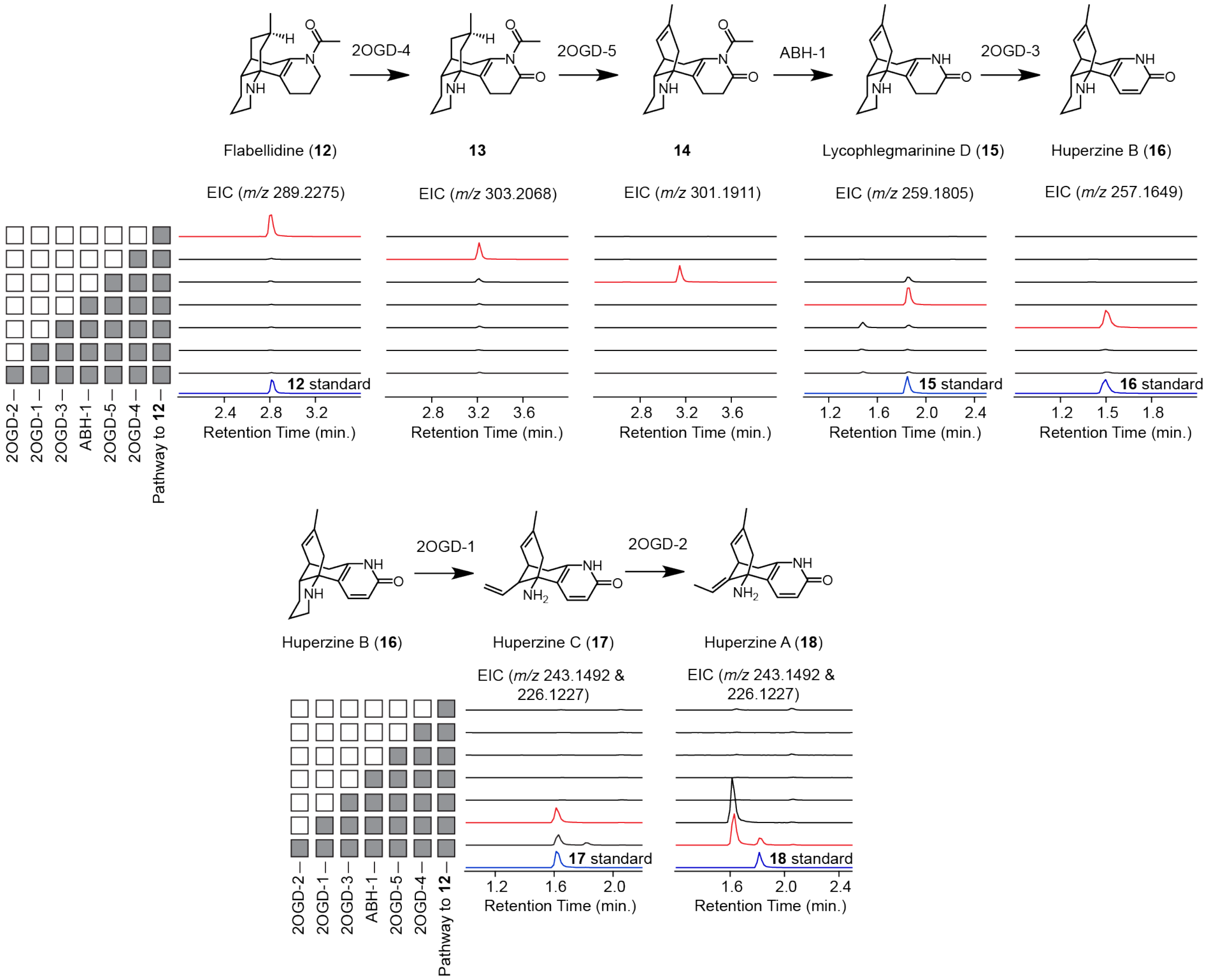

**Fig. S12. Step-by-step reconstruction of the full huperzine A (18) pathway**.
LC-MS analysis of extracts from *N. benthamiana* expressing combinations of genes on route to huperzine A (**18**). Shown here is each step from biosynthetically-produced flabellidine (**12**).

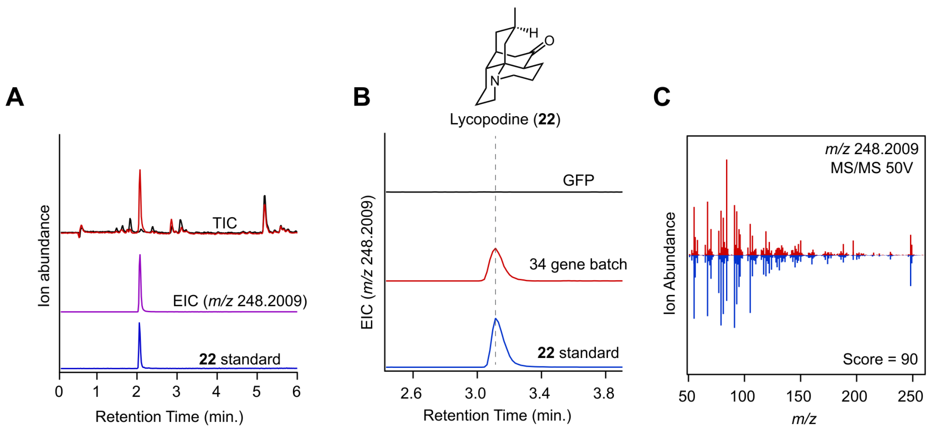

**Fig. S13. Detection of lycopodine (22) in *D. obscurum* and production in *N. benthamiana.***
**(A)** LC-MS analysis of *D. obscurum* new-growth tissue. The top trace shows the total ion chromatogram (TIC) from *D. obscurum* new growth (red) overlaid with a solvent blank (black). The middle trace (purple) shows the extracted ion chromatogram (EIC) for lycopodine (**22**) at *m/z* 248.2009. The lower trace (blue) is a lycopodine (**22**) standard. **(B)** Heterologous expression of the 34-gene candidate set in *N. benthamiana* produces a peak that co-elutes with lycopodine (**22**). EICs (*m/z* 248.2009) are shown for a GFP negative control (black), the 34-gene expression condition (red), and a lycopodine (**22**) standard (blue). **(C)** $MS^2$ fragmentation of the product from *N. benthamiana* (red) compared to the lycopodine (**22**) standard (blue).

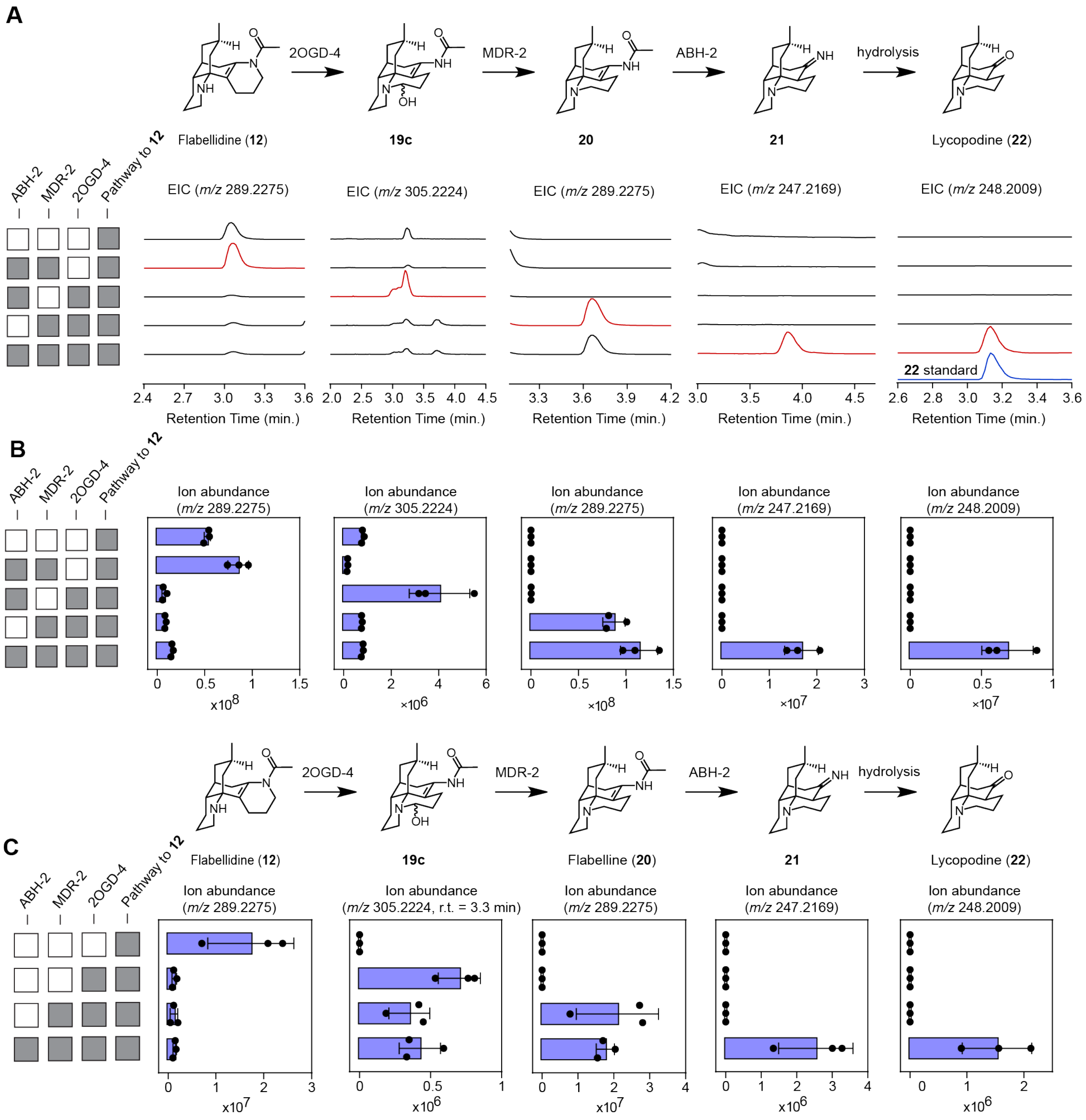

**Fig. S14. Reconstruction of lycopodine (22) biosynthesis in *N. benthamiana*.**
**(A)** LC-MS analysis of *N. benthamiana* extracts expressing individual dropouts from the batch of 34 that prevent lycopodine (**22**) accumulation along with putative intermediates identified through LC-MS with HILIC chromatography. **(B)** Quantification of EIC peak areas from the chromatograms shown in panel (A). For the **19c** panel, all extracted ions between approximately 2.9 and 3.4 min were summed for *m/z* 305.2224 **(C)** Quantification of pathway intermediates during stepwise lycopodine (**22**) pathway reconstruction from LC-MS analysis with C18 chromatography.

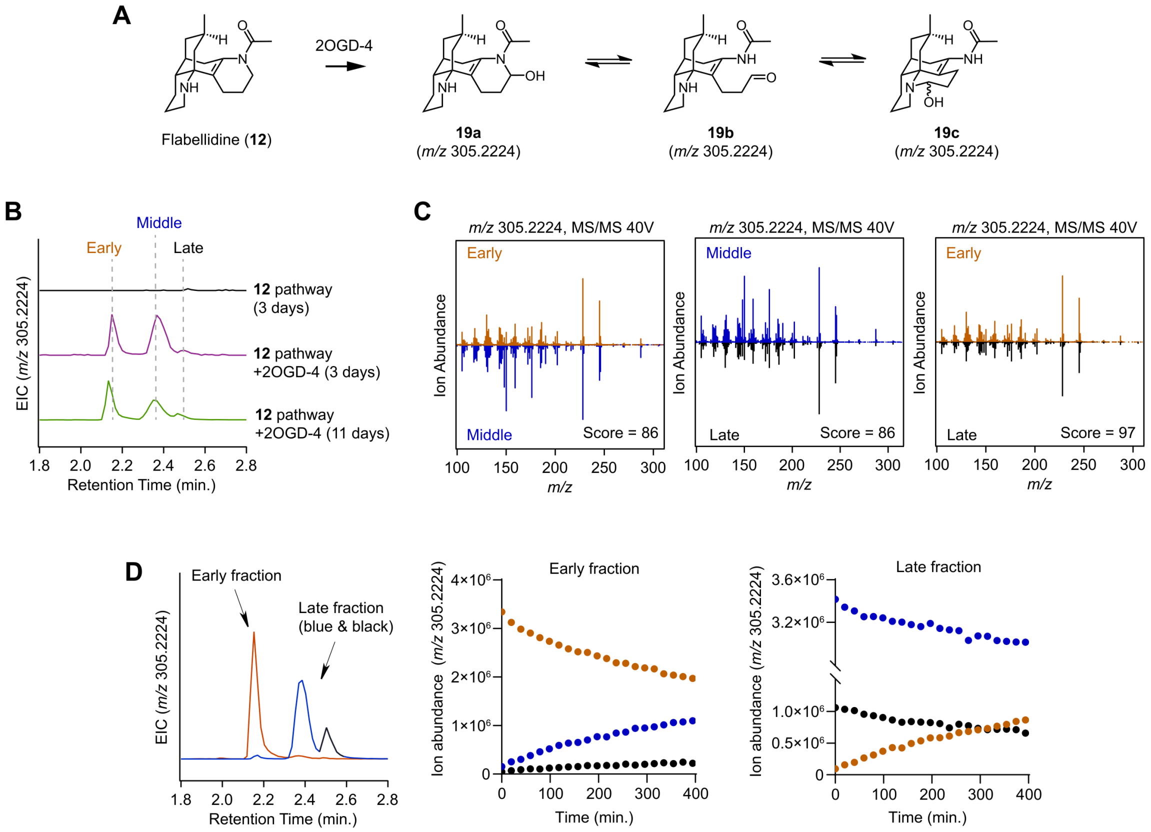

**Fig. S15. Equilibration of 19 isomers.**

**(A)** Proposed equilibrium of **19** isomers following oxidation of flabellidine (**12**) by 2OGD-4. **(B)** Expression of the flabellidine (**12**) pathway (black) vs flabellidine pathway +2OGD-4 for three days (purple) generates three distinct *m/z* 305.2224 peaks, which are retained when the pathway is expressed for 11 days (green) in preparation for HPLC fractionation. **(C)** $MS^2$ analysis of the early eluting (orange) and late eluting (black) isomers from panel (B) show that these molecules have identical fragmentation patterns, suggesting they are likely diastereomers. The middle eluting isomer (blue, from B) is similar, but distinct from the early and late-eluting molecules. **(D)** HPLC fractionation of the *m/z* 305.2224 mass features enables partial separation of *m/z* 305.2224 fractions, with isolation of an early fraction containing mostly the early isomer (orange) and the isolation of a later fraction containing mostly the middle (blue) and late (black) isomers. Repeated injections of these fractions into LC-MS shows time dependent decreases in the initially purified species with a corresponding increase in the other isomers.

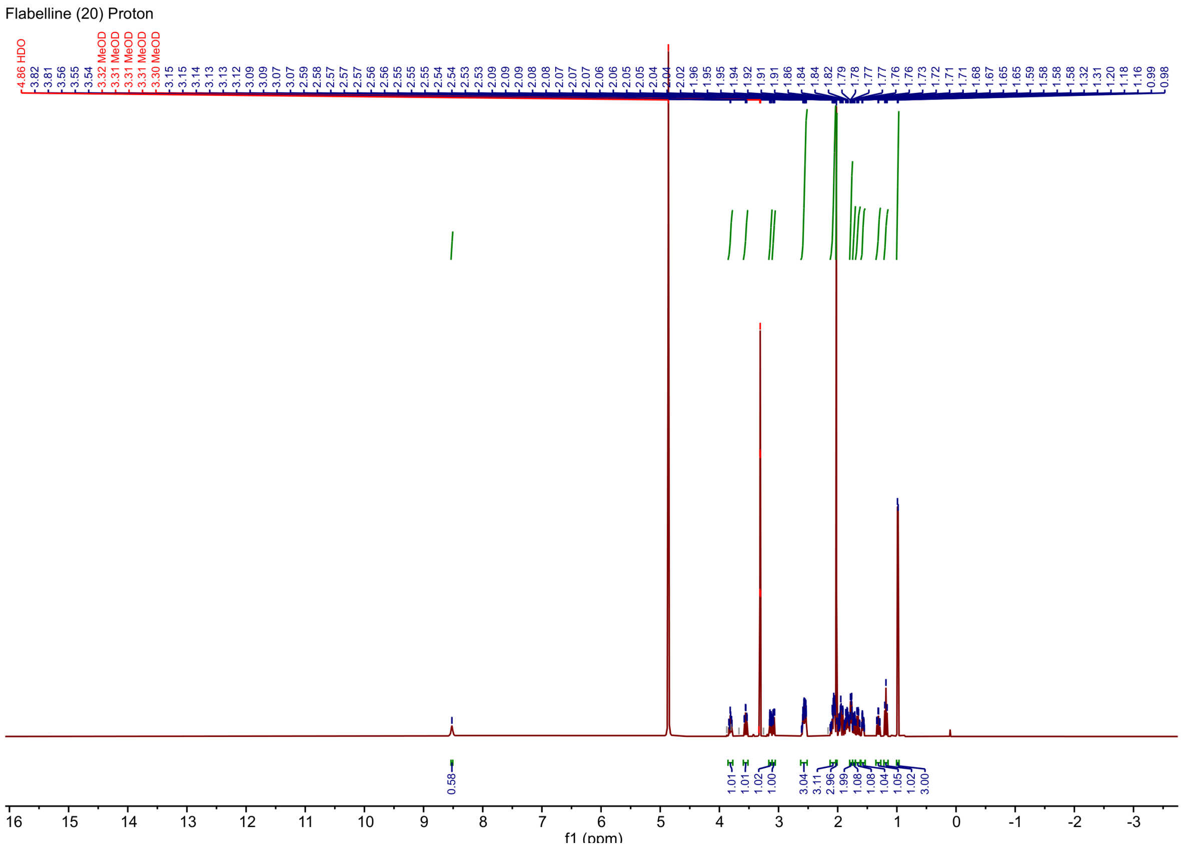

**Fig. S16. $^1$H NMR spectrum of flabelline (20)**

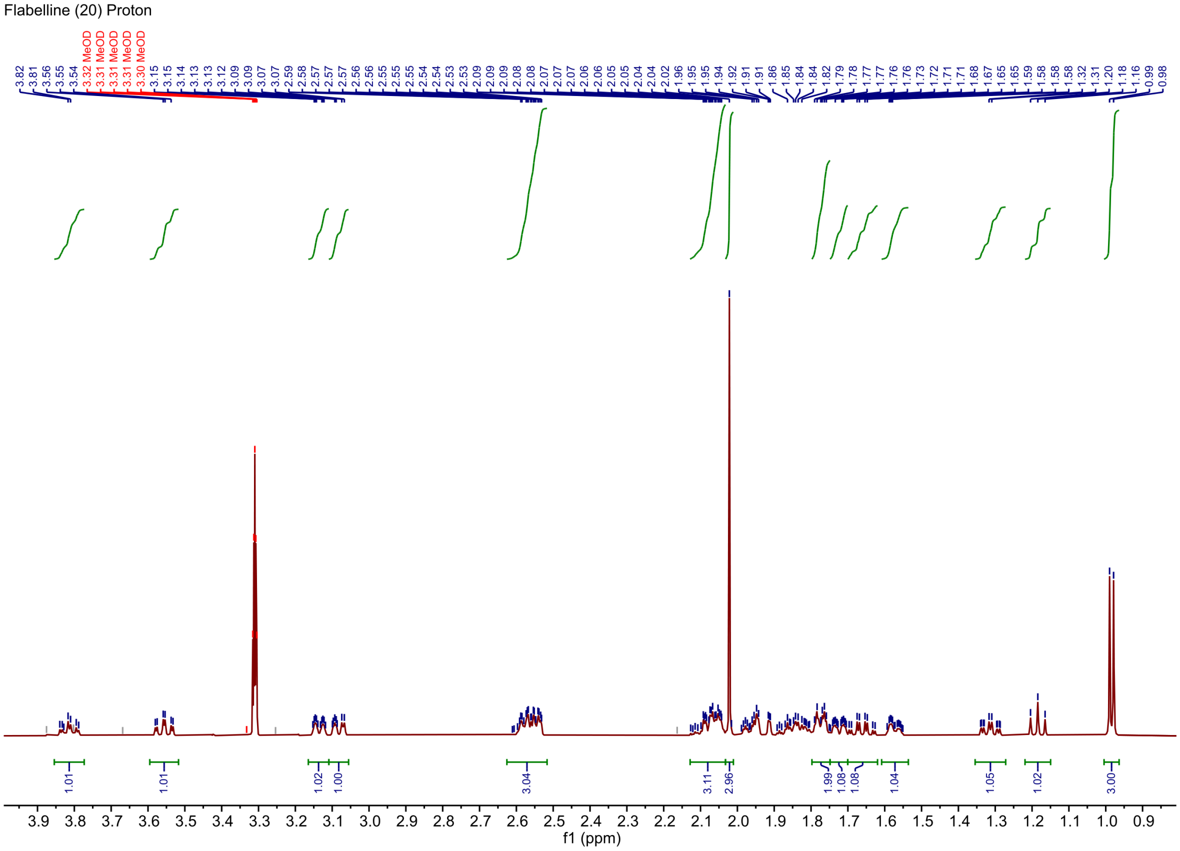

**Fig. S17. Zoomed in $^{1}$H NMR spectrum of flabelline (20)**

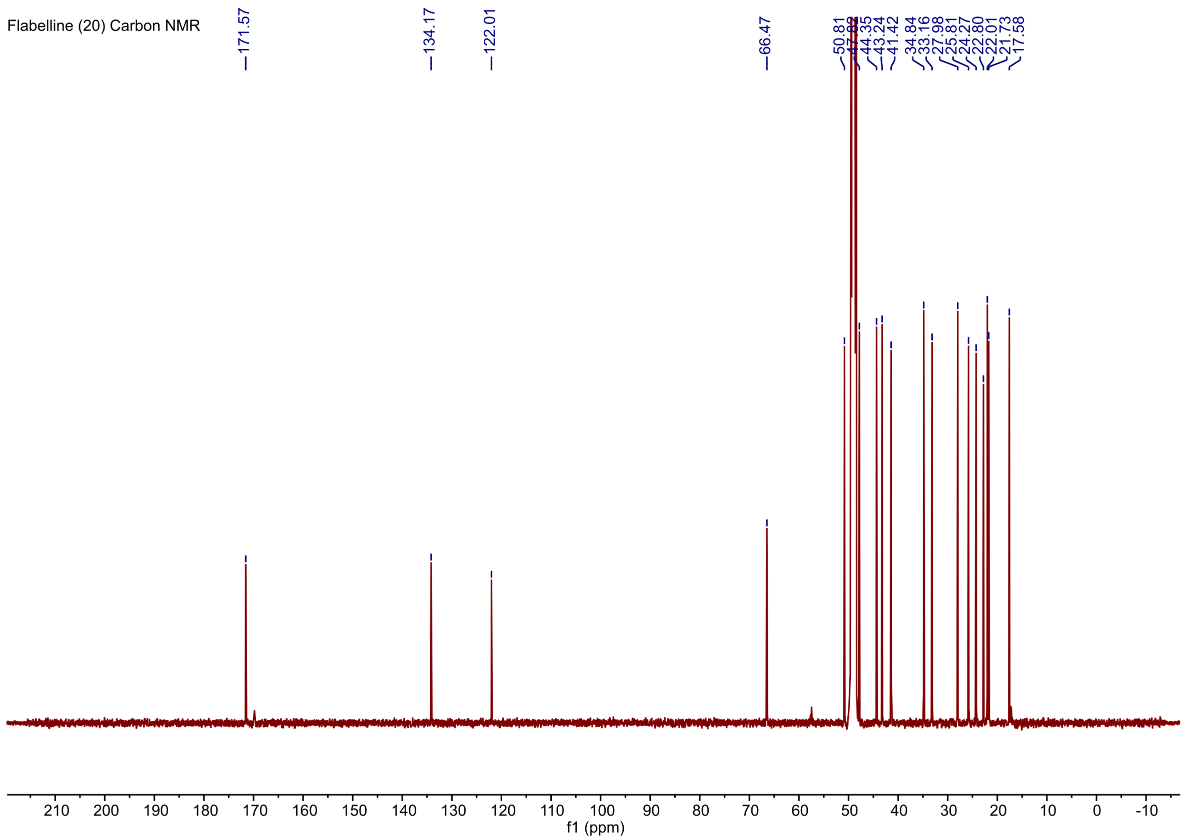

**Fig. S18. Carbon NMR spectrum of flabelline (20)**

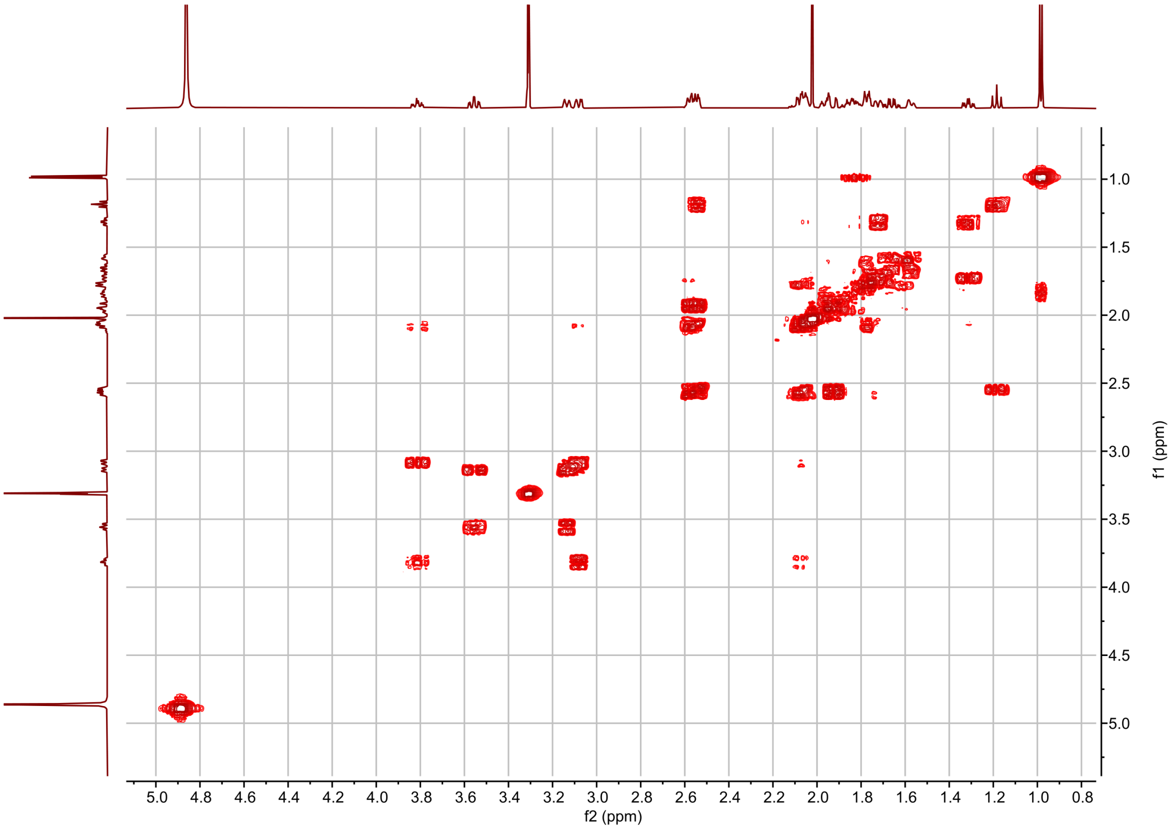

**Fig. S19. COSY spectrum of flabelline (20)**

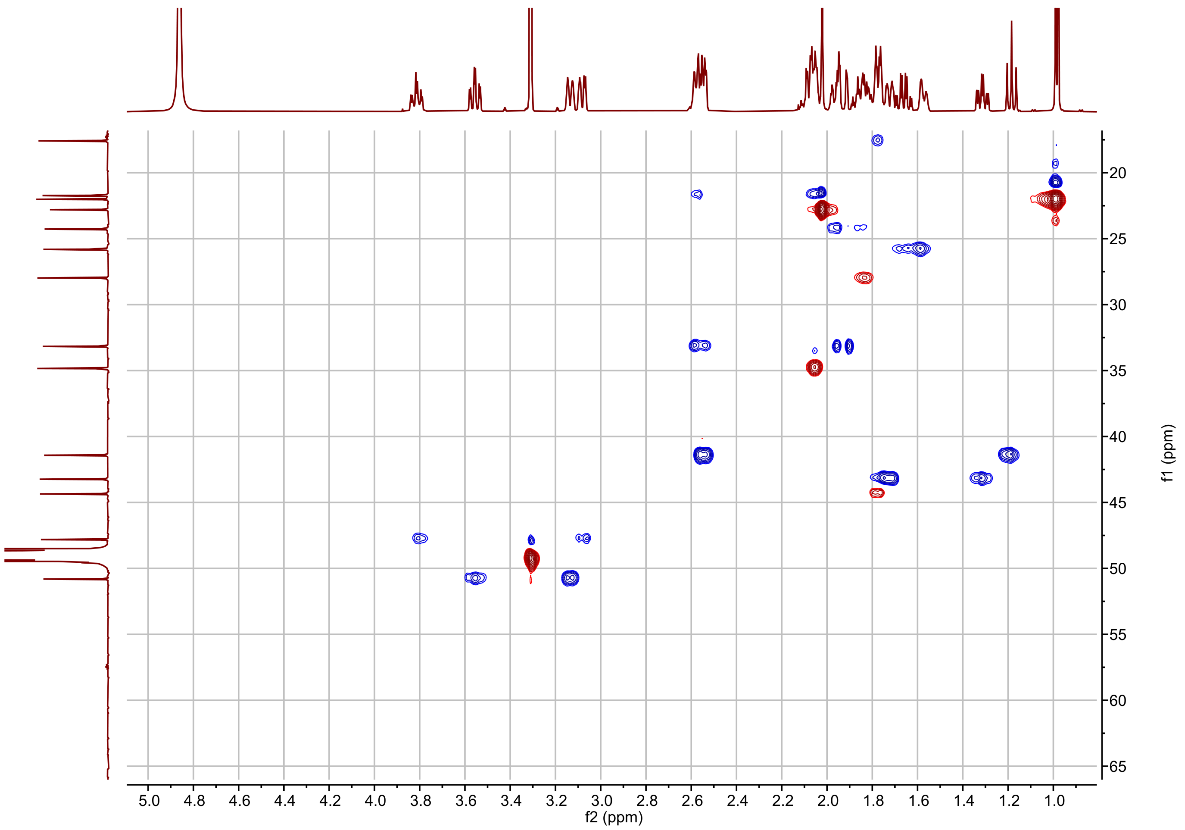

**Fig. S20. HSQC spectrum of flabelline (20)**

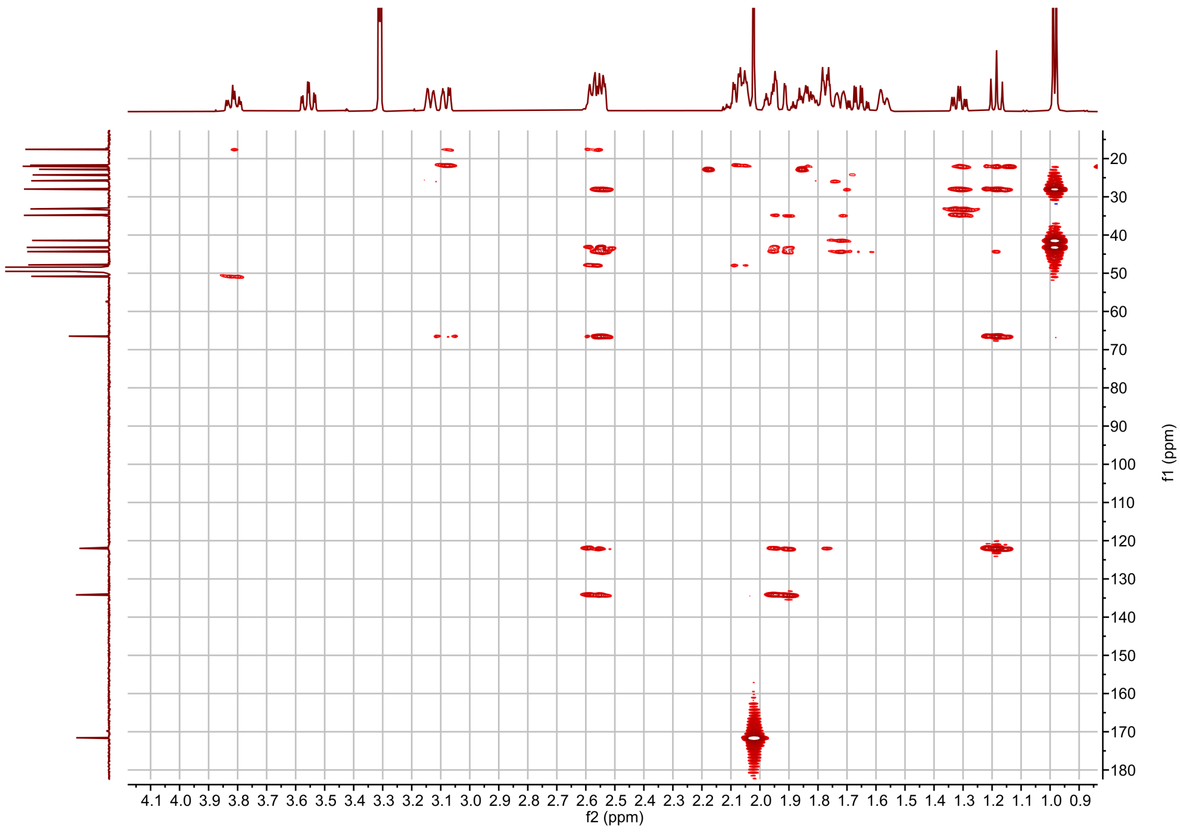

**Fig. S21. HMBC spectrum of flabelline (20)**

A

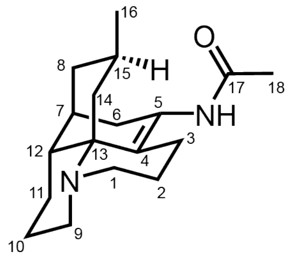

Flabelline (**20**)

B

| Carbon | | C δ (ppm) | H1 δ (ppm) | H2 δ (ppm) |
|---|---|---|---|---|
| **1** | CH2 | 47.82 | 3.08 (1H, m) | 3.81 (1H, td, J = 13.3, 4.0 Hz) |
| **2** | CH2 | 17.58 | 1.78 (1H, m) | 2.09 (1H, m) |
| **3** | CH2 | 21.73 | 2.04 (1H, m) | 2.56 (1H, m) |
| **4** | C | 125.22 | | |
| **5** | C | 137.39 | | |
| **6** | CH2 | 33.17 | 1.91 (1H, m) | 2.58 (1H, m) |
| **7** | CH | 34.84 | 2.05 (1H, m) | |
| **8** | CH2 | 43.24 | 1.31 (1H, m) | 1.72 (1H, m) |
| **9** | CH2 | 50.81 | 3.14 (1H, ddt, J = 12.5, 4.2, 2.3 Hz) | 3.56 (1H, td, J = 13.0, 3.1 Hz) |
| **10** | CH2 | 24.27 | 1.95 (1H, m) | 1.87 (1H, m) |
| **11** | CH2 | 25.81 | 1.66 (1H, qd, J = 12.7, 3.8 Hz) | 1.57 (1H, ddt, J = 14.4, 5.5, 3.5 Hz) |
| **12** | CH | 44.35 | 1.78 (1H, m) | |
| **13** | C | 69.68 | | |
| **14** | CH2 | 41.42 | 1.18 (1H, t, J = 11.9 Hz) | 2.54 (1H, m) |
| **15** | CH | 27.98 | 1.84 (1H, m) | |
| **16** | CH3 | 22.01 | 0.98 (3H, d, J = 6.6 Hz) | |
| **17** | C | 174.79 | | |
| **18** | CH3 | 22.80 | 2.02 (3H, s) | |

C

**COSY Correlations**

| H | Correlates with H |
|---|---|
| 1 | 1, 2 |
| 2 | 2 |
| 3 | 2 |
| 6 | 6 |
| 7 | 6 |
| 8 | 7, 8 |
| 9 | 9, 10 |
| 10 | 10 |
| 14 | 14 |
| 15 | 16 |

**HMBC Correlations**

| C | Correlates with H |
|---|---|
| 1 | 2, 3 |
| 2 | 1, 3 |
| 3 | 1, 2 |
| 4 | 3, 6, 12, 14 |
| 5 | 3, 6 |
| 6 | 8 |
| 7 | 6, 8, 10 |
| 8 | 6, 14, 15, 16 |
| 9 | 1, 11 |
| 10 | 11 |
| 11 | 8, 9 |
| 12 | 8, 10, 11, 14 |
| 13 | 1, 3, 9, 14 |
| 14 | 8, 15, 16 |
| 15 | 8, 14, 16 |
| 16 | 8, 14, 15 |
| 17 | 18 |

**Fig. S22. NMR assignment of flabelline (20) spectra**.

**(A)** Structure of flabelline (**20**) with atom numbering used for NMR assignments. **(B)** H and C NMR assignments for flabelline (**20**) **(C)** Observable 2D correlations. Numbering is based on the carbon numbering shown in **(A)**. In many cases, protons connected to the same carbon are magnetically distinct, but are not reported as separate hydrogens in these tables. All hydrogens are numbered based on which carbon they are connected to.

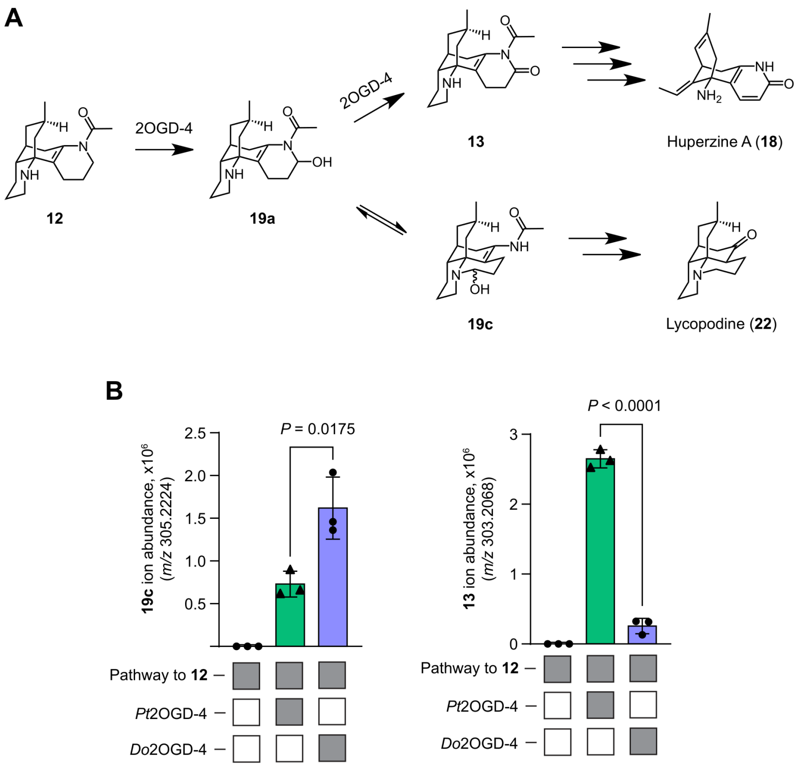

**Fig. S23. 2OGD-4 orthologs from *D. obscurum* and *P. tetrastichus* yield distinct product outcomes.**

(**A**) The branching point between lycopodine (**22**) and huperzine A (**18**) biosynthesis occurs at the step where 2OGD-4 oxidizes flabellidine (**12**). If only a single oxidation occurs, the scaffold is free to rearrange to the lycopodane scaffold (**19c**) en route to lycopodine (**22**). A double oxidation locks the structure into a product with the lycodane scaffold (**13**). **(B)** LC-MS quantification of product outcome using either the *P. tetrastichus* or *D. obscurum* 2OGD-4 on top of the pathway to flabellidine (**12**) in *N. benthamiana* transient expression. The left panel quantifies the ion abundance of **19c** and the right panel quantifies **13**. P-values are the results of unpaired Student's t-tests (N=3).

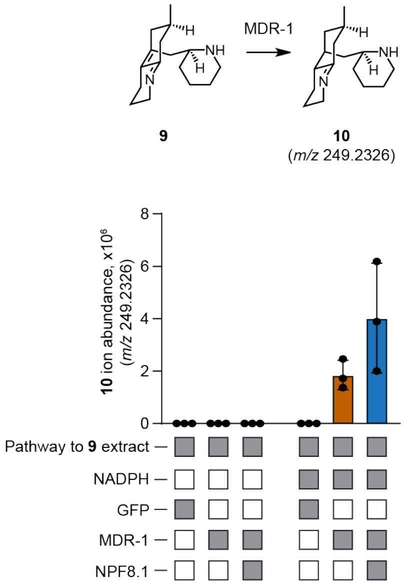

**Fig. S24. MDR-1 activity in vitro does not require NPF8.1.**
Quantification of data from Fig. 3A. LC-MS analysis of reactions with in vitro protein lysates from *N. benthamiana* expressing MDR-1 +/- NPF8.1.

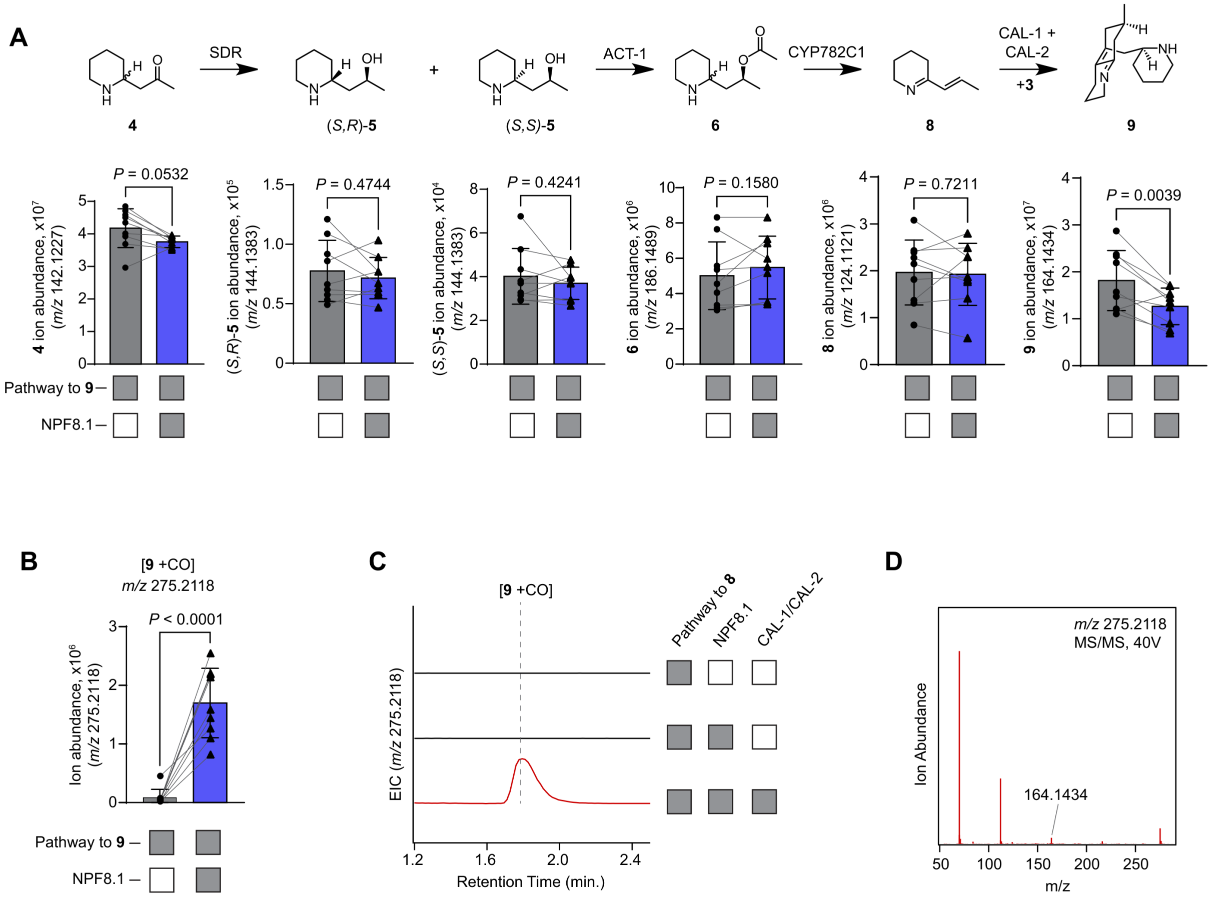

**Fig. S25. The effect of NPF8.1 expressed on top of the pathway to 9 in *N. benthamiana*.** **(A)** Quantification of pathway intermediate abundances from *N. benthamiana* transient expression of the pathway to **9** +/- NPF8.1. P-values shown are the result of paired Student t-tests (N=9). **(B)** Through XCMS analysis, an unknown mass feature (*m/z* 275.2118; tentatively 9+CO) is substantially increased with the co-expression of NPF8.1 on top of the pathway to **9**. P < 0.0001 using paired Student's t-test (N=9). **(C)** The *m/z* 275.2118 mass feature identified in **(B)** requires CAL-1/CAL-2 to accumulate, suggesting it is a downstream product of **9**. **(D)** MS$^2$ analysis of the *m/z* 275.2118 mass feature identified in **(B)** shows that it retains a small *m/z* 164.1434 fragment, the same in source fragment that forms during LC-MS ionization and detection of **9**.

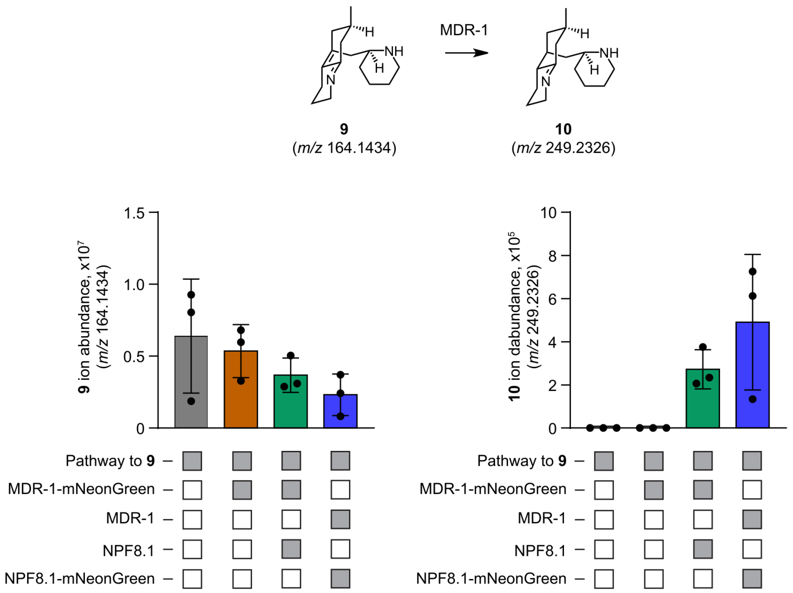

**Fig. S26. Functional validation of MDR-1 and NPF8.1 fluorophore fusions.**
LC-MS quantification of pathway intermediates from transient expression in *N. benthamiana* demonstrates that MDR-1-mNeonGreen produces **10** and still requires NPF8.1 to enable production of **10** in planta. Similarly, NPF8.1-mNeonGreen still enables MDR-1 to produce **10** in planta during pathway reconstruction.

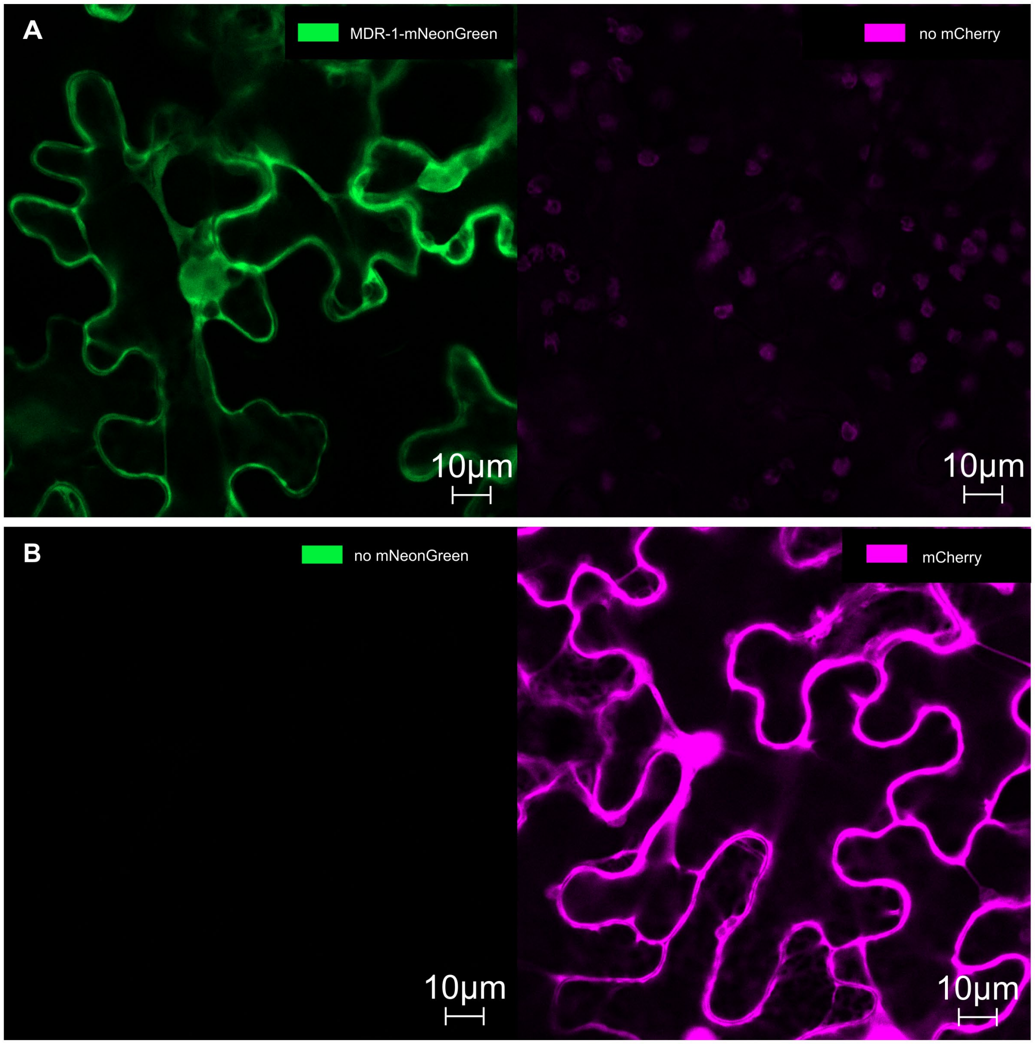

**Fig. S27. Single fluorophore controls for confocal imaging of MDR-1-mNeonGreen.**
**(A)** Control images of MDR-1-mNeonGreen in the absence of mCherry show mNeonGreen channel specificity. **(B)** Control images of mCherry in the absence of MDR-1-mNeonGreen show mCherry channel specificity.

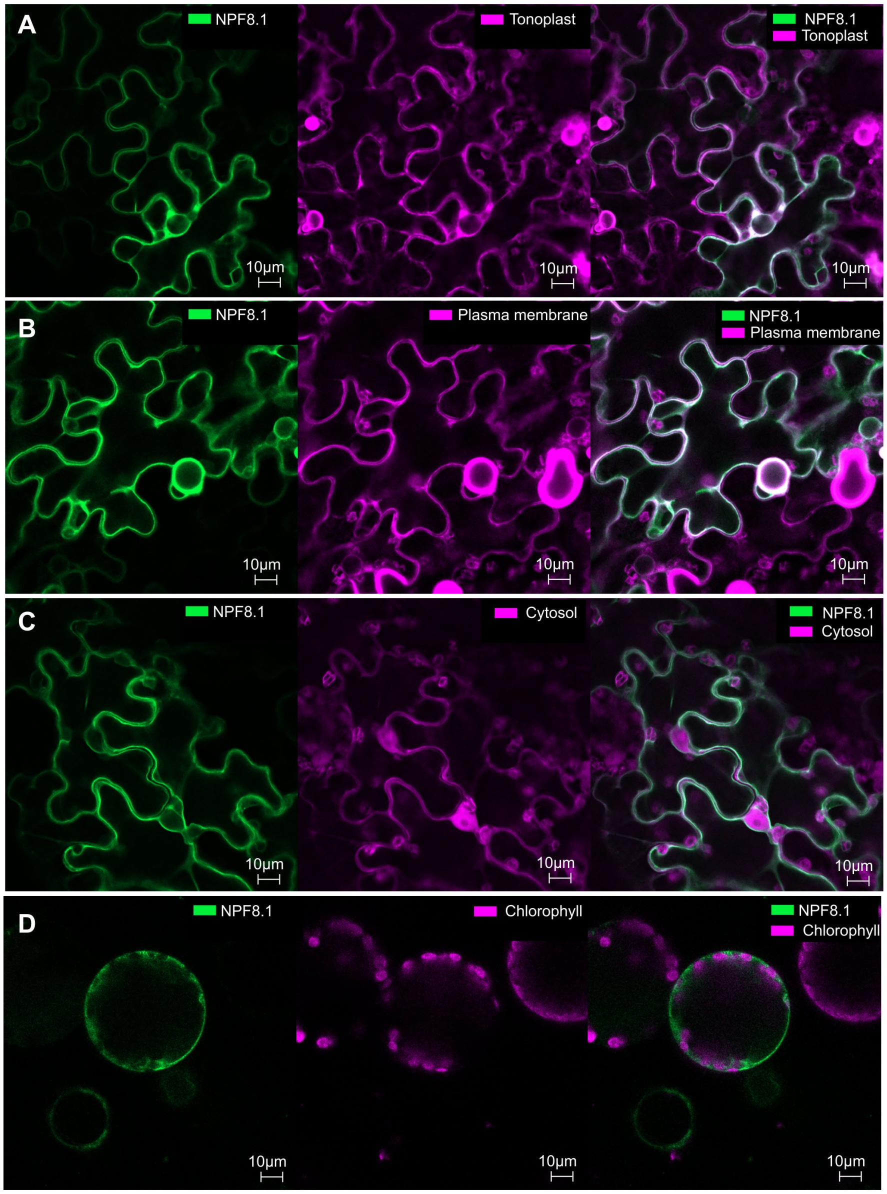

**Fig. S28. Confocal microscopy of NPF8.1-mNeonGreen with localization markers**
**(A)** NPF8.1-mNeonGreen colocalizes with a tonoplast marker γ-TIP-mCherry. **(B)** NPF8.1-mNeonGreen colocalizes with a plasma membrane marker PIP2A-mCherry. **(C)** NPF8.1-mNeonGreen visualized with cytosolic mCherry for reference does not clearly distinguish exclusive plasma membrane vs tonoplast localization. **(D)** NPF8.1-mNeonGreen visualized in protoplasts also does not clearly show signal on one particular side of chloroplasts over the other.

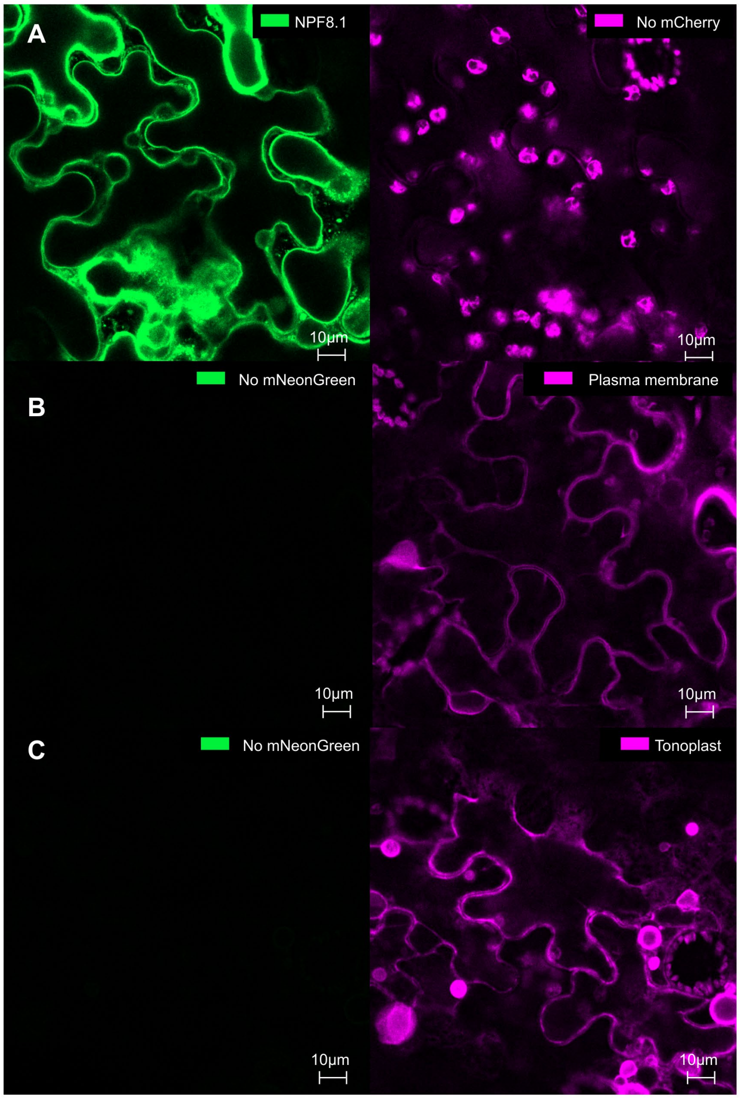

**Fig. S29. Single fluorophore controls for confocal imaging of localization markers with NPF8.1-mNeonGreen**

**(A)** Control images of NPF8.1-mNeonGreen in the absence of membrane marker show channel specificity **(B)** Control images of the plasma membrane marker PIP2A-mCherry in the absence of NPF8.1-mNeonGreen show channel specificity **(C)** Control images of the tonoplast marker γ-TIP-mCherry in the absence of NPF8.1-mNeonGreen show channel specificity.

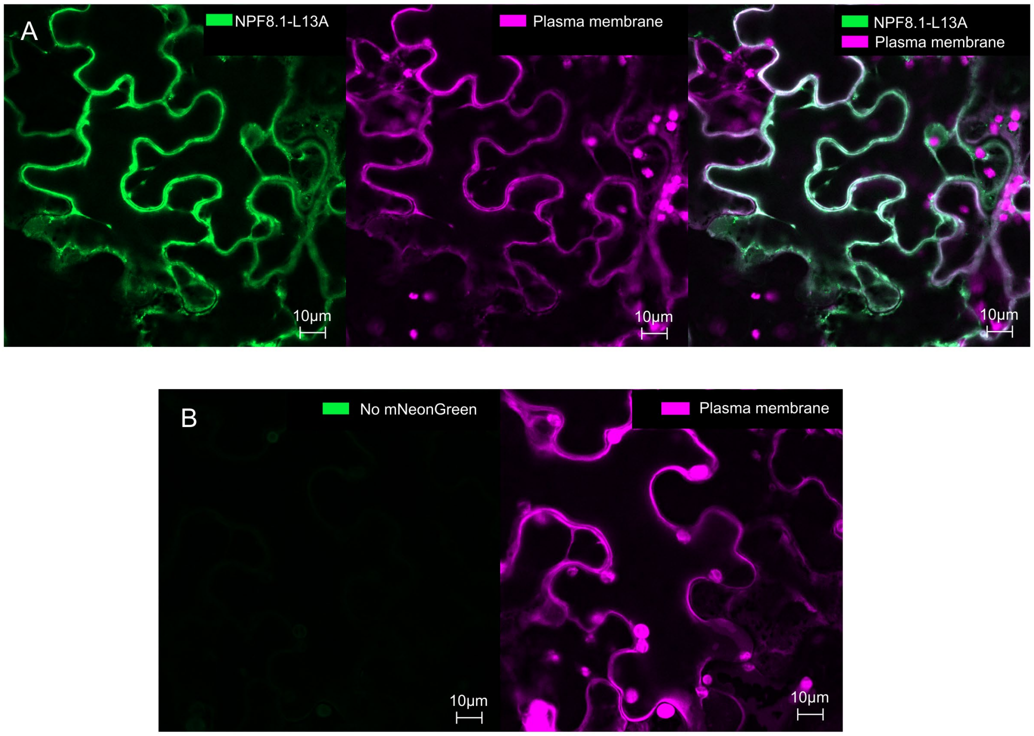

**Fig. S30. Confocal microscopy of NPF8.1-L13A-mNeonGreen with the plasma membrane marker**

**(A)** NPF8.1-L13A-mNeonGreen colocalizes with a plasma membrane marker PIP2A-mCherry. **(B)** Single fluorophore control images of the plasma membrane marker PIP2A-mCherry in absence of NPF8.1-L13A-mNeonGreen show channel specificity. Note the single NPF8.1-L13A-mNeonGreen image without PIP2A-mCherry is displayed in Fig. 3F.

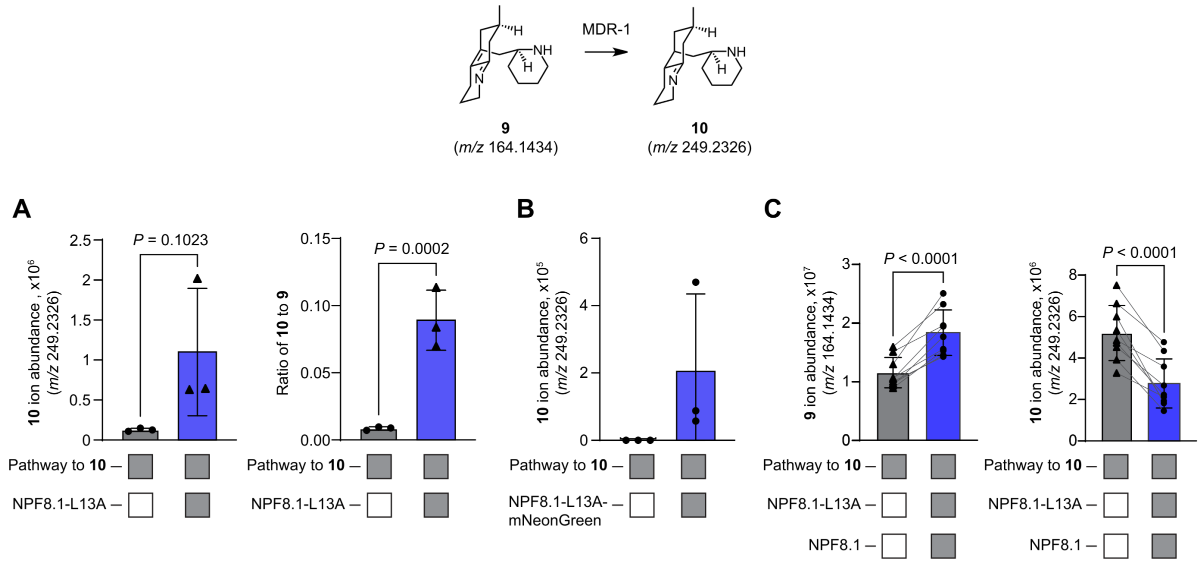

**Fig. S31. Characterization of NPF8.1-L13A via pathway reconstruction in *N. benthamiana*.** **(A)** Quantification of LC-MS data from transient expression of the pathway to **10** in *N. benthamiana* +/- NPF8.1-L13A shows that this mutant transporter still enables enriched production of **10** during pathway reconstitution. $P = 0.1023$ using unpaired Student's t-test (N=3) for total **10** abundance. $P = 0.0002$ using lognormal Student's t-test on the ratio of **10** to **9** on data from the same experiment. **(B)** Quantification of LC-MS data from transient expression of the pathway to **10** in *N. benthamiana* +/- NPF8.1-L13A-mNeonGreen shows the mutant transporter with a fluorophore fusion still enables enriched production of **10** during pathway reconstitution (quantification of trace from Fig. 3E) **(C)** Quantification of LC-MS data from transient expression of the pathway to **10** using either wildtype NPF8.1 or NPF8.1-L13A shows the mutant enables conversion of **9** to **10** less efficiently than the wildtype version. $P < 0.0001$ for both intermediates using a paired Student's t-test (N=9).

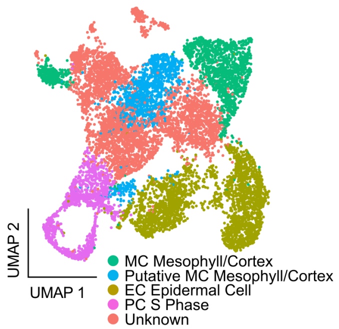

**Fig. S32. Cell type annotations of clusters from *D. obscurum* snRNA-seq data.** Using previously reported marker genes from *Lycopodium japonicum* shoot-apex tissue (*72*), nuclei clusters were assigned to known cell types. Shown here is the Uniform manifold approximation and projection (UMAP) of the cell type-annotated nuclei transcriptomes.

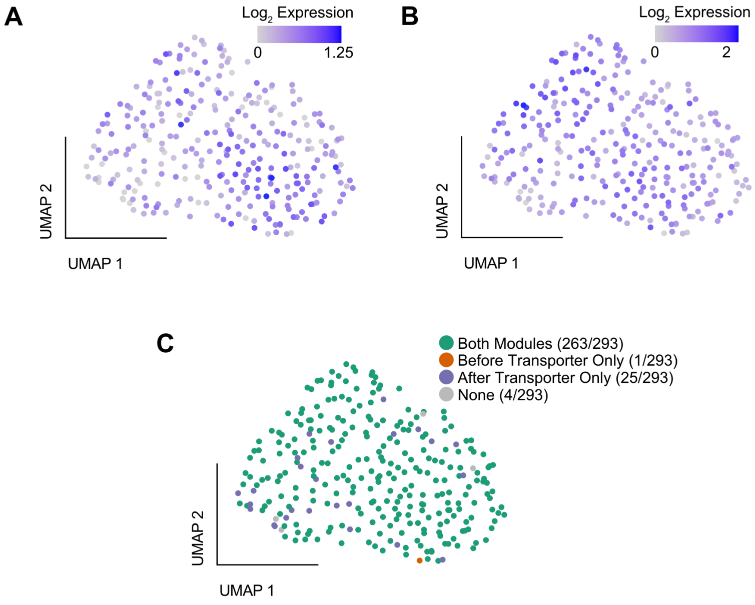

**Fig. S33. Expression analysis of pre- and post-transporter lycopodine (22) biosynthetic modules within cluster 9 from *D. obscurum* snRNA-seq data.**
**(A)** UMAP of 293 cells found in cluster 9 (see Fig 3G) colored by the mean expression of pre-transporter lycopodine (**22**) pathway genes (LDC, CAO, PIKS, CAL-3, SDR, ACT-1, CYP782C1, CAL-1, CAL-2). **(B)** UMAP of 293 cells found in cluster 9 colored by the mean expression of post-transporter lycopodine (**22**) pathway genes (NPF8.1, MDR-1, CYP7032A12, ACT-2, 2OGD-**4**, MDR-2, ABH-2). **(C)** Binary classification of individual nuclei based on co-expression of pre- and post-transporter modules. 89.8% of nuclei in cluster 9 are classified as expressing both modules with Fisher's exact test showing a significant positive association between two modules across individual nuclei (*P* = 0.000367, odds ratio = 40.86).

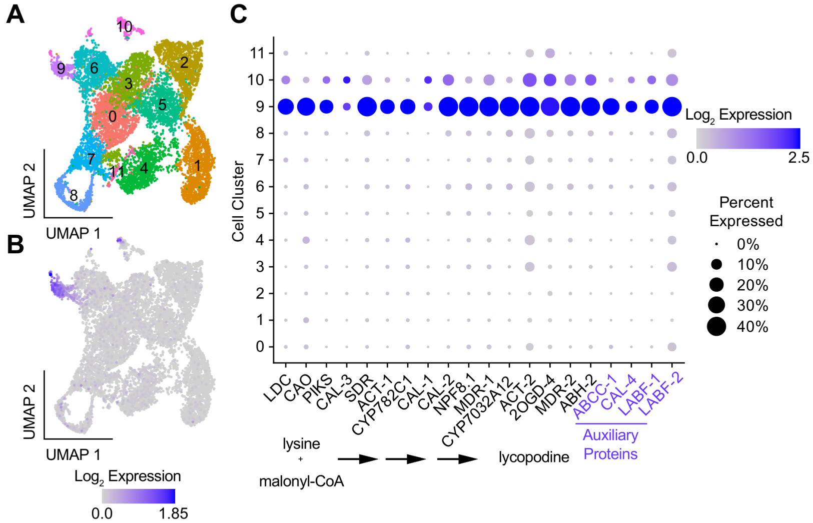

**Fig. S34. Cell type enrichment of auxiliary proteins in *D. obscurum* snRNA-seq data**
**(A)** UMAP of 10,679 single-nucleus transcriptomes colored by Seurat graph-based clusters (11 clusters; PCs 1-20; resolution = 0.5). **(B)** Full lycopodine (**22**) biosynthetic gene-set expression on UMAP. Color scale indicates the summed $\log_2$(average expression +1) of all 16 biosynthetic genes from panel (C). **(C)** Gene expression dot plot of lycopodine (**22**) biosynthetic genes across cell clusters detected in (A) ordered from upstream to downstream (left to right). Color scale shows the average scaled expression of each gene at each cell cluster determined based on Seurat clustering

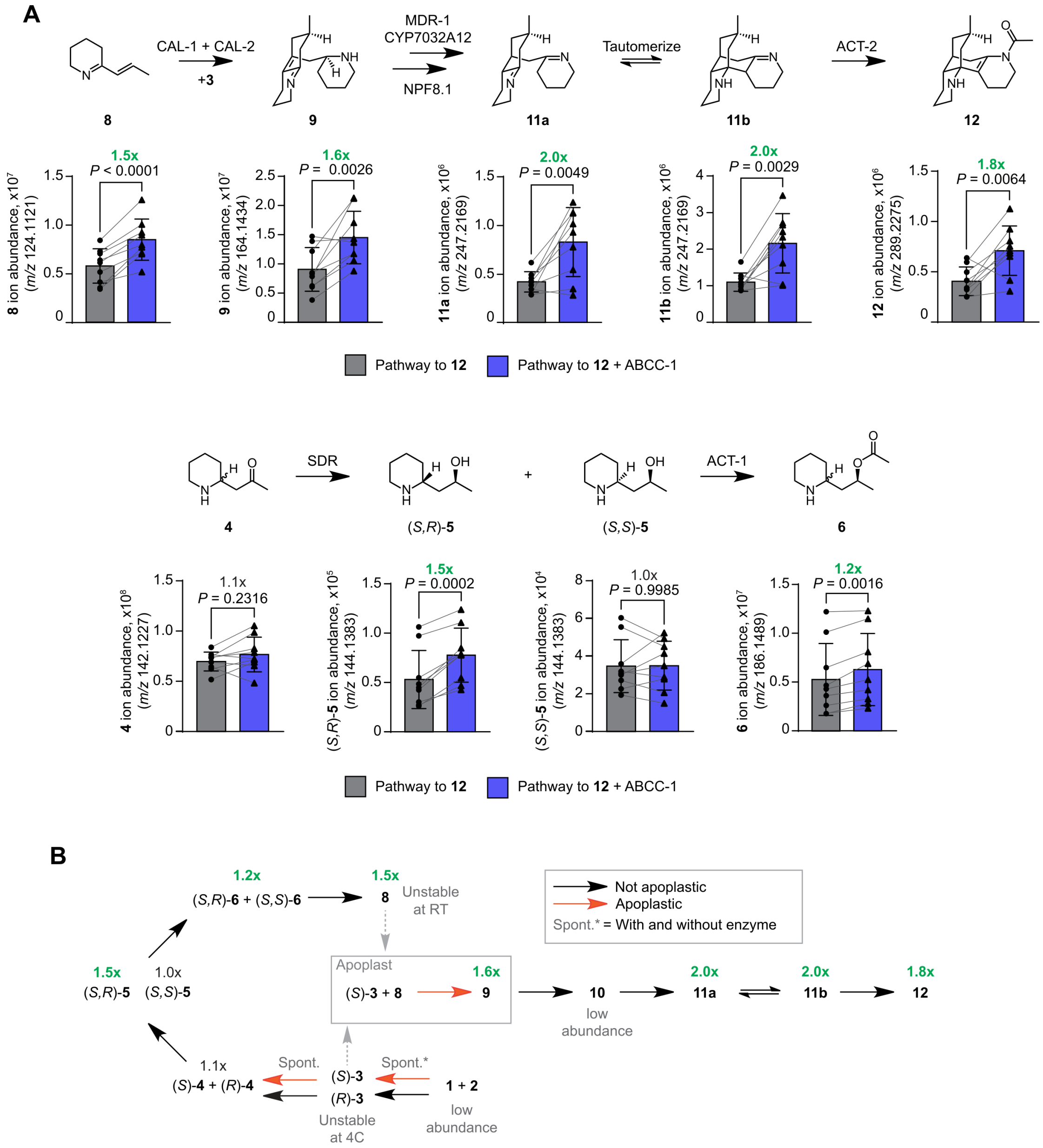

**Fig. S35. The effect of ABCC-1 on the flabellidine (12) pathway**.
**(A)** Quantification of pathway intermediates in *N. benthamiana* during flabellidine (**12**) pathway reconstruction in the absence (gray) or presence (blue) of ABCC-1. All P-values are the result of paired Student's t-tests (N=9). **(B)** Schematic of the effect of ABCC-1 on quantifiable intermediates from data shown in (A).

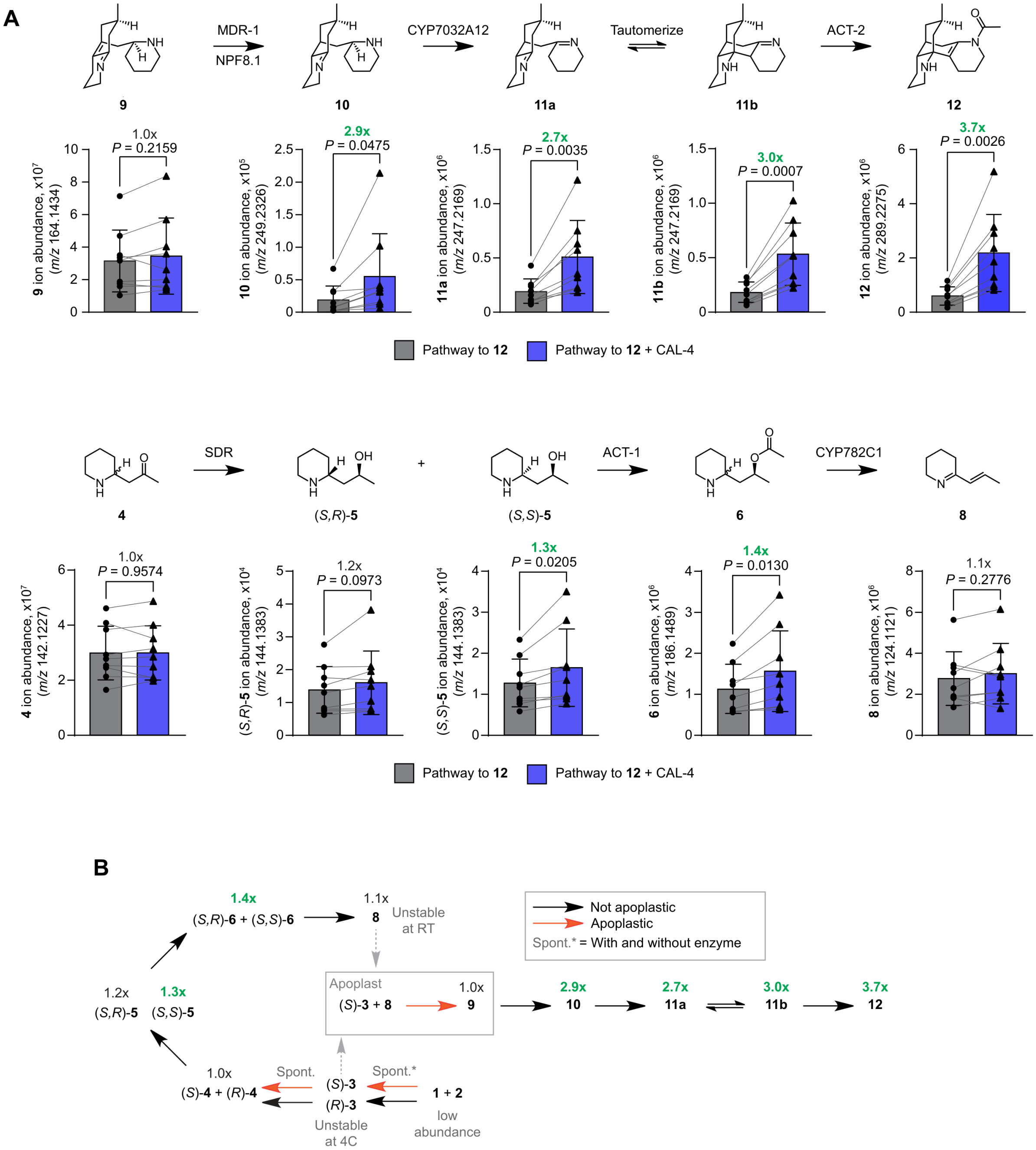

**Fig. S36. The effect of CAL-4 on the flabellidine (12) pathway**.
**(A)** Quantification of pathway intermediates in *N. benthamiana* during flabellidine (**12**) pathway reconstruction in the absence (gray) or presence (blue) of CAL-**4**. All P-values are the result of paired Student's t-tests (N=9). **(B)** Schematic of the effect of CAL-**4** on quantifiable intermediates from data shown in panel (A).

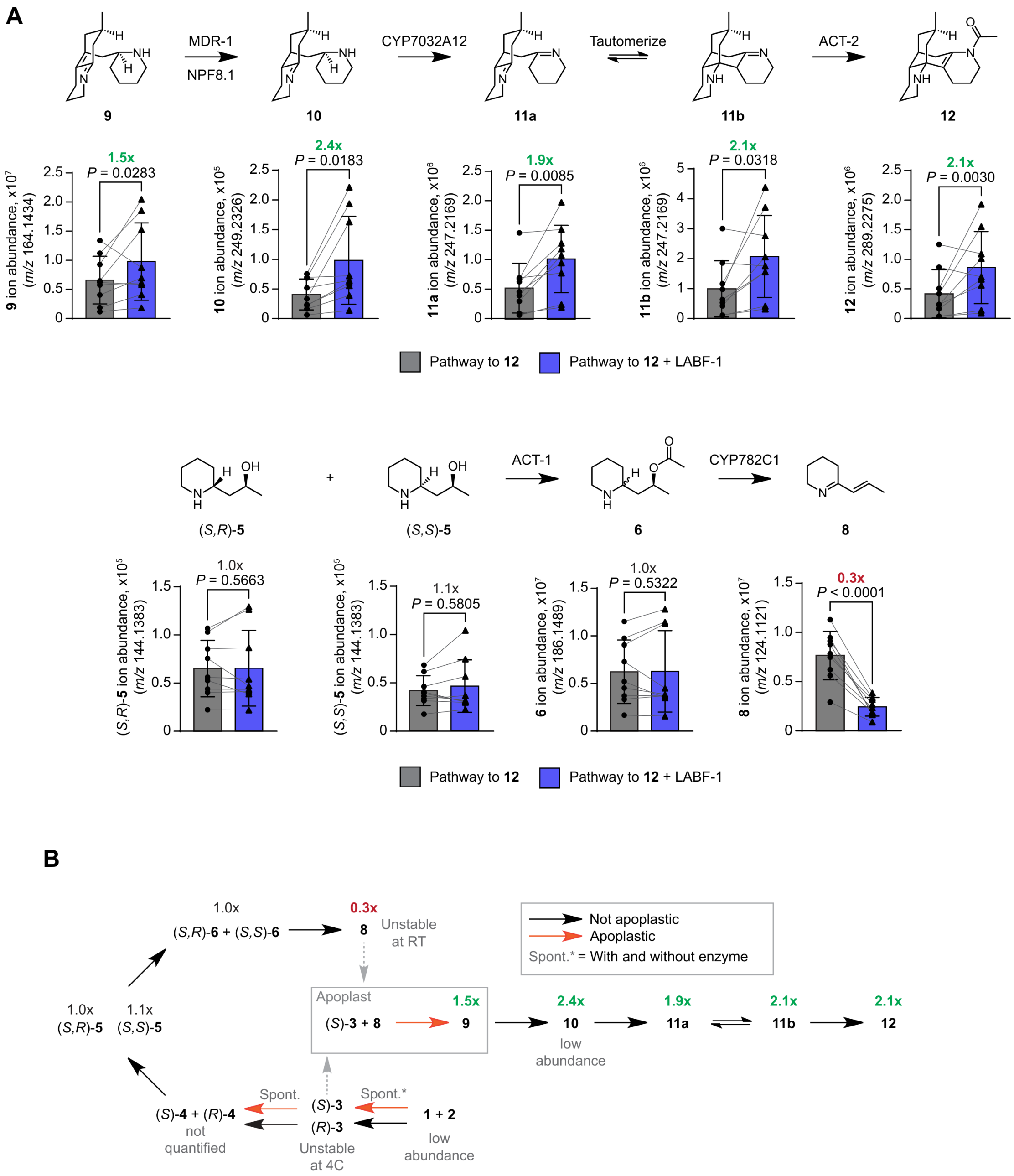

**Fig. S37. The effect of LABF-1 on the flabellidine (12) pathway**.
**(A)** Quantification of pathway intermediates in *N. benthamiana* during flabellidine (**12**) pathway reconstruction in the absence (gray) or presence (blue) of LABF-1. All P-values are the result of paired Student's t-tests (N=9). **(B)** Schematic of the effect of LABF-1 on quantifiable intermediates from data shown in panel (A).

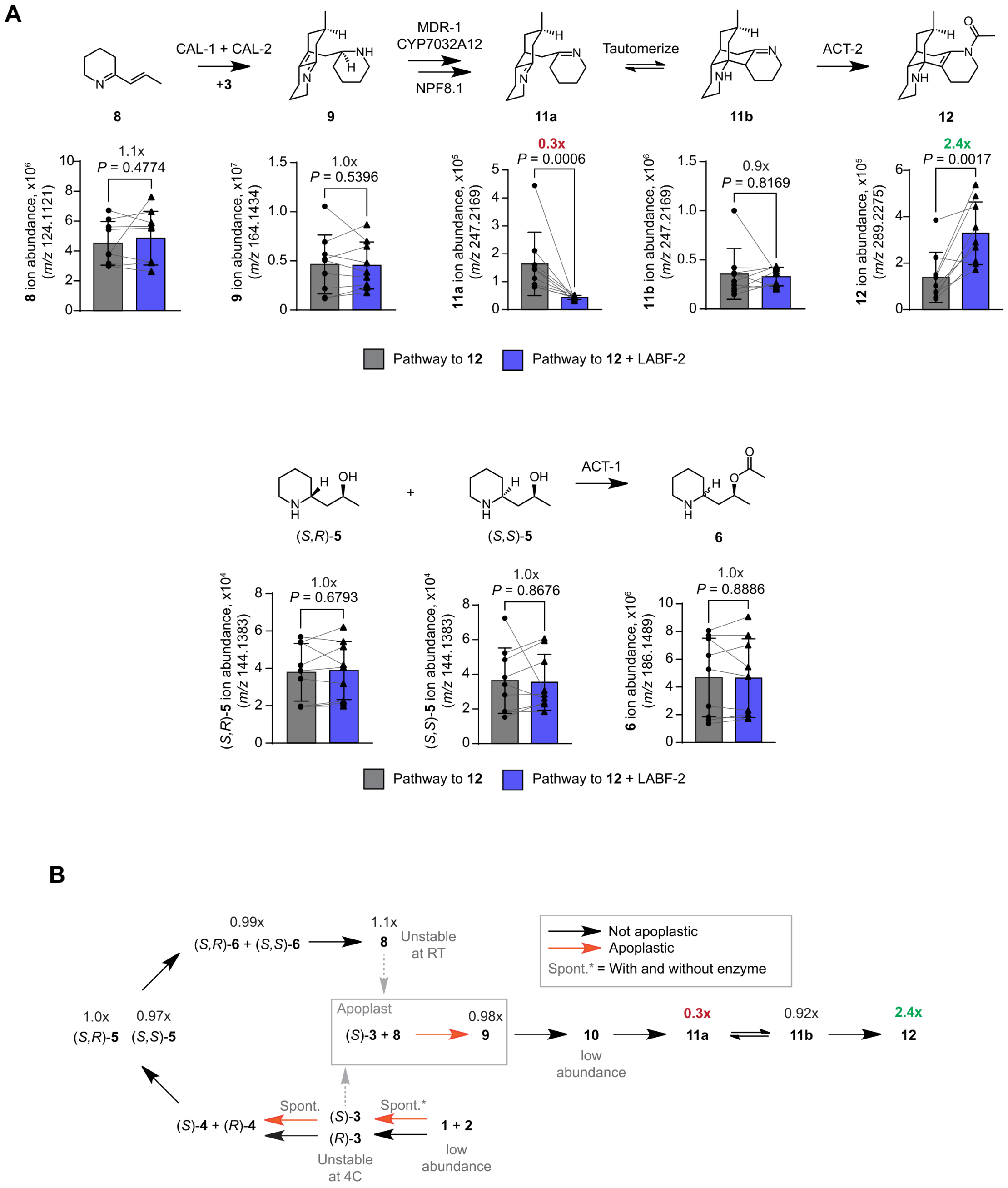

**Fig. S38. The effect of LABF-2 on the flabellidine (12) pathway**.
**(A)** Quantification of pathway intermediates in *N. benthamiana* during flabellidine (**12**) pathway reconstruction in the absence (gray) or presence (blue) of LABF-2. All P-values are the result of paired Student's t-tests (N=9). **(B)** Schematic of the effect of LABF-2 on quantifiable intermediates from data shown in panel (A).

#### Supplementary Methods

Listed below are LC-MS methods and parameters used for data acquisition and methods for HPLC and Flash chromatography used for small molecule purification.

**HPLC method parameters**

| | |
|---|---|
| **Instrument:** | Agilent 6546 LC-MS |
| **Method:** | C18 9 minute method |
| **Column:** | ZORBAX RRHD Eclipse Plus C18 (2.1 × 50 mm, 1.8 µm) |
| **Solvent A:** | water with 0.1% formic acid |
| **Solvent B:** | acetonitrile with 0.1% formic acid |
| **Flow rate:** | 0.6 mL/min |
| **Injection vol.:** | 1 µL |
| **Notes:** | Used for analysis of intermediates **12–20**, and **22** |

| Time (min) | A [%] | B [%] |
|---|---|---|
| 0.00 | 97.00 | 3.00 |
| 0.50 | 97.00 | 3.00 |
| 6.00 | 50.00 | 50.00 |
| 6.50 | 5.00 | 95.00 |
| 8.00 | 5.00 | 95.00 |
| 8.01 | 97.00 | 3.00 |
| 9.00 | 97.00 | 3.00 |

| | |
|---|---|
| **Instrument:** | Agilent 6546 LC-MS |
| **Method:** | HILIC 16 minute method |
| **Column:** | InfinityLab Poroshell 120 HILIC (2.1 × 150 mm, 1.9 µm) |
| **Solvent A:** | water with 10 mM ammonium formate and 0.1% formic acid |
| **Solvent B:** | 90% acetonitrile / 10% water with 10 mM ammonium formate and 0.1% formic acid |
| **Flow rate:** | 0.4 mL/min |
| **Injection vol.:** | 1 µL |
| **Notes:** | Used for analysis of intermediates **4**, **5**, **6**, **7**, **9**, **10**, **11**, **12**, **19**, **20**, **21**, **22** |

| Time (min) | A [%] | B [%] |
|---|---|---|
| 0.00 | 0.00 | 100.00 |
| 0.50 | 0.00 | 100.00 |
| 10.50 | 40.00 | 60.00 |
| 12.00 | 40.00 | 60.00 |
| 12.01 | 0.00 | 100.00 |
| 16.00 | 0.00 | 100.00 |

| | |
|---|---|
| **Instrument:** | Agilent 6546 LC-MS |
| **Method:** | HILIC-Z 14 minute method |
| **Column:** | Poroshell 120 HILIC-Z (2.1 × 100 mm, 1.9 µm) |
| **Solvent A:** | water with 10 mM ammonium formate and 0.1% formic acid |
| **Solvent B:** | 90% acetonitrile / 10% water with 10 mM ammonium formate and 0.1% formic acid |
| **Flow rate:** | 0.25 mL/min |
| **Injection vol.:** | 1 µL |
| **Notes:** | Used for analysis of intermediates **4**, **5**, **6**, **7**, **9**, **10**, **11**, **12** |

| Time (min) | A [%] | B [%] |
|---|---|---|
| 0.00 | 0.00 | 100.00 |
| 3.00 | 0.00 | 100.00 |
| 8.00 | 40.00 | 60.00 |
| 9.00 | 0.00 | 100.00 |
| 14.00 | 0.00 | 100.00 |

**Mass spectrometer parameters**

| Instrument: 6546 LC-MS | |
|---|---|
| **Source** | Dual AJS ESI |
| **Mode** | ESI positive, MS |
| **Data type** | Centroid |
| **Drying gas temp** | 325 °C |
| **Drying gas flow rate** | 10 L/min |
| **Nebulizer** | 35 psig |
| **Sheath gas temp** | 350 °C |
| **Sheath gas flow** | 12 L/min |
| **VCap** | 4000 V |
| **Fragmentor** | 135 V |
| **Skimmer** | 45 V |
| **OCT 1 RF Vpp** | 750 V |

| | |
|---|---|
| **Instrument:** | Agilent 1260 Infinity II |
| **Method:** | HILIC-Z 10 minute purification method |
| **Column:** | Poroshell 120 HILIC-Z (2.1 × 150 mm, 2.7 µm) |
| **Solvent A:** | water with 10 mM ammonium formate and 0.1% formic acid |
| **Solvent B:** | 95% acetonitrile with 10 mM ammonium formate and 0.1% formic acid |
| **Flow rate:** | 0.3 mL/min |
| **Injection vol.:** | 100 µL |
| **Temperature:** | 30 °C |
| **Notes:** | HILIC-Z HPLC method for purification of **9** |

| Time (min) | A [%] | B [%] |
|---|---|---|
| 0.00 | 20.00 | 80.00 |
| 1.00 | 20.00 | 80.00 |
| 6.00 | 40.00 | 60.00 |
| 8.00 | 40.00 | 60.00 |
| 8.01 | 20.00 | 80.00 |
| 10.00 | 20.00 | 80.00 |

| | |
|---|---|
| **Instrument:** | Agilent 1260 Infinity II |
| **Method:** | HILIC 10 minute purification method |
| **Column:** | Poroshell 120 HILIC (4.6 × 100 mm, 2.7 µm) |
| **Solvent A:** | water with 10 mM ammonium formate and 0.1% formic acid |
| **Solvent B:** | 95% acetonitrile with 10 mM ammonium formate and 0.1% formic acid |
| **Flow rate:** | 1.0 mL/min |
| **Injection vol.:** | 100 µL |
| **Temperature:** | 40 °C |
| **Notes:** | HPLC method for purification of **12** |

| Time (min) | A [%] | B [%] |
|---|---|---|
| 0.00 | 0.00 | 100.00 |
| 1.00 | 0.00 | 100.00 |
| 5.00 | 16.00 | 84.00 |
| 5.01 | 40.00 | 60.00 |
| 7.00 | 40.00 | 60.00 |
| 7.01 | 0.00 | 100.00 |
| 10.00 | 0.00 | 100.00 |

| | |
|---|---|
| **Instrument:** | Agilent 1260 Infinity II |
| **Method:** | EC-C18 11 minute purification method |
| **Column:** | Poroshell 120 EC-C18 (4.6 × 100 mm, 2.7 µm) |
| **Solvent A:** | water with 0.1% formic acid |
| **Solvent B:** | acetonitrile with 0.1% formic acid |
| **Flow rate:** | 1.0 mL/min |
| **Injection vol.:** | 100 µL |
| **Temperature:** | 40 °C |
| **Notes:** | EC-C18 HPLC method for purification of **20** |

| Time (min) | A [%] | B [%] |
|---|---|---|
| 0.00 | 97.00 | 3.00 |
| 0.50 | 97.00 | 3.00 |
| 7.00 | 70.00 | 30.00 |
| 7.01 | 5.00 | 95.00 |
| 9.00 | 5.00 | 95.00 |
| 9.01 | 97.00 | 3.00 |
| 11.00 | 97.00 | 3.00 |

| | |
|---|---|
| **Instrument:** | Biotage Selekt |
| **Method:** | Flash chromatography amino-column gradient |
| **Cartridge:** | Biotage® Sfär KP-Amino D Duo 50 µm, 11 g |
| **Column volume:** | 15 mL |
| **Flow rate:** | 12 mL/min |
| **Solvent A:** | hexanes |
| **Solvent B:** | ethyl acetate |
| **Notes:** | Normal-phase flash chromatography method |

| Column volume (CV) | A [%] | B [%] |
|---|---|---|
| 0 | 100 | 0 |
| 3 | 100 | 0 |
| 10 | 0 | 100 |
| 5 | 0 | 100 |

1466

**Supplementary Code**

This file contains all code used for data analysis and processing for this work. See the readme file for descriptions of each file. Certain code and related text documents were generated using the assistance of AI tools (e.g. ChatGPT, Codex).

**Supplementary Scheme**

Numbering and lettering scheme for atoms of Lycopodium alkaloids containing the lycopodane and lycodane scaffolds.

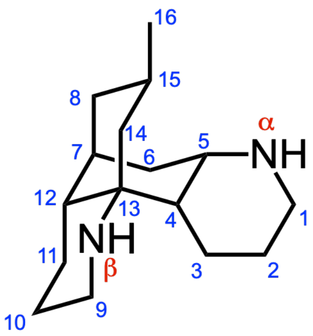

lycopodane scaffold

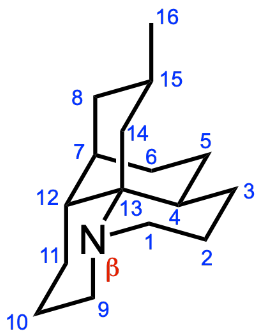

lycodane scaffold
